# Schizophrenia risk conferred by protein-coding *de novo* mutations

**DOI:** 10.1101/495036

**Authors:** Daniel P. Howrigan, Samuel A. Rose, Kaitlin E. Samocha, Menachem Fromer, Felecia Cerrato, Wei J. Chen, Claire Churchhouse, Kimberly Chambert, Sharon D. Chandler, Mark J. Daly, Ashley Dumont, Giulio Genovese, Hai-Gwo Hwu, Nan Laird, Jack A. Kosmicki, Jennifer L. Moran, Cheryl Roe, Tarjinder Singh, Shi-Heng Wang, Stephen V. Faraone, Stephen J. Glatt, Steven A. McCarroll, Ming Tsuang, Benjamin M. Neale

**Affiliations:** Analytic and Translational Genetics Unit, Massachusetts General Hospital, Boston, Massachusetts, USA; Broad Institute of MIT and Harvard, Cambridge, Massachusetts, USA; Icahn School of Medicine at Mount Sinai, New York, New York, USA; University of California, San Diego, California, USA; Harvard School of Public Health, Boston, Massachusetts, USA; National Taiwan University, Taiwan; SUNY Upstate Medical University, Syracuse, New York, USA; China Medical University, Taiwan; Harvard University, Cambridge, Massachusetts, USA

## Abstract

Protein-coding *de novo* mutations (DNMs) in the form of single nucleotide changes and short insertions/deletions are significant genetic risk factors for autism, intellectual disability, developmental delay, and epileptic encephalopathy. In contrast, the burden of DNMs has thus far only had a modest documented impact on schizophrenia (SCZ) risk. Here, we analyze whole-exome sequence from 1,695 SCZ affected parent-offspring trios from Taiwan along with DNMs from 1,077 published SCZ trios to better understand the contribution of coding DNMs to SCZ risk. Among 2,772 SCZ affected probands, the increased burden of DNMs is modest. Gene set analyses show that the modest increase in risk from DNMs in SCZ probands is concentrated in genes that are either highly brain expressed, under strong evolutionary constraint, and/or overlap with genes identified as DNM risk factors in other neurodevelopmental disorders. No single gene meets the criteria for genome-wide significance, but we identify 16 genes that are recurrently hit by a protein-truncating DNM, which is a 3.15-fold higher rate than mutation model expectation of 5.1 genes (permuted 95% CI=1-10 genes, permuted *p*=3e-5). Overall, DNMs explain only a small fraction of SCZ risk, and this risk is polygenic in nature suggesting that coding variation across many different genes will be a risk factor for SCZ in the population.

## Introduction

Schizophrenia (SCZ) is a severe psychiatric disorder that affects 0.7-1% of the general population, yet its etiology and pathophysiology is only beginning to be understood. The high heritability of SCZ suggests that inherited genetic factors make up a substantial proportion of the genetic risk, and recent advances from large-scale GWAS point to a broad polygenic network of brain expressed genes contributing to its pathophysiology (Schizophrenia Working Group of the Psychiatric Genomics 2014; Sekar, et al. 2016). Furthermore, genome-wide SNP-based heritability estimates suggest that ∼27% of genetic variation in SCZ is effectively tagged by variants with MAF > 1% in the population (Lee, et al. 2012).

Nevertheless, a large proportion of the heritability remains to be discovered or quantified. One potential source of undiscovered risk may come from rare, often deleterious, variants of more recent origin. The marked reduction in fecundity found in patients with SCZ suggests that natural selection may be removing the largest effect risk alleles from the population (Power, et al. 2013). Indeed, the strongest individual genetic risk factors discovered so far for SCZ are from large copy number variants (CNVs) that often occur *de novo* in affected offspring (Malhotra and Sebat 2012; Rees, et al. 2014; Marshall, et al. 2017). With large-scale whole-exome sequencing, we can discover *de novo* mutations (DNMs) in the protein-coding region of the genome at base-pair resolution, facilitating the discovery of individual SCZ risk genes.

DNMs discovered from parent-offspring trios have yielded novel genetic insights for an array of severe neurodevelopmental disorders, including intellectual disability (de Ligt, et al. 2012; Rauch, et al. 2012; Lelieveld, et al. 2016), autism spectrum disorders (Iossifov, et al. 2012; Neale, et al. 2012; O’Roak, et al. 2012; Sanders, et al. 2012; De Rubeis, et al. 2014; Iossifov, et al. 2014), epileptic encephalopathy (Epi, et al. 2013; Epi 2016), and developmental delay (Deciphering Developmental Disorders 2015, 2017). Along with an elevated burden of disruptive and damaging coding DNMs in these cohorts, specific biological pathways and individual genes have been robustly identified as *de novo* risk factors. For SCZ, an enrichment of damaging and disruptive DNMs has been modest by comparison. Recent published reports using either parent-child trio or case-control studies have shown enrichment in specific brain-expressed gene sets (e.g. ARC/NMDAR protein complexes, FMRP interactors) and overlap with genes implicated in the neurodevelopmental disorders listed above (Fromer, et al. 2014; Genovese, et al. 2016), however only a single gene, *SETD1A*, has been robustly identified as a *de novo* risk factor for SCZ (Takata, et al. 2014; Singh, et al. 2016). These findings suggest that highly penetrant SCZ risk genes are unlikely to exist, and larger samples are needed to robustly identify genes with more modest contributions to SCZ risk.

### SCZ Trio Cohort from Taiwan

Here, we report whole-exome sequencing results from 1,695 complete parent-proband trios (1,033 male and 662 female probands). Families were recruited from mental hospitals, community care centers and primary care clinics across the island of Taiwan. To our knowledge, this is the largest trio exome sequence study to date in SCZ. All trios consist of a SCZ affected proband and unaffected parents. Trios consist predominantly of sporadic case probands (91%), with the remainder reporting a history of mental illness in the family. All samples were exome-sequenced and analyzed at the Broad Institute in Cambridge, MA, USA, with a target of 20x depth in at least 80% of the exome target. All reported DNMs were either validated or showed strong confidence in the initial call (**supplementary section 4**). Exome sequencing was performed in three distinct waves, consisting of 575, 532, and 588 trios, respectively.

Among measured covariates that could affect mutation rates, parental age was the most significant predictor of increased DNM burden (*p*=3.6e-7). None of the other tested study covariates tested significantly predicted DNM rates, but there are suggestive indications that females carry a modestly higher burden of DNMs (*p*=0.06), while probands with a positive family history of mental illness have a slightly lower burden of DNMs (*p*=0.14; **supplementary section 9**). Of note, the larger capture target used in the 3^rd^ wave of sequencing did lead to higher overall DNM rates, but we accounted for this discrepancy by excluding DNMs outside of overlapping exome capture intervals when evaluating exome-wide DNM burden. The strong correlation of paternal and maternal age (*r*=0.49) makes it difficult to fully delineate parent-of-origin effects, however when we fit both paternal age and maternal age in the regression model, paternal age (*p*=3e-6) associated with increased DNM rates over and above maternal age (*p*=0.38). Further follow up on parental age effects revealed a modest quadratic effect of increased maternal age (*p*=0.01) associated with increased DNM rate, something not seen in paternal age (*p*=0.6). These results show a clear association of older paternal age with increasing DNM rates and suggest that older maternal age may also lead to increased DNM rates in a possibly non-linear fashion, albeit attenuated relative to paternal age.

### Exome-wide rates of *de novo* mutation

Using a Poisson distributed model specification, exome-wide DNM rates in the Taiwanese cohort were compared against 846 trios from previous published exome DNM studies in SCZ (Girard, et al. 2011; Xu, et al. 2011; Xu, et al. 2012; Gulsuner, et al. 2013; Fromer, et al. 2014; Guipponi, et al. 2014; McCarthy, et al. 2014) (**see supplementary section 10 for study inclusion criteria**). After restricting our search to overlapping exome capture intervals, we find no significant difference in the overall DNM rate of the three Taiwanese sequencing waves when compared to published SCZ trios (rate-ratios=1.01, 0.97, and 1.02, respectively; all *p* > 0.05). We combined the Taiwanese cohort with published SCZ trios (2,541 trios), and compared DNM rates against a DNM expectation model restricted to 17,925 well-covered genes (Samocha, et al. 2014; Lek, et al. 2016)(**supplementary section 8**) as well as 2,216 published unaffected siblings and control trios (collectively termed ‘controls’; **Fig. 1a**). We find the overall DNM rate in SCZ probands only slightly above the DNM model expectation (rate-ratio=1.02, *p*=0.43), but significantly enriched over controls (rate-ratio=1.08, *p*=9e-3). When we partition DNMs by coding annotation, we see a significantly lower rate of synonymous DNMs in SCZ probands relative to the DNM model (rate-ratio=0.87, *p*=8e-4), however this synonymous rate is consistent with the synonymous DNM rate observed in controls (rate-ratio=1.05, *p*=0.44), suggesting that DNM calling is not biased among SCZ probands relative to other exome sequenced cohorts, and that the DNM model is a conservative expectation exome-wide. Among protein-truncating variants (PTV), we find evidence of enrichment relative to DNM model expectations (rate-ratio=1.18, *p*=9e-3), albeit not significantly above controls (rate-ratio=1.11, *p*=0.22). For missense variants, we see a modest enrichment relative to both the DNM model (rate-ratio=1.06, *p*=0.03) and when compared to controls (rate-ratio=1.09, *p*=0.03).

**Figure 1:**
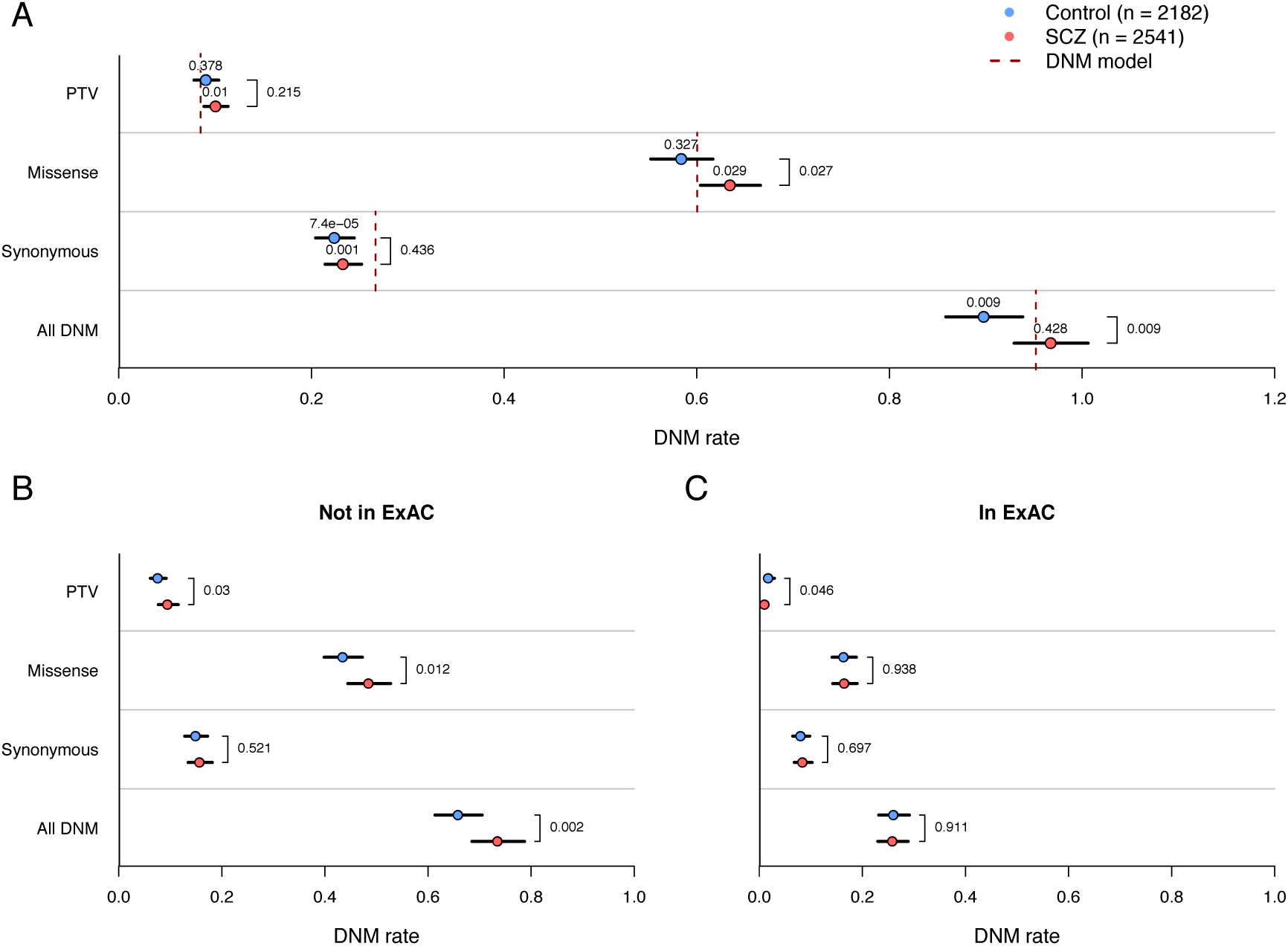
Proband DNM rates and significance among SCZ and control probands. **Figure 1A)** Exome-wide DNM rates split by primary annotation, where “All DNM” encompasses synonymous, PTV, and missense DNMs. Poisson rate *p*-values compared to the DNM model expectation (dotted lines) are directly above the rate estimate in cases and controls, and two-sample, two-sided Poisson exact test *p*-values between cases and controls are listed to the right of the rate estimates. DNM rates split by the absence (**Figure 1B**) or presence (**Figure 1C**) of an extant allele at the same position in the 45k non-psychiatric ExAC cohort, with two-sample, two-sided Poisson exact test *p*-values between cases and controls to the right of the rate estimates.

### SCZ DNMs are more likely to be annotated as deleterious

To see if SCZ DNMs were enriched for pathogenic alleles, we filtered on allele frequency from the Exome Aggregation Consortium (ExAC v0.3) and predicted functional consequence. First, we filtered out DNMs at variable sites in the non-psychiatric ExAC cohort (∼45k individuals; (Lek, et al. 2016), inferring that the presence of the same allele in ostensibly healthy individuals lowers the likelihood of being a pathogenic variant (**Fig. 1b,c**). Similar to DNMs found in probands with autism spectrum disorder (Kosmicki, et al. 2017), we find that 26% of DNMs found in SCZ probands are observed in the non-psychiatric ExAC cohort. When we filter out these alleles in both SCZ probands and controls, we see an increase in DNM enrichment for both PTV (from rate-ratio=1.11, *p*=0.21 to rate-ratio=1.25, *p*=0.03) and missense variants (from rate-ratio=1.09, *p*=0.03 to rate-ratio=1.12, *p*=0.01), but not in synonymous variants (from rate-ratio=1.05, *p*=0.52 to rate-ratio=1.05, *p*=0.7).

Second, we examined a variety of functional annotations that could further delineate the likely pathogenic alleles beyond the primary coding consequence. Previous SCZ case-control exome studies found that missense variants predicted as damaging from multiple prediction algorithms were associated with SCZ risk (Genovese, et al. 2016). All but one of the twelve missense prediction algorithms tested increased the missense DNM enrichment among SCZ probands relative to controls after filtering out predicted non-damaging missense variants (**supplementary section 11**). Outside of missense prediction algorithms, a recent analysis found that synonymous variants near exon splice junctions and within brain-derived DNAse hypersensitivity site (DHS) peaks were enriched in both published autism and SCZ DNMs (Takata, et al. 2016). When we replicate the analysis in the Taiwanese cohort (1,695 trios) and an independent set of controls (1,485 trios), we do not find a significant enrichment in synonymous variants within 30bp of the splice site (fold-enrichment=1.14, one-tailed *p*=0.26) and within cerebrum-frontal cortex DHS peaks (fold-enrichment=1.26, one-tailed *p*=0.09). Further analysis, however, reveals that a broader definition of being near a splice site (60 bp rather than 30 bp) shows more consistency in both samples and significant enrichment in the combined SCZ cohort (fold-enrichment=1.29, *p*=5e-4; **supplementary section 12**).

### DNM burden relative to other neurodevelopmental disorders

When we examine the combined cohort of 2772 SCZ probands within the larger context of mental illness, the enrichment signal is markedly reduced relative to probands diagnosed with early-onset neurodevelopmental disorders among both PTVs exome-wide (**Fig. 2a**) and missense DNMs in evolutionarily constrained genes (i.e. missense constrained (Samocha, et al. 2014), pLI > 0.99 (Lek, et al. 2016), or RVIS intolerant (Petrovski, et al. 2013) genes; **Fig. 2b**) – two categories where there is a significant enrichment in DNM burden across all disorders analyzed. This comparison clearly indicates that the contribution of DNMs towards a SCZ diagnosis accounts for a much smaller fraction of samples than earlier onset neurodevelopmental disorders.

**Figure 2:**
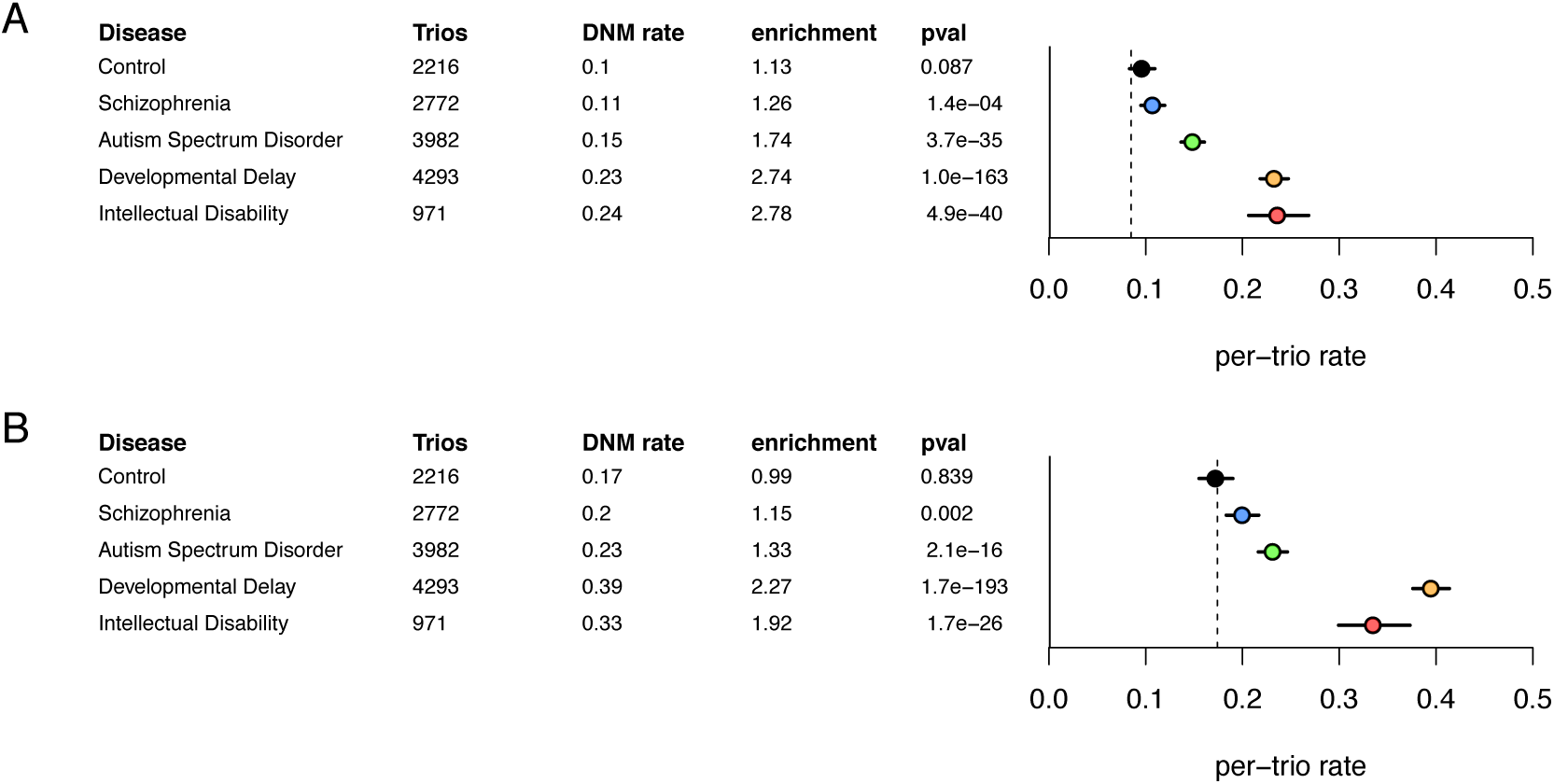
DNM burden for exome-sequenced trios with various mental disorders. **(Figure 2A)** Exome-wide PTV rate and enrichment relative to DNM model expectations. **(Figure 2B)** Missense rate among evolutionarily constrained genes (defined here as the union of high pLI, missense constraint, and RVIS intolerant gene sets) and enrichment relative to DNM model expectations.

### Enrichment in gene sets implicated by multiple constraint metrics and in multiple neurodevelopmental disorders

Along with DNM burden, we examined enrichment in specific gene sets after conditioning on the overall DNM rates (**supplementary section 14**) to understand the pattern of genetic risk conferred by DNMs. Among gene sets defined from evolutionarily constrained metrics defined above (**Fig. 3a**) and coding DNM studies of intellectual disability, developmental delay, and autism spectrum disorder (**Fig. 3b**), we find DNMs in SCZ probands are significantly enriched for genes repeatedly identified in multiple metrics and diseases. Among all DNMs, enrichment is biased towards genes implicated in all three measures of evolutionary constraint (350 genes), and among genes with recurrent PTVs identified in multiple neurodevelopmental disorders (32 genes). These results highlight the notion that despite the lower overall DNM burden in SCZ relative to other neurodevelopmental disorders, the disruption of some of the most critical genes in brain development confers risk towards a variety of mental disorders, including SCZ.

**Figure 3:**
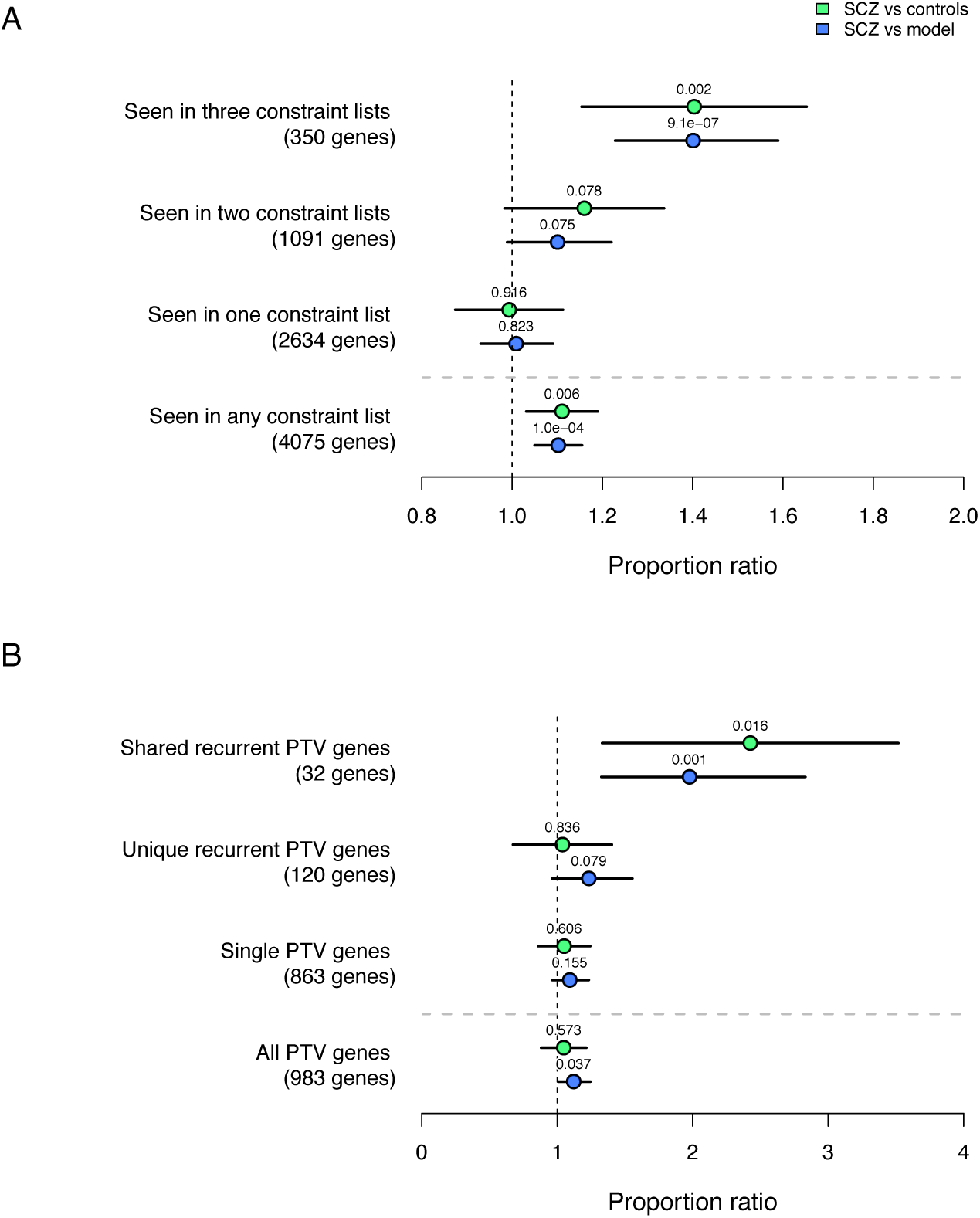
Partitioning gene set enrichment among evolutionarily constrained and neurodevelopmental disorder gene sets. **Figure 3A:** Gene set enrichment in three evolutionary constraint metrics (missense constraint, pLI, and RVIS), with partitioning among how often genes are present in each set. Proportion enrichment is tested among all DNMs in SCZ probands relative to controls (green) and the DNM model (blue). **Figure 3B:** Gene set enrichment for genes with observed DNM PTVs in intellectual disability, developmental delay, and autism spectrum disorder probands. Shared recurrent PTV genes have identified 2 or more PTVs in more than one disorder. Unique recurrent PTV genes have identified 2 or more PTVs in only one disorder. Single PTV genes are not recurrent within any disorder, although may be shared across disorders. Proportion enrichment is tested among all DNMs in SCZ probands relative to controls (green) and the DNM model (blue).

### Modest support for genes implicated in previous SCZ analyses

Among 461 genes implicated in the PGC GWAS (Schizophrenia Working Group of the Psychiatric Genomics 2014) and rare CNV (Marshall, et al. 2017) studies of SCZ, we see a non-significant enrichment relative to the DNM model (all DNM fold-enrichment=1.2, *p*=0.07; PTV fold-enrichment=1.55, *p*=0.11) and compared to controls (all DNM fold-enrichment=1.25, *p*=0.16; PTV fold-enrichment=1.28, *p*=0.59). Moreover, no single comparison reached experiment-wide significance, with the most significant enrichment coming in genes overlapping SCZ associated CNVs (151 genes tested, case/control all DNM fold-enrichment=1.98, *p*=0.05). Previously published SCZ trio cohorts reported several gene sets with significant association, namely prenatally biased genes (Xu, et al. 2012), chromatin modifiers (McCarthy, et al. 2014; Takata, et al. 2014), and the ARC/NMDAR subunits (Fromer, et al. 2014). None of these gene sets reached experiment-wide significance in the current analysis, with the most significant gene set coming from PTVs in proteins that interact with the ARC subunit complex (28 genes tested, PTV fold-enrichment=7.9, *p* = 2e-3), where an additional two genes, *IQSEC1* (IQ Motif and Sec7 Domain 1) and *ATP1A1* (ATPase Na+/K+ Transporting Subunit Alpha 1), are hit by PTV DNMs in the Taiwanese cohort. Notably, there is also a PTV hit in *ARC* in the Taiwanese cohort (which is by default not included in the ARC subunit gene set).

### Enrichment in highly brain expressed and constrained genes

Many biologically plausible gene sets have been previously implicated to refine the polygenic basis of SCZ risk towards more biologically tractable components. Among 85 candidate gene sets analyzed (**supplementary section 14**), BrainSpan high brain expression (8928 genes tested) and GTEx brain enriched (6214 genes tested; see (Ganna, et al. 2016)) represented two of the four gene sets that surpassed multiple-testing correction in both the DNM model and against controls (the other two gene sets being missense constraint and RVIS intolerant). Both gene sets represent gene expression across the entirety of human brain tissues and cell types measured from *post-mortem* brain tissue (http://www.brainspan.org/ and https://www.gtexportal.org/home/), and reinforce the notion that the genetic risk for SCZ, while brain-specific, is polygenic across the allele frequency spectrum.

### DNM enrichment in potentially synaptic components of the brain

In an exome case-control study of SCZ among Swedish participants, a variety of gene sets defined as mRNA targets of regulatory proteins (FMRP, RBFOX, CELF4) highly active in the synapse were enriched for damaging rare variation in SCZ cases (Purcell, et al. 2014; Genovese, et al. 2016). When we compare against the DNM model, we see a consistent pattern of enrichment in SCZ probands, with FMRP interactors and RBFOX1/3 splicing targets surpassing multiple-testing correction (*p* < 8e-4). Following the strategy presented in (Genovese, et al. 2016), we stratified both neuronal cell type expression and high brain expression gene sets by potentially synaptic genes (FMRP, RBFOX2, CELF4, and SynaptomeDB gene sets) identified from these mRNA targets, finding that the pattern of enrichment among neuronally expressed genes is driven primarily by synaptically localized genes (**Fig. 4**). Of note, the signal is not significantly enriched when comparing against controls, owing in part to the statistical power differences between the model and case-control tests (**supplementary section 14**). These findings provide independent support to the enrichment seen among Swedish SCZ case-control exomes and indicate that these gene sets are tapping into the elements of synaptic biology that, when perturbed, increase the risk for SCZ.

**Figure 4:**
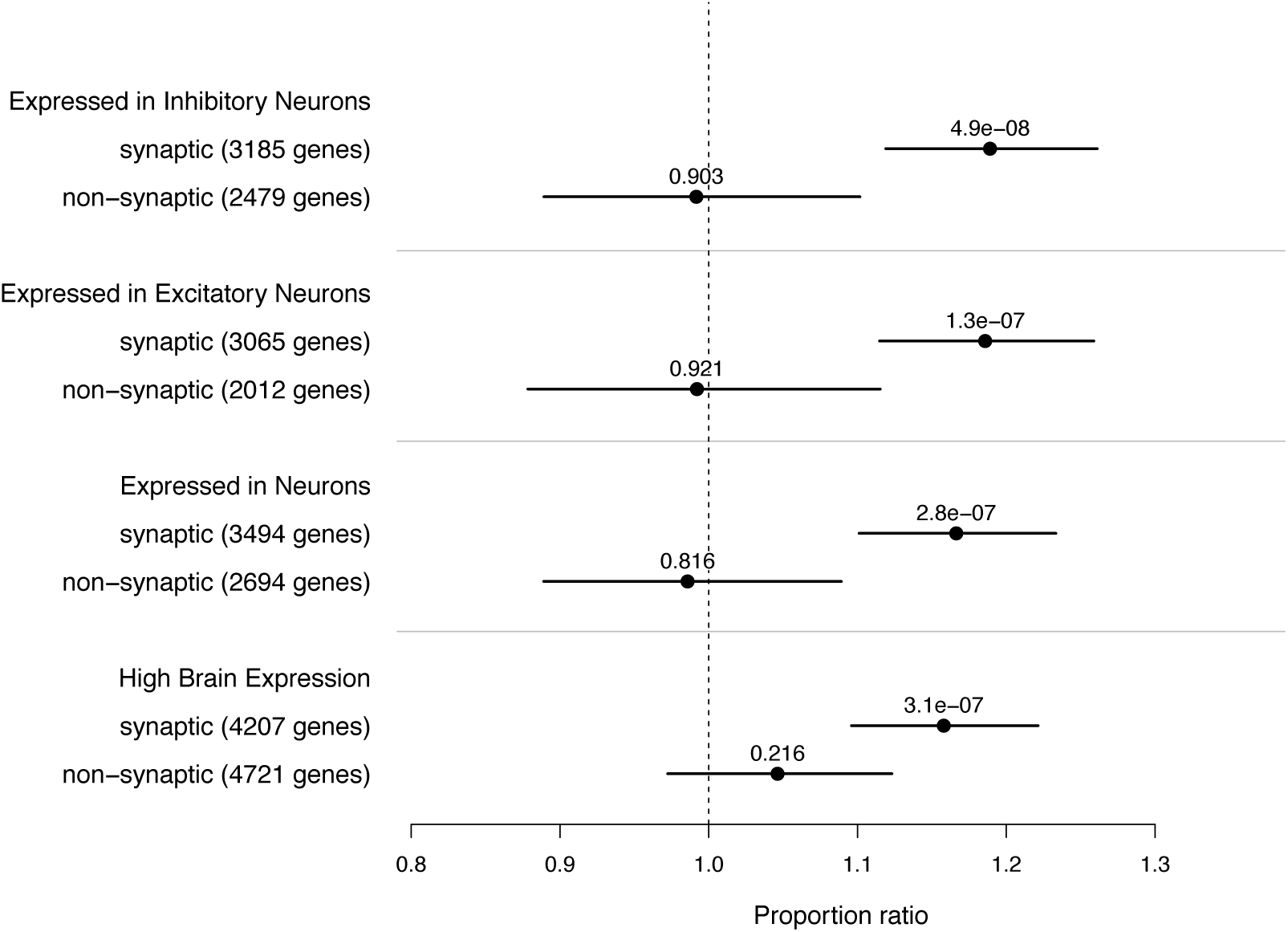
Partitioning of the genes expressed in neuronal cell types by their interaction with mRNAs highly active in the synapse (e.g. FMRP, RBFOX, CELF4), with interacting genes listed as “potentially synaptic”. Also included are the results from BrainSpan highly expressed genes (bottom comparison). Proportion enrichment is tested against the mutation model.

### GO and SynaptomeDB databases implicate neurotransmitter secretion and chromatin organization

Outside of candidate gene sets, we ran an unbiased gene set scan using annotations from the Gene Ontology (GO) and SynaptomeDB databases. We restricted our analysis to only gene sets with at least 50 genes and analyzed a total of 911 gene sets (**supplementary section 15**). We ran the same permutation procedure as the candidate gene set analysis to estimate our multiple testing correction cutoff at a 5% alpha level (cutoff *p*=6e-5 for the DNM model test and cutoff *p*=2e-4 for the case/control test). No single gene set surpassed multiple testing correction in either comparison; the most significant gene set among all DNMs overlapping genes involved the neurotransmitter secretion biological process (GO:0007269, 64 genes tested, fold-enrichment=2.12, *p*=4e-4). Among PTVs, the most significant enrichment overlaps genes involved in chromatin organization biological process (GO:0006325, 207 genes tested, fold-enrichment=2.81, *p*=4e-4;). When comparing against controls, the most significant gene set among all DNMs overlapping genes is the biosynthetic process in Synaptome DB (73 genes tested, fold-enrichment=15.8, *p* = 3e-4).

### Single gene association and gene recurrence rates

To see if any single gene was a putative risk factor for SCZ, we tested for per-gene enrichment of DNMs in the combined SCZ cohort against the gene level mutation expectation (**supplementary section 16**). Exome-wide significance threshold was set at *p* = 8.7e-7 to correct for multiple testing, and no gene surpassed exome-wide significance, with our lowest *p*-value across all three tests being *p*=7.7e-6.

Among PTVs, the most significant gene is SET Domain Containing 1A *(SETD1A*), with 3 PTVs observed in two previously published SCZ trio cohorts, ((Guipponi, et al. 2014; Takata, et al. 2014), pLI=1, *p*=7.7e-6). *SETD1A* has since been identified as an exome-wide significant gene association in combined trio and case/control exome sequencing of SCZ, whereby follow-up of patients carrying a PTV in *SETD1A* often presented with an associated neurodevelopmental disorder (Singh, et al. 2016).

### Enrichment in genes with recurrent PTVs

While no single gene association surpasses exome-wide correction, many genes are “recurrently” hit (i.e. more than one coding DNM hits the gene) among SCZ probands. The rate of recurrently hit genes can indicate how likely such genes are SCZ risk factors. To test observed rates of gene recurrence against the mutation expectation, we used bootstrap re-sampling to simulate an equivalent count of observed DNMs hitting genes, with the probability of hitting a gene being its mutation expectation (**supplementary section 17**). We also tested control DNMs to ensure that any significant results were not the result of mis-specification in the DNM model. We observe sixteen genes with recurrent PTVs (fold-enrichment=3.15, *p*=3e-5), a result significantly enriched above mutation model expectation of 5.1 genes recurrently hit by PTV DNMs (**Table 1**). When we perform the same analysis in controls, we do not find a significant enrichment in genes with recurrently hit PTVs (fold-enrichment=1.51, *p*=0.27). While we see modest enrichment in genes recurrently hit by missense and/or synonymous DNMs, the enrichment is similar in controls, and suggests deviations of the DNM model to the subtle effects of exome coverage and QC parameters. Further dissection of DNM recurrence within gene sets also indicates that highly expressed brain genes, particularly those not considered as evolutionarily constrained, are enriched for recurrent hits (**supplementary section 17**).

**Table 1:** **Gene recurrence rates in SCZ probands and controls**

| | DNM | Observed recurrent genes | Expected recurrent genes | Fold-enrichment | Empirical $p$-value |
| --- | --- | --- | --- | --- | --- |
| *2772 SCZ probands* |  |  |  |  |  |
| All DNM | 2694 | 314 | 286.2 | 1.10 | 0.02 |
| PTV | 296 | 16 | 5.1 | 3.15 | 3e-5 |
| Missense | 1763 | 150 | 136.6 | 1.10 | 0.09 |
| Synonymous | 635 | 29 | 21.2 | 1.37 | 0.05 |
| *2216 controls* |  |  |  |  |  |
| All DNM | 2044 | 195 | 177.4 | 1.10 | 0.06 |
| PTV | 212 | 4 | 2.7 | 1.51 | 0.27 |
| Missense | 1320 | 89 | 81.2 | 1.10 | 0.17 |
| Synonymous | 512 | 18 | 14.1 | 1.28 | 0.16 |

To further assess the contribution of recurrently hit PTV genes in SCZ probands, we examined the contribution of rare transmitted variation within these genes (**Table 2**). While the lack of a SCZ diagnosis in the parents is expected to lower the contribution that rare inherited variants have in proband risk, the polygenic nature of SCZ risk predicts that undiagnosed individuals carry SCZ risk alleles. Using parent-proband transmission counts in the full Taiwanese cohort (1695 trios), TDT among 15 recurrently hit PTV genes (excluding *TTN*) shows a 3.25-fold enriched ratio of transmitted ultra-rare PTVs to non-transmitted ultra-rare PTVs (*p*=0.02, **supplementary section 18**). The contribution of transmitted PTVs comes largely from two genes, the Trio Rho Guanine Nucleotide Exchange Factor gene (*TRIO*; 4 transmitted to 0 non-transmitted), and the Dynein Axonemal Heavy Chain 9 gene (*DNAH9*, 6 transmitted to 1 non-transmitted). While we do not observe any significant enrichment in ultra-rare damaging missense variants (defined here as ‘probably damaging’ in PolyPhen2, ‘deleterious’ in SIFT, and not seen in ExAC) among these 15 recurrently hit PTV genes (OR=1.09, *p*=0.64), *TRIO* shows a suggestive over-transmission among ultra-rare and predicted damaging missense variants (11 transmitted to 2 non-transmitted, *p*=0.01).

**Table 2:** **Genes recurrently hit by PTV DNMs**

| Gene symbol | Coding bases (canonical) | pLI | PTV DNM Poisson $p$-value | PTV DNM | Missense DNM | PTV transmitted | PTV non-transmitted | Damaging Missense transmitted | Damaging Missense non-transmitted |
| --- | --- | --- | --- | --- | --- | --- | --- | --- | --- |
| *SETD1A* | 5124 | 1 | 7.77E-06 | 3 | 0 | 0 | 0 | 1 | 4 |
| *GALNT9* | 714 | 0.56 | 1.27E-05 | 2 | 0 | 0 | 0 | 2 | 0 |
| *TAF13* | 375 | 0.08 | 2.43E-05 | 2 | 0 | 0 | 0 | 0 | 0 |
| *HENMT1* | 1182 | 0 | 3.61E-05 | 2 | 0 | 1 | 0 | 0 | 0 |
| *SV2B* | 2052 | 0.34 | 1.50E-04 | 2 | 0 | 1 | 0 | 1 | 2 |
| *NRXN3* | 3186 | 1 | 2.12E-04 | 2 | 0 | 0 | 0 | 6 | 6 |
| *SMARCC2* | 3645 | 1 | 6.03E-04 | 2 | 0 | 0 | 0 | 0 | 1 |
| *RB1CC1* | 4785 | 1 | 8.32E-04 | 2 | 0 | 0 | 0 | 3 | 3 |
| *HIVEP3* | 7221 | 0.84 | 1.04E-03 | 2 | 0 | 0 | 0 | 2 | 4 |
| *MKI67* | 9771 | 0 | 1.55E-03 | 2 | 1 | 0 | 3 | 4 | 3 |
| *CHD8* | 7746 | 1 | 3.04E-03 | 2 | 1 | 0 | 0 | 1 | 3 |
| *TRIO* | 9294 | 1 | 3.11E-03 | 2 | 0 | 4 | 0 | 11 | 2 |
| *DNAH9* | 13461 | 0 | 5.07E-03 | 2 | 0 | 6 | 1 | 8 | 7 |
| *KIAA1109* | 15018 | 0.72 | 7.70E-03 | 2 | 1 | 1 | 0 | 14 | 12 |
| *KMT2C* | 14736 | 1 | 8.13E-03 | 2 | 2 | 0 | 0 | 6 | 7 |
| *TTN* | 107976 | 0 | 1.83E-01 | 2 | 3 | 14 | 8 | 59 | 58 |

## Discussion

By combining the Taiwanese and published SCZ trio cohorts, we report on 2772 trios, the largest exome-wide SCZ study on coding DNMs to date. SCZ trios show a significant, although modest, elevation of PTV and missense DNMs over both DNM model expectations and published control trios. *In silico* predictors of DNM deleteriousness, such as absence in exome reference panels, missense damaging predictors, and near splice-site synonymous changes, are all further enriched among SCZ cases above controls. Gene set analysis reveals that the SCZ DNM risk is broadly enriched for highly brain-expressed and evolutionarily constrained genes, and further enriched within putatively synaptic pathways and among genes identified as DNM risk factors in other neurodevelopmental disorders. In general, however, the signal is spread across many genes, and not confined to a specific cell type or curated biological pathway. Despite an elevation of genes recurrently hit by DNM PTVs, no single gene association surpasses genome-wide significance. These findings further support the polygenic nature of SCZ risk and suggest that larger samples will be required before individual gene associations will be discovered.

Exome sequence of SCZ case-control samples offers a natural comparison of the efficacy of different study designs and ascertainment criteria. The exome analysis of a large Swedish SCZ cohort (Genovese, et al. 2016) examined 4.9k SCZ cases and 6.2k controls. In parsing the contribution of rare coding variation to individual SCZ diagnoses, the Swedish SCZ cohort estimated that ∼10% of damaging and disruptive ultra-rare variants (dURVs) analyzed are DNMs, leaving 90% as inherited variants. One question that arises is how much DNMs are driving the enrichment seen among dURVs. Among the SCZ trio cohorts, we find a per-trio rate of 0.38 damaging and disruptive DNMs among SCZ probands compared to 0.3 per-trio in controls (fold enrichment=1.25), whereas case-control exomes find 0.25 additional dURVs in cases over controls (fold enrichment=1.07). Thus, our current estimate is that approximately 1/3 of the enrichment (or 0.08 of the 0.25 dURVs) observed in the case-control exomes are DNMs. While this difference supports the notion that damaging and disruptive DNMs are more penetrant than inherited variants in terms of their influence on an SCZ diagnosis, the general pattern is that both *de novo* and inherited rare coding variants confer a small, but significant impact on SCZ risk. When we consider the pattern of enrichment among various gene sets analyzed, most show a consistent direction and effect size in both trio DNM and case-control dURVs, pointing towards a similar polygenic burden of rare variation in brain expressed and constrained gene sets (**supplementary section 19**). Overall, these results indicate that a similar pattern of polygenic burden emerges across study designs, and for rare variant analysis in SCZ, both case-control and trio-based studies are likely to reveal a similar genetic signature of rare variation conferring risk for SCZ.

Outside of sequence data, structural variants analyzed from genotype array studies provides another source of rare genetic variation in SCZ. A mega-analysis of rare copy number variation (CNV) in SCZ case-control data from the Psychiatric Genetics Consortium (PGC; (Marshall, et al. 2017)) has refined the established association of rare CNV with SCZ. While a handful of previously SCZ-associated CNV (most co-morbid with severe neurodevelopmental features) are confirmed in the mega-analysis, there persists a modest genome-wide enrichment of ultra-rare CNV (defined there as MAF < 0.1%, OR=1.11, *p*=1.3e-7) overlapping genes outside of known disease-relevant CNV hotspots. Provided these ultra-rare CNVs are presumably a mixture of *de novo* and inherited events, the enrichment observed is consistent with coding DNMs described here, and further supports the notion that rare coding variants confer a small, but significant impact on SCZ risk.

One key consideration is the role that cognitive impairment plays on DNM rates within SCZ probands. Fromer et al. (Fromer, et al. 2014) observed that the elevated rate of PTV mutations in the Bulgarian SCZ trio cohort were largely confined to individuals with lowest scholastic attainment, suggesting that the enrichment of deleterious coding DNMs in SCZ may simply be from a subset of individuals with more severe cognitive impairment that also receive a diagnosis of SCZ. This finding has been further supported when combined with case-control SCZ exomes (Singh, et al. 2017), and is more prominent in autism probands, where the enrichment in PTV DNMs is confined to autism probands with lower IQ (Iossifov, et al. 2014; Robinson, et al. 2014; Samocha, et al. 2014; Kosmicki, et al. 2017). While direct measures of cognitive ability were not collected in the Taiwanese cohort, measures of sustained attention and executive function were available for most trios, providing a proxy for cognitive ability (**supplementary section 9**). Among these trios, we find enrichment in missense DNMs among low scores of sustained attention, and enrichment in PTV DNMs among low scores on executive function, particularly among brain expressed and constrained genes, providing further support that the elevated rate of DNMs in SCZ probands is, in part, a reflection of co-morbid intellectual impairment within these ascertained cohorts.

Overall, the rates and patterns of coding DNMs in SCZ probands reveal no loci of large effect, but rather a modest elevation of risk for DNM carriers, and one that is largely polygenic in its genetic architecture. The overlap in relative risk and gene set enrichment between trio and case-control designs so far strongly suggests that either approach will ultimately identify individual risk genes in larger samples.

## Methods summary

### Phenotype assessment

SCZ probands were recruited from hospitals, community care centers, and primary care clinics throughout the island country of Taiwan. Clinical data, blood sample collection, and research diagnostic assessment was conducted at the Department of Psychiatry, National Taiwan University Hospital and College of Medicine, National Taiwan University, Taipei, Taiwan. Of the 3,008 affected proband trios collected during the 5-year period of sample collection, 1,732 trios were enrolled in the current study, with 91% of trios reporting no family history of SCZ.

### Exome Sequencing, variant identification

Trios were sequenced at the Broad Institute using paired-end whole-exome sequencing on the Illumina HiSeq (1,140 trios) and X10 (598 trios) sequencers. For the Illumina HiSeq reads, we applied the Agilent SureSelect Human All Exon v.2 kit for exome capture. For the Illumina X10 reads, we applied the Illumina Nextera/ICE exome capture. Sequence reads were aligned with human reference genome (hg19) and variants called using the BWA/Picard/GATK pipeline. After sequence generation, 1,695 trios passed all quality control criteria for DNM analysis.

### Detection, annotation, and validation of DNMs

Identification of putative DNMs was done using custom Python scripts, incorporating posterior probabilities of being a true DNM based on population allele frequency from the NHLBI exome sequencing project. DNMs were annotated annotated using the Variant Effect Predictor (VEP) version 81 using GENCODE v19 mapped to the GRCh37 genome build, and validated using targeted amplicon re-sequencing. Unconfirmed DNMs were further validated using Sanger sequencing.

### Modeling rate expectations of DNMs

Expectation of DNM was determined using a mutational model framework across the exome. Details of the model are described in the supplementary information. In brief, the model incorporates the trinucleotide context of a given base position to assign probability of mutation to another base. For DNM burden across the exome, rates of DNM from published controls were also used as a comparison.

### Testing DNM rates and patterns

Per-exome DNM rates and single gene tests were compared against the mutational model, assuming a Poisson distribution of mutational events. Gene set tests incorporate a binomial model of greater than expected DNMs relative to mutational model expectations or published control DNMs.

### Pathway analyses

Pathway analyses were divided into two sets: ‘candidate’ gene sets and ‘discovery’ gene sets. Candidate gene sets represent previously reported gene sets associated with schizophrenia and other neurodevelopmental disorders. Discovery gene sets are derived from Gene Ontology (GO) annotations (http://www.geneontology.org/GO.downloads.ontology.shtml) and SynaptomeDB (http://metamoodics.org/SynaptomeDB/index.php). Candidate and discovery gene sets were corrected for multiple testing independently of each other.

## Supporting information

## Acknowledgements

This study was supported by grants from The National Human Genome Research Institute (U54 HG003067, R01 HG006855), The Stanley Center for Psychiatric Research, and the National Institute of Mental Health (R01 MH077139, R01 MH085521, and RC2 MH089905).

## Author contributions

N.L., J.L.M., S.V.F., S.J.G., S.A.M., M.T., and B.M.N. initiated the project. H-G.H. and W.J.C. led sample recruitment in Taiwan.

H-G.H., W.J.C., S.V.F., S.J.G., N.L., and M.T., provided the sample and phenotype collection.

F.C., K.C., S.D.C., A.D., J.L.M., and C.R. managed the sample collection and processing.

D.P.H., K.E.S., and M.F. processed sequence data and generated de novo mutation calls.

S.A.R., F.C., and S.A.M. undertook validation of mutations and additional lab work.

D.P.H. undertook the main bioinformatics/statistical analyses in close co-ordination with K.E.S., M.F., G.G., J.A.K., T.S., and B.M.N.

The main findings were interpreted by K.E.S., M.F., M.J.D., G.G., J.K., T.S., S.V.F., S.J.G., and B.M.N.

D.P.H. drafted the manuscript in close co-ordination with B.M.N. and C.C., with editing assistance from S.A.R., K.E.S., G.G., J.A.K., S.V.F., and S.J.G.

## Materials and correspondence

Daniel P. Howrigan

Benjamin M. Neale

