## Supplementary material for "Schizophrenia risk conferred by protein-coding *de novo* mutations"

**Supplementary Information**

**Supplementary Sections**

1. Taiwanese Trio sample ascertainment
2. Exome sequence generation and variant calling
3. DNM filtering
4. DNM validation
5. DNM annotation
6. Incorporation of published DNM
7. Statistical tests used to analyze DNM rates and patterns
8. Mutation rate model testing
9. DNM burden and covariates in Taiwanese probands
10. DNM burden in combined SCZ cohort
11. DNM burden in predicted damaging missense sites
12. DNM burden in predicted functional synonymous sites
13. SCZ enriched DNM subset
14. DNM burden in candidate gene sets
15. DNM burden in GO and SynaptomeDB databases
16. Single gene DNM enrichment
17. Gene recurrence enrichment analysis
18. Ultra-rare variant transmission among recurrently hit genes
19. Contribution of coding DNM to SCZ risk

**Supplementary Figures**

1. Supplementary Figure 1: SCZ vs control synonymous DNM rate – section 6
2. Supplementary Figure 2: Taiwanese vs published SCZ synonymous DNM rate – section 6
3. Supplementary Figure 3: Mutation model coverage simulation – section 8
4. Supplementary Figure 4: Mutation model gene set size simulation – section 8
5. Supplementary Figure 5: Exome wide coverage adjustment – section 8
6. Supplementary Figure 6: Exome-wide burden in Agilent capture – section 9
7. Supplementary Figure 7: Parental age on DNM burden – section 9
8. Supplementary Figure 8: Participant sex on autosomal DNM burden – section 9
9. Supplementary Figure 9: Participant sex on X chromosome burden – section 9
10. Supplementary Figure 10: Family history of mental illness on DNM burden – section 9
11. Supplementary Figure 11: DNM burden on CPT scores of sustained attention – section 9
12. Supplementary Figure 12: DNM burden on WCST scores of executive function – section 9
13. Supplementary Figure 13: Synonymous DNM rate by study – section 10
14. Supplementary Figure 14: Taiwanese vs. published SCZ DNM rate – section 10
15. Supplementary Figure 15: Missense predictor enrichment – section 11
16. Supplementary Figure 16: Non-ExAC missense predictor enrichment – section 11
17. Supplementary Figure 17: Synonymous DNM enrichment at NSS-ESR sites – section 12

18. Supplementary Figure 18: Synonymous annotation enrichment – section 12
19. Supplementary Figure 19: SCZ enriched annotation – section 13
20. Supplementary Figure 20: FWER passing gene set enrichment – section 14
21. Supplementary Figure 21: Evolutionary constraint gene set enrichment – section 14
22. Supplementary Figure 22: Evolutionary constraint gene set PTV enrichment – section 14
23. Supplementary Figure 23: Brain expression gene set enrichment – section 14
24. Supplementary Figure 24: Neurodevelopmental disease gene set enrichment – section 14
25. Supplementary Figure 25: Synaptic enrichment by exome coverage – section 14
26. Supplementary Figure 26: Neurodevelopmental disease gene set enrichment by exome coverage – section 14
27. Supplementary Figure 27: GO/SynaptomeDB gene set model enrichment – section 15
28. Supplementary Figure 28: GO/SynaptomeDB gene set case-control enrichment – section 15
29. Supplementary Figure 29: CPT scores of recurrent PTV gene carriers in Taiwanese cohort – section 17
30. Supplementary Figure 30: WCST scores of recurrent PTV gene carriers in Taiwanese cohort – section 17
31. Supplementary Figure 31: Synonymous transmission by variant quality score – section 18
32. Supplementary Figure 32: Gene set model comparison with SCZ case-control exomes – section 19
33. Supplementary Figure 33: Gene set case-control comparison with SCZ case-control exomes – section 19

**Supplementary tables**

1. Supplementary Table 1: Filtered DNM posterior probabilities prior to validation – section 3
2. Supplementary Table 2: DNM validation counts – section 4
3. Supplementary Table 3: Mutation model simulation results – section 8
4. Supplementary Table 4: Multiple regression model predicting DNM count – section 9
5. Supplementary Table 5: DNM burden partitioned by presence in non-psych ExAC cohort – section 10
6. Supplementary Table 6: Gene recurrence rates in highly brain expressed genes not under constraint – section 17
7. Supplementary Table 7: Ultra-rare variant transmission rates in recurrently hit genes – section 18

**Supplementary spreadsheets**

1. Supplementary Spreadsheet 1: Sheet Descriptions – section 2

2. Supplementary Spreadsheet 2: Taiwanese cohort sample list – section 2
3. Supplementary Spreadsheet 3: Taiwanese cohort trio list – section 2
4. Supplementary Spreadsheet 4: Taiwanese cohort DNM list – section 4
5. Supplementary Spreadsheet 5: DNM studies – section 5
6. Supplementary Spreadsheet 6: Combined cohorts DNM list – section 5
7. Supplementary Spreadsheet 7: SCZ DNM calls – section 10
8. Supplementary Spreadsheet 8: Candidate gene sets – section 14
9. Supplementary Spreadsheet 9: Candidate gene set results – section 14
10. Supplementary Spreadsheet 10: GO+SynaptomeDB enrichment – section 15
11. Supplementary Spreadsheet 11: Recurrent PTV genes – section 17
12. Supplementary Spreadsheet 12: Recurrent PTV+Missense genes – section 17
13. Supplementary Spreadsheet 13: Gene recurrence by gene set – section 17

Spreadsheets are also available as .tsv and .csv files at:
https://personal.broadinstitute.org/howrigan/taiwanese_trios_SCZ_supplement/

**Section 1 - Taiwanese Trio sample ascertainment**

3,008 parent-child trios, consisting of offspring diagnosed with schizophrenia and unaffected parents, were recruited from mental hospitals, community care centers and primary care clinics across the Island country of Taiwan. Of these 3,008 trios, 1,151 had no reported family history of Schizophrenia in their immediate family, and were recruited for the current project. The Department of Psychiatry, National Taiwan University Hospital and College of Medicine, National Taiwan University, Taipei, Taiwan served as the headquarters for data collection and research diagnosis.

Being the first report for this specific cohort from Taiwan, overall demographic information regarding the ascertainment of samples is listed below. The island country of Taiwan is situated in the Pacific Ocean about 160 km from the southern coast of the Chinese mainland. As of September 2014, the population size of Taiwan is 23,271,643. Recruitment catchment areas included metropolitan Taipei (population 7,030,620), Northern Taiwan (population 3,588,318), Middle Taiwan (population 4,520,155), Southern Taiwan (population 3,387.035), KaoPing (population 3,728,269) and Eastern Taiwan (population 1,017,246). In these six recruitment areas, 76, 28, 45, 31, 36, and 24 clinical settings participated in recruitment, respectively. Successfully recruited probands for trio analysis was 704, 558, 569, 576, 497 and 104, respectively. In turn, the recruitment ratio of probands per 10,000 in the population is 1.0, 1.5, 1.26, 1.70, 1.33, and 1.02, respectively.

<u>Clinical ascertainment and diagnostic assessment</u>

The clinical ascertainment of the probands and the trio family was comprised of three stages: (1) Obtaining IRB approval and informed consent, (2) Recruitment of probands diagnosed with schizophrenia and their parents, and (3) Collection of clinical data and diagnostic assessment.

**Stage 1**: We obtained approval from all IRBs of the hospitals participating in this study for proband and parent recruitment, data and sample collection, and informed consent approval for both proband and parents. The major content of the informed consent form composed of the following elements: goal of the study, inclusion/exclusion criteria of study subjects, methods of the study (diagnostic interview, blood sample collections), management of any remaining samples (e.g. DNA samples), potential adverse effects of study activities and its management, the expected results, requested activities of the recruited participants of this study, confidentiality, compensation and insurance of the participants for any adverse effect encountered in this study, rights of the recruited participants, withdrawal and termination for participating this study, and de-linking the basic identifiable personal data from the sample. Of the 813 IRB approvals, almost all are written in Chinese, and can be provided upon request.

**Stage 2**: Recruitment of the probands and the parents spanned a 5-year period, starting in June 2009 and concluding in March 2014. Recruitment included clinical screening and obtaining informed consent. Clinical screening, using a clinical screening sheet, was employed to (a) exclude those potential subjects with an ancestor of aboriginal origin; (b) include those potential subjects fulfilling the DSM-IV criteria of schizophrenia for proband recruitment based on clinical observation and interview by the attending psychiatrist providing the psychiatric services.

After identifying the potential proband, the parents were informed about the details of this study, and initial oral consent was obtained. The proband and the parents were then given the IRB-approved consent forms and the details of this study were explained. We then obtained the signed informed consent documents, and the proband joined this study.

**Stage 3**: Collection of clinical data and diagnostic assessment consisted of four main steps.

(1) A DIGS (Diagnostic Interview for Genetic Studies) interview of each proband was performed by trained research assistants with backgrounds in nursing, psychology, or social work. Following the interview, a review of clinical chart records in the hospital or in the clinical service settings was made prior to completing the clinical summary.

(2) Blood samples were drawn for DNA analysis.

(3) Two board-certified psychiatrists independently completed an initial research diagnostic assessment based on integrated clinical information in the DIGS interview data and summary notes of clinical course, symptom manifestations, and social functioning derived from the records of medical charts. If both research diagnostic assessments reached a consensus diagnosis of schizophrenia, the research diagnosis was finalized. If there was a discrepancy in the diagnostic assessment, then the case was subject to the last step of research diagnosis assessment.

(4) The last research diagnostic assessment was done by senior research psychiatrist, Professor Hai-Gwo Hwu, based on the information in the clinical screening sheet, the DIGS interview data, and the clinical summary note. If necessary, the research psychiatrist would call up the field-attending psychiatrists for clarification of clinical information crucial for the diagnostic assessment. Twenty-seven subjects were excluded from this study due to the diagnostic deviation from schizophrenia. After exclusion, 3008 trio families in total were recruited for research.

Parents of probands were not subject the interview stage (stage 3) of diagnostic assessment, and designation of a schizophrenia diagnosis in parents was based on family history information. From this information, 1151 trios were identified with unaffected parents, and recruited for exome sequencing.

**Section 2 – Exome sequence generation and variant calling**

After sample collection, blood DNA samples were shipped to Rutgers University Cell and DNA Repository. Samples identified for project inclusion were sent to the Broad Institute for exome sequencing.

Exome capture for waves 1 and 2

Exome targeting of DNA for the first two waves of data used hybrid selection capture from the Agilent SureSelect Human All Exon v2, targeting 33Mb of exon sequence, or 99% of human exons as defined by NCBI September 2009 consensus. Illumina HISeq 2000 sequencers were used to generate sequence reads, producing paired-end reads spanning 76 bases on average, with coverage goals of 20X sequencing depth or greater for at least 80% of the exome target. Sequencing was performed in two separate waves, consisting of 582 and 558 trios, respectively. Trios that did not meet coverage goals in initial sequencing passes were attempted in subsequent passes.

Exome capture for wave 3

Exome targeting of DNA for the third wave of data used hybrid selection capture using Illumina's Nextera Rapid Capture Exome v1 (ICE), targeting 37.7Mb of exon sequence. ICE capture covers the same exome target as Agilent SureSelect, with additional capture designed to include added content from the consensus coding sequence (CCDS) and RefSeq in the March 2012 UCSC genome database, as well as Gencode V11 database. Illumina X10 sequencers were used to generate sequence reads producing paired-end reads spanning 76 bases on average, with coverage goals of 20X sequencing depth or greater for at least 80% of the exome target. Sequencing was performed on 598 trios.

We performed additional quality control (QC) to confirm relatedness among trios, ensure DNA was whole blood, and sequence coverage was met. In all, 43 trios were removed in these QC steps, leaving 1695 trios for downstream analysis. On average, samples had 86% of the exome target meeting 20X sequencing depth (87% for Agilent captured samples and 84% for ICE captured samples), and 93% of the exome target meeting 10X sequencing depth (93% for both Agilent and ICE captured samples).

Sample information for the 1695 trios passing QC are available in **Supplementary spreadsheet 2: Taiwanese cohort sample list** and **Supplementary spreadsheet 3: Taiwanese cohort trio list**, with column level descriptions available in **Supplementary spreadsheet 1: Sheet_descriptions.**

Variant calling

The Burrows-Wheeler Algorithm (BWA: (Li and Durbin 2009)) was used to align unmapped sequence reads to the human reference (hg19), and the Picard pipeline (https://github.com/broadinstitute/picard) performed additional sequence QC checks and metrics to create a final BAM (Binary Sequence Alignment/Map format) file for each sample. GATK version 3.4 (McKenna, et al. 2010) was used to call single-nucleotide (SNV) and small insertion/deletion (indel) variants from individual BAM files, and recalibrated variant quality scores (VQSR) were generated to determine the overall variant quality in the VCF (variant calling format) file.

DNA Barcode switching

During the first wave of sequence generation, it was discovered that DNA barcodes inadvertently mislabeled individuals sequenced in the same lane during multiplex sequencing. During the process of adding DNA barcode identifiers to everyone's sheared DNA prior to sequencing, the active enzyme carrying the individual-specific barcode did not fully denature after bringing to higher temperature. Thus, when two samples were mixed (duplexed) together during sequence generation, there was still active enzyme labeling additional DNA fragments among the mixed DNA. This led to a non-trivial proportion of DNA barcodes of one individual attaching to the DNA of another individual, which we term "DNA barcode switching".

Quantifying the level of the DNA barcode switching was determined by examining the correlation coefficient between a) shared alleles of duplexed samples and b) 1K Genomes allele frequency. Once the enzyme protocol was fixed, evidence of DNA barcode switching disappeared. For the current analysis, DNA barcode switching presented a unique challenge to the discovery of *de novo* variation, as overall Mendelian error rates were increased across the board. Given that DNA barcode switching was restricted to sample pairs mixed in multiplex sequence, we could run direct comparisons of variant quality for *DNM*s between samples.

To identify *de novo* variants that were the result of barcode switching, we examined the variant quality of everyone's putative *de novo* call in their barcode switching partner (BSP). In general, variants present in an individual due barcode switching will be of lower sequence quality than the true variant. To start, we removed any *de novo* call where the BSP was called as homozygous alternate. We then compared the phred-likelihood (PL) score in both individuals, and removed *de novo* calls that had greater than 1.2 times higher PL score in the BSP. While many variants were removed using this approach, we still observed a significantly higher rate than expectation. In turn, we pursued an aggressive validation approach to confirm the presence of true *de novo* variants in the first wave of sequencing (supplementary section 4).

#### Section 3 – DNM filtering

##### Summary

For all three cohort waves, genotype calls were required to have a minimum PHRED-scaled likelihood (PL) > 20, analogous to at most a 1% probability of being a false genotype call. Allelic depth, or the proportion of non-reference genotype reads, was required to be at least 20% in offspring, and no more than 5% in either parent. We also removed sites where the offspring had < 10% read depth than the combined parental read depth, suggesting poor sequence capture or copy number deletion in the offspring.

<u>Cohort waves 1 and 2</u>: *De novo* calling did not use the conditional probability model or ExAC allele counts as filters (outlined below), and validation was pursued for all putatively called *de novo* variants. Validated calls, even if not passing all filters in the final combined VCF, were retained in the final set.

<u>Cohort wave 3</u>: *De novo* calling used the conditional probability model and the ExAC allele counts as filters. Validation was pursued for all putatively called *de novo* variants. Validation was pursed only for protein truncating variants.

##### Collapsing multiple DNMs in a single gene within a single individual

A small fraction of DNM calls from a single individual were observed in the same gene, often adjacent to one another. Most of these calls were confirmed in validation, suggesting a more “complex” *de novo* event is occurring at the site. Similar events have been reported and confirmed in previous studies (Fromer, et al. 2014). Following previous studies, we selected the single DNM with the most severe consequence in our final call set. In all, we observed 27 DNMs from 12 individuals considered “complex” DNMs, and selected 12 DNMs that had the most severe consequence (any ties were broken by selecting the higher depth call).

##### DNM filtering using posterior probability calculation

*De novo* single nucleotide variants (SNVs) and short indels were identified in the VCF calls, whereby the proband offspring genotype was heterozygous while the parental genotypes were both homozygous reference that the genotype call. Additional QC was performed to filter out genotype calls that were likely to be sequencing artifacts and not true DNMs. The filtering criteria used have been described in detail in previous studies (Neale, et al. 2012; Fromer, et al. 2014), with the current study making only minor adjustments to the cutoffs used.

The hard filters described above are less stringent than previous studies, as we include an independent Bayesian probability estimate informed by the minor allele frequency (MAF) to estimate whether the observed *DNM* is in fact real or more simply a missed heterozygote call in the parent. The method has been implemented and described in a recent autism paper (De Rubeis, et al. 2014). The algorithm, listed in equation 1, compares the posterior probability of a called *de novo* variant (equation 2) to the prior probability of the site already being variant in the population and a missed heterozygote (or het) call in one or both parents (equation 3). Using Bayes theorem, the conditional probabilities are used to calculate the relative probability of being a true *de novo* variant in the offspring:

[1] *p*(true *de novo*) = *p*(*de novo* | data) / (*p*(*de novo* | data) + *p*(missed het in parent | data))

The posterior probability, *p*(*de novo* | data), is the product of the three PL scores (PL father het call * PL mother het call * PL child reference hom call) in the VCF , *p*(data | *de novo*), multiplied by the probability of a mutation in the exome, *p*(*de novo*), which is estimated at 1 in 30Mb of diploid sequence (the per-exome mutation rate). Higher numbers increase the likelihood of being a true *de novo* event.

[2] *p*(*de novo* | data) = *p*(data | *de novo*) * *p*(*de novo*)

The prior probability, *p*(missed het in parent | data), is the sum of either parent being a missed het call in the VCF multiplied by the estimated MAF in the population. Similar to equation 2, the conditional probability rests upon the observed likelihood of a missed heterozygote call in at least one parent, *p*(data | missed het in parent), using the PL scores in the VCF (e.g. (PL father hom call * PL mother het call + PL Mother hom call * P mom_ref) * PL child reference hom call). This number is multiplied by the prior likelihood of at least one parent being het in the population, *p*(one parent het), which is 1 – (1 - MAF)^4. To calculate MAF, we used the maximum MAF estimated from either the Taiwanese trio VCF calls or from the extensively curated National Heart, Lung, and Blood Institute (NHLBI) Exome Sequencing Project (ESP) reference sequence database. In short, higher MAF variant calls increase the prior probability of being an inherited variant, thus lowering the probability of being a true *DNM*.

[3] *p*(missed het in parent | data) = *p*(data | missed het in parent) * *p*(one parent het)

Relative probabilities of *p*(true *de novo*) vary from 0 to 1, and are split into three broad categories. HIGH *de novo* variant calls have *p*(true *de novo*) of 0.99 or greater, MEDIUM *de novo* variant calls have *p*(true *de novo*) between 0.5 and 0.99, and LOW *de novo* variant calls have *p*(true *de novo*) below 0.5. The proportion of coding *de novo* events falling into each category is listed in **Supplementary Table 1: Filtered DNM posterior probabilities prior to validation**.

**DNM filtering using ExAC cohort allele frequencies**

While the initial *de novo* calling algorithm above filters out most false positive *de novo* calls, we pursued several additional steps to ensure a high-quality set of filtered calls in wave 3. We used allele count information from the exome aggregation consortium (ExAC v0.1; (Lek, et al. 2016)) to filter out additional variant sites. Of note, we excluded the contribution of the Taiwanese trio sample to the ExAC dataset when filtering variants. For LOW and MEDIUM confidence calls, we only included sites with at least 30% non-reference genotype reads, a singleton allele count in the Taiwanese trio sample, and allele count < 10 in ExAC. For HIGH confidence calls, we did not filter on allele balance, but retained only calls with a singleton allele count in the Taiwanese trio sample, and allele count < 10 in ExAC.

**Supplementary Table 1: Filtered DNM posterior probabilities prior to validation**

| Taiwanese cohort | Trios | HIGH SNV | HIGH indel | MEDIUM SNV | MEDIUM indel | LOW SNV | LOW indel | total |
|---|---|---|---|---|---|---|---|---|
| Agilent wave 1 | 575 | 547 | 29 | 61 | 5 | 391 | 27 | 1060 |
| Agilent wave 2 | 532 | 488 | 13 | 26 | 3 | 85 | 13 | 628 |
| Nextera wave 3 | 588 | 602 | 33 | 2 | 6 | 6 | 4 | 653 |

**Table legend**: Wave 1 and 2 DNMs were called and underwent validation prior to using either the posterior probability calculation or ExAC filtering, thus explaining the higher rate of LOW SNVs/indels in the putative DNM call set. In addition, barcode switching in wave 1 dramatically increased the number of LOW SNVs/Indels despite the additional filtering with the barcode switching partner.

**Section 4 – DNM validation**

**Cohort waves 1 and 2**

For the first two waves, we performed molecular validation of 1688 putative *DNM*s using an Illumina targeted amplicon sequencing framework. With the presence of DNA barcode switching in most of wave 1, we expected a substantial fraction of putative DNM calls in this cohort to be rejected in the validation process.

First, target region specific primers were designed using Primer3 (Untergasser, et al. 2012) and checked for specificity using the in-silico PCR tool from the UCSC Genome Browser. Oligo tails were added to the designed primers for subsequent hybridization to Illumina specific adapter sequence. Target regions were amplified in a first-round PCR for each father, mother, and child corresponding to a given DNM call. These PCR products were pooled by individual of origin, purified using the Agencourt AMPure XP system, then put into a second-round PCR in which Illumina specific adapter sequence and barcodes were added via hybridization to the complementary oligo tail sequence. After purification of the round 2 PCR products, completed libraries were pooled, with quantification and quality control being done by QuBit fluorometric quantitation, Agilent BioAnalyzer high sensitivity DNA kits, and Kapa Biosystems Illumina library quantification kits. Amplicon libraries were loaded on a MiSeq or HiSeq 2500 with a 5% PhiX spike in for sequencing.

Demultiplexed fastq files from the validation sequencing runs were aligned with BWA-MEM (Li and Durbin 2009), reads with greater than 3 soft clipped bases in the beginning of the alignment were removed to control for sequencing artifacts. Allele pile-ups downsampled to 1000 were counted using the GATK UnifiedGenotyper (McKenna, et al. 2010) in a targeted fashion after filtering for mapping quality, base quality, and edit distance. Only sites with at least 30 reads overlapping the variant site were considered. Genotype calls were made based on allelic fraction of reads at each locus, with a non-reference allele fraction within 30-70% being considered heterozygous. All target sites were visually inspected in IGV (Robinson, et al. 2011) for accuracy, with a selected subset of calls confirmed by Sanger sequencing.

**Cohort wave 3**

For the third wave of trios, validation was pursued specifically on protein-truncating variants, and use more stringent filtering criteria as a general method to keep the false positive call rate low. Primers were designed for 68 PTV variants, with validation performed through an external contract with Sequenom. The 9 variants that didn't return a confident call either way were followed up using Sanger sequencing. In all, we validated 59 PTVs (86.8% validation rate) from the putative DNM list.

**Supplementary Table 2: DNM validation counts**

| Taiwanese cohort | Trios | Putative called DNMs | DNMs submitted for validation | Validated DNMs | Final DNM count |
|---|---|---|---|---|---|
| Agilent wave 1 | 575 | 1060 | 1060 | 586 (55.3%) | 586 |
| Agilent wave 2 | 532 | 628 | 628 | 517 (82.3%) | 517 |
| Nextera wave 3 | 588 | 653 | 68 | 59 (86.8%) | 644 |

Final DNM calls for the full Taiwanese trio cohort are listed in **Supplementary Spreadsheet 4: Taiwanese cohort DNM list.**

#### Section 5 – DNM annotation

##### Primary annotation - Variant Effect Predictor (Ensembl VEP) annotation

Putative DNMs were primarily annotated using the Variant Effect Predictor (VEP) version 81, which uses GENCODE v19 mapped to the GRCh37 genome build. All non-coding variants were removed from further analysis. Remaining variants comprise three broad categories – synonymous, missense, and protein-truncating variants. Synonymous annotation includes synonymous amino acid changes and stop-retained changes. Missense annotation includes non-synonymous single amino acid changes, inframe insertions and deletions, and stop-lost changes. Protein-truncating variant (PTV) annotation includes stop-gain, start-lost, frameshift, essential splice site changes. To maintain consistency, annotation was updated for published DNMs, resulting in a few minor changes to published annotations.

##### Missense prediction annotation

Within missense mutations, we used annotations curated through the dbNSFP v2.9 database (Liu, et al. 2011, 2013). We assessed PolyPhen2 HDIV and HVAR, SIFT, LRT, MutationTaster, MutationAssessor, FATHMM, MetaSVM, MetaLR, Provean, and CADD v1.2 predictions and scores to measure predicted pathogenicity.

##### Secondary annotation

Secondary analyses of published synonymous DNMs reported a significant enrichment in near-splice site synonymous changes among ASD probands, as well as enrichment in DNAse Hypersensitivity sites (DHS) drawn from the cerebrum, cerebellum, and frontal cortex region of post-mortem tissues in SCZ probands (Takata, et al. 2016). Following their protocol, we annotated the distance to the nearest splice site using SeattleSeq Annotation 137 (Ng, et al. 2009), exon-splicing enhancers (ESE) / silencers (ESS) from hexamer motifs (Fairbrother, et al. 2002; Wang, et al. 2004; Ke, et al. 2011), and candidate DHS regions from ENCODE Experiment Matrix (**https://genome. ucsc.edu/ENCODE/dataMatrix/encodeDataMatrixHuman.html**).

##### ExAC allele frequency

We also used the ExAC v0.3 allele frequency to further interpret the potential for pathogenicity of the *DNM*. The first two waves of the Taiwanese cohort were included in the ExAC database, and consisted predominantly of parents. For trios with high levels of DNA barcode switching in the first wave, the ExAC analysts prioritized individuals with the lowest amount of barcode switching, and in some cases, selected only the proband for inclusion in the ExAC callset. Fortunately, we could account for all these trios, and adjust the ExAC allele count to exclude these individuals. Furthermore, in the non-psychiatric version of the ExAC database, the Taiwanese trios were not included. For exome-wide burden and gene-set analyses, we examine the role of variants seen in ExAC and those not seen in ExAC.

**Section 6 – Incorporation of published DNM**

Along with our mutational model, we also wanted to assess how the patterns and rates of observed DNMs compare with published control trios and unaffected siblings in the published literature, as well as incorporate published *DNM*s in SCZ to further refine their impact of on SCZ risk. We collected results from independent exome-sequenced SCZ trios, their unaffected siblings and control trios, as well as unaffected siblings collected from other neuropsychiatric disorders. Specific publications and descriptive data are listed in **Supplementary Spreadsheet 5: DNM studies**. In total, we assessed 1077 published SCZ trios and 2216 control trios and unaffected siblings of ASD probands.

To assess overall DNM rates, we performed cross-study comparisons and checked against the mutational model expectation. Datasets with significantly different rates when compared with similar studies and against the mutation model were flagged and not incorporated in exome-wide DNM rate calculations, but were retained for single gene and gene-set analyses (**Supplementary Figure 1: SCZ vs control synonymous DNM rate**). Within the Taiwanese trio cohort, the wave 3 cohort had a significantly higher synonymous rate due to the larger exome target in Nextera capture (37.7 Mb vs 33 Mb in previous waves). When we restricted to only DNMs within the Agilent capture targets, the synonymous rate was comparable to previous waves (**Supplementary Figure 2: Taiwanese vs published SCZ synonymous DNM rate**). We therefore restricted to only DNMs within the Agilent capture targets among all cohorts when analyzing exome-wide burden.

The full list of DNMs analyzed are available in **Supplementary Spreadsheet 6: Combined cohorts DNM list**

**Section 7 - Statistical tests used to analyze DNM rates and patterns**

All statistical tests were performed using R statistical software **(www.r-project.org)**. To examine overall rates, recurrence, and single gene enrichment of DNMs in SCZ probands, we fit the data to Poisson distributed models. To examine the role of covariates on DNM rates within the Taiwanese trio cohorts, we use a Poisson-distributed multiple regression model. For comparing against mutation model expectations, we used a one-sample exact Poisson test, with the mutation expectation as our lambda parameter. For comparing against control DNMs, we used a two-sample exact Poisson test. To test gene recurrence against the mutation model, we used a bootstrap re-sampling method to sample recurrent genes using per-gene probabilities and used the empirical distribution to test for significance.

To test for gene set enrichment, we fit the data to a binomial model. This model is conditional on the overall rate, and examines the relative proportions of enrichment rather than the observed rate. For example, if SCZ probands had twice the rate of PTV DNMs than controls, but 50% fell in brain-expressed genes in each sample, then we would have two-fold enrichment in the PTV rate, but no enrichment for brain-expressed genes among PTVs. For comparing against mutation model expectations, we used a one-sample exact binomial test, with the mutation expectation as our null probability of gene set overlap. For comparing against control DNMs, we used a two-sample exact binomial test.

Additional information about the tests used for each analysis are detailed in subsequent sections.

#### Section 8 – Mutation rate model testing

Accurately modeling the expected probability of a germline mutation in the exome is a valuable tool for testing the significance of observed mutations against the underlying expectation under the null. By measuring DNM expectation at the base position level, the model can be applied to individual genes, gene-sets, and across the entire exome. The mutational model used has been described in detail in (Neale, et al. 2012) and (Samocha, et al. 2014). Briefly, the model assigns a probability to each nucleotide base mutation based primarily on the tri-nucleotide context at the site (Krawczak, et al. 1998; Kryukov, et al. 2007). Informed by intergenic rates of base changes between orthologous chimp and human sequence in the 1000 Genomes project (Genomes Project, et al. 2015), predicted per-gene probabilities were correlated strongly with observed rats of rare (MAF < .001) synonymous variants of mutation ($r$ = 0.940), performing significantly better than gene length alone (Samocha, et al. 2014). The rates at which DNMs appear in the exome follow a Poisson distribution, whereby the expectation and variance are assumed to be equal. Mutation expectations are calculated by each segregating chromosomes, so genes on chromosome X need to account for the male/female ratio in the sample (DNMs on chromosome Y and MT were not examined). In turn, per-exome burden and single gene tests use a Poisson-modeled distribution and Poisson regression to test for significance.

##### Previous adjustment for sequence coverage in the mutation model

Exome sequence coverage varies across the exome target, the expected DNM rate needs to be adjusted to correspond to the proportion of the exome that was called with adequate sequencing depth and quality. Given that the capture method and sequencing depth differs across cohorts, mapping onto a single mutation rate model can lead to biased inference due to miscalibration of expected mutation rates.

Adjustments to the mutation model accounting for sequencing depth have previously used coverage information in the ExAC database, which was predominantly restricted to the Agilent SureSelect v2 capture target (Samocha, et al. 2014; Lek, et al. 2016). Mutation rates were adjusted by the proportion of trios in the ExAC database with 10x coverage at a given coding base, and tested against observed synonymous singletons in the Exome Sequencing Project (ESP) to assess the efficacy of adjustment. With the availability of exomes sequenced on Nextera capture, we also ran a similar coverage adjustment using samples with Nextera coverage, and created a hybrid adjustment model for both Agilent and Nextera adjustments.

We contrasted this coverage adjusted model (Agilent/Nextera adjusted model) against unadjusted mutation expectations from Gencode v19 canonical transcripts (the full model). In both models, we examined coverage for 19,166 autosomal and X-linked genes.

##### Sequence coverage simulation on mutation model

To assess the performance of both the full and Agilent/Nextera adjusted mutation expectations on the current dataset, we developed a simulation framework to evaluate how well both randomly generated gene sets and coverage-biased gene sets map onto the current DNM list. If the coverage adjustment to the mutation model is adequate, we should see no overlap enrichment of simulated coverage-biased gene sets relative to randomly generated gene sets, and both gene sets should not significantly differ from model expectations when tested against observed DNMs in SCZ probands. Deviation of random gene sets from expectation suggests a miscalibration of the mutation expectations, and deviation of coverage-biased gene sets from expectation suggest a miscalibration of the coverage adjustment.

To develop coverage-biased gene sets and test a variety of custom coverage adjustments, we used available sequence depth information from the three Taiwanese and six Bulgarian trio cohorts, comprising of 2312 SCZ trios (or 83.7% of SCZ probands assessed). For each cohort, we calculated per-gene coverage, which is the proportion of each gene covered at mean 10x depth in the cohort. Using the per-gene coverage in each cohort, we developed two coverage metrics. The first, which we call “weighted coverage”, determines the probability of being selected in the coverage-biased gene set. The second, which we call “coverage cutoff”, determines the inclusion of the gene in the model if it meets the specified coverage cutoff across trio cohorts.

Weighted coverage is the summed per-gene coverage by cohort divided by the full 2312 trios, and we used this as our probability for selecting a gene in the coverage-biased gene sets. As opposed to randomly selected genes, where the probability is equal for each gene (all weights=1), fully covered genes have equal probability to other fully covered genes (weight=1), whereas partially covered genes are penalized (weight < 1).

The coverage cutoff is the proportion of size-weighted cohorts meeting a specified amount of 10x coverage. The idea behind this metric is that if enough of a gene meets 10X coverage in a cohort, we should include the entire mutation expectation for that gene. For example, if the Taiwanese wave 1 cohort had 50% of a gene with 10X mean coverage, we would include all 575 trios this cohort for any cutoff 50% and below, but contribute 0 trios for anything above 50%. Mutation rates are then adjusted on a passing cohort level, where if 1256 trios (or 50% of 2312 trios) surpassed the coverage cutoff, the mutation rate would be multiplied by 50%. Any gene that had less than 20% of trios passing the coverage cutoff was subsequently removed from the model.

Simulated gene sets generated from 19,166 genes:

- Two gene set selection models
    - Random: genes selected with equal probability
    - Coverage-biased: genes selected with probability equal to weighted coverage
- Five different gene set sizes (1000, 2500, 5000, 7500, and 10000 genes)
- Created 1000 simulated gene sets for each setting

Two mutation models examined:

- Gencode v19 canonical transcript (Full model)
- Agilent/Nextera coverage adjusted (Agilent/Nextera adjusted model)
- All models are tested as is (0% coverage cutoff) up to 100% coverage cutoff

For each mutation model and coverage cutoff, we tested all 10 gene set configurations against the SCZ DNMs, comparing the difference between the observed proportion of DNMs to proportion predicted from the mutation model. For the 1000 gene sets, we retained the empirical mean and 95% CI of the differences. Genes in the mutation model removed due to not meeting the coverage cutoff we also removed from the observed DNM list. To determine the best-fit model, we first selected the coverage cutoff with the smallest mean difference between observed and expected proportions among the coverage-biased gene sets. To verify the consistency the best-fit model, we checked it against all five separate gene set sizes. We also ran the full simulation protocol against observed controls DNMs to gauge the consistency of the best-fit model.

**Sequence coverage simulation results**

For randomly generated gene sets, we see a modest, but significant deviation of observed DNMs above expectation in the coverage-adjusted model, whereas the full mutation model matched expectations **(Supplementary Figure 3: Mutation model coverage simulation)**. The bias in the coverage-adjusted model persisted across the additional coverage cutoffs made here, suggesting that the pre-existing coverage adjustment is slightly biased towards underestimating mutation probabilities in a uniform manner across all genes. Thus, the full model mutation expectations were preferred against the previously adjusted coverage model.

For coverage biased gene sets, the DNMs in SCZ are strongly enriched against both the full and coverage adjusted model, and additional coverage adjustment to the models are required to account for the bias that variable coverage has on observed DNMs. As expected, the depth-adjusted models do require less additional coverage adjustment than the full model to match the null expectation. The table below lists the optimal adjustment that was made to fully correct for coverage bias in each model when looking at our largest gene set simulation (10k gene set size).

Assessing both random gene set effects and coverage cutoff adjustments, the 75% coverage cutoff using the full model was our best performing model in the simulation, and follow up tests show that this model is consistently null across various gene set sizes (all $p > 0.05$), and performs reasonably well when comparing to observed control DNMs (**lowest $p = 0.01$; Supplementary Figure 4: Mutation model gene set size simulation**). This model incorporates 17,925 of the 19,166 genes (94%), with 13,655 (71%) genes using the full mutation model expectations (**Supplementary Figure 5: Exome wide coverage adjustment**) We used this adjusted mutation model in overall mutation burden, gene set, and single gene analyses.

**Supplementary Table 3: Mutation model simulation results**

| Model | Optimal coverage cutoff | Genes failing coverage | Genes adjusted for coverage | Fully covered genes | Per trio expectation |
|---|---|---|---|---|---|
| Full | 75% | 1239 (6.5%) | 4272 (22.3%) | 13655 (71.2%) | 0.952 |
| Agilent/Nextera | 60% | 746 (3.9%) | 3193 (16.7%) | 15227 (82.7%) | 0.938 |

**Table legend**: The full model uses Gencode v19 canonical transcript expectations, whereas the Agilent/Nextera adjusted model uses the combination of capture targets and coverage adjustment to derive mutation expectations.

#### Section 9 - DNM burden and covariates in Taiwanese probands

We wanted to see if DNM rates were influenced by individual level and experimental level variables available in the Taiwanese trio cohort. At the phenotype level, we had information on parental age, proband age at SCZ onset, gender, and a family history of mental illness (including, but not limited to a SCZ diagnosis). At the genotype level, we had information on sequencing coverage, estimated barcode contamination level, and parental DNA type. While we only retained probands with whole blood DNA in the analysis, trios with parents sequenced using lymphoblastoid cell line DNA were retained, as mutations arising in the parent from cell-passaging had negligible effects on the DNM rate. We also examined how DNM burden affected measures of sustained attention and executive function among the subset of probands with available information.

##### Overall DNM burden in Taiwanese trio cohorts

Among the three cohorts sequenced, we expected a higher rate of DNMs ascertained in cohort wave 3 due to the larger Nextera exome capture target (37.7 Mb) relative to the Agilent SureSelect v2 capture target (33 Mb). We controlled for this enrichment by restricting to DNMs seen within the hg19 Agilent v2 target intervals (**Supplementary Figure 6: Exome-wide burden in Agilent capture**). Subsequently, for all whole exome burden analyses moving forward, we restricted our analysis to DNMs within the Agilent target intervals. Further comparisons of DNMs to the mutation model and against controls are detailed in section 10.

##### Covariate association to DNM burden

Among the analysis covariates, we wanted to ensure that the number of DNMs did not significantly vary due to sex, parental DNA type, and estimated contamination rate. Given that barcode contamination occurred exclusively in the cohort wave 1, we examined two models, with the first model using sequencing wave as a covariate, and the second model using exome capture kit and estimated contamination level as covariates. Based on previous literature, we expect to see a nominal increase in the DNM rate with higher parental age, particularly in fathers (Fromer, et al. 2014; Francioli, et al. 2015).

To test for effects of individual and experimental level variables on the per-trio DNM rate, we fitted a multivariate generalized linear model using a Poisson link function, including all covariates in the primary model. Significant variables were followed up to further determine the source of influence on the DNM rate.

##### Results of covariates on DNM burden

Within the multiple regression framework including all covariates (exome capture and contamination, sequencing coverage, biological sex, family history of mental illness, parental age, age at onset, and parental DNA type), parental age was the strongest predictor of DNM rate ($\beta(1685)=0.03$, $p$=6.5e-7). No other covariates uniquely predicted DNM rates in the multiple regression (all $p$ > .05). Higher estimated contamination levels, which required additional QC and full validation to discover true DNMs, did predict a lower DNM rate, however the lower rate was not significant ($\beta$(1685)=-1.49, $p$=0.08). Earlier age of SCZ onset did not predict a higher DNM rate, and this comparison held up when we restricted to PTVs in constrained genes and compared early onset probands (the 26% with onset under 18 years of age) to the remaining probands ($\beta$(1694)=-1.2, $p$=0.51).

**Supplementary Table 4: Multiple regression model predicting DNM count**

| Covariate | $\beta$ | SE | *p*-value |
|---|---|---|---|
| Parental Age | 0.03 | 0.006 | 6.5e-7 |
| Sex is Male | -0.08 | 0.05 | 0.12 |
| Barcode switching contamination % | -1.45 | 0.85 | 0.09 |
| Age at onset | -0.004 | 0.004 | 0.29 |
| Has a family history of mental Illness | -0.15 | 0.10 | 0.14 |
| Percent 10X coverage | -2.77 | 2.28 | 0.22 |
| Parental DNA source (LCL) | 0.055 | 0.13 | 0.67 |
| Exome capture is Nextera | -0.03 | 0.07 | 0.70 |

Further evaluation of parental age showed that paternal and maternal age is strongly correlated ($r$=0.494), and both paternal age ($\beta$(1693)=0.025, $p$=7e-9) and maternal age ($\beta$(1693)=0.02, $p$=5e-4) predicted increased DNM rates in bivariate regression models (**Supplementary Figure 7: Parental age on DNM burden**). However when we fit paternal age and maternal age in the regression model, paternal age ($\beta$(1692)=0.023, $p$=3e-6) predicted increased DNM rates over and above maternal age ($\beta$(1692)=0.01, $p$=0.38). Further follow up on parental age effects revealed a modest quadratic effect of maternal age ($\beta$(1692)=0.002, $p$=0.01) but not for paternal age ($\beta$(1692)=2e-4, $p$=0.6). These results show a clear association of older paternal age and increasing DNM rates, and suggest that maternal age association to increased DNM rates acts in a non-linear fashion.

**PTV enrichment restricted to female SCZ probands**

Published Autism DNM studies show a female ‘protective’ effect, where a higher burden of deleterious DNMs are seen among females diagnosed with ASD relative to males (Jacquemont, et al. 2014; Robinson, et al. 2014). To see if this effect is present among SCZ probands in the Taiwanese cohort, we split by DNM burden by biological sex (662 females, 1033 males). We split autosomal and X-linked DNMs, as females are expected to have a 2 to 1 ratio of germline DNMs relative to males on the X chromosome while having 1 to 1 ratio on the autosomes. When we compare against the mutation model, we find that female probands are modestly enriched for PTV DNMs in both the autosome (fold-enrichment=1.33, $p$=0.01) and on the X chromosome (fold-enrichment=2.48, $p$=0.05), whereas male probands show no enrichment (autosomal fold-enrichment=1.03, $p$=0.78; X-linked fold-enrichment=0.64, $p$=1). Missense mutations show a similar enrichment in females, however to a lesser degree than PTVs (autosomal fold-enrichment=1.10, *p*=0.05; X-linked fold-enrichment=0.93, *p*=0.89; **Supplementary Figure 8: Participant sex on autosomal DNM burden**).

**Male probands have low rates of X-linked DNMs**

When we compare X-linked DNMs between males and females, females are significantly enriched for X-linked DNMs relative to males (28 to 8, fold-enrichment=2.73, *p*=0.01), with PTVs showing the largest difference (5 to 1, fold-enrichment=3.9, *p*=0.36). Note that this is after accounting for the inheritance of two X chromosomes in females and one in males. To understand whether this represents an enrichment in female DNMs or a depletion in male DNMs, we compared X-linked rates against the mutation model. Female rates are modestly above expectation (fold-enrichment=1.18, *p*=0.35), whereas males are significantly below expectation (fold-enrichment=0.47, *p*=0.02; **Supplementary Figure 9: Participant sex on X chromosome burden**). We could not find evidence that this depletion was due to a lack of sensitivity in DNM calling for males on the X chromosome, and suggest this depletion may be indicative of haploinsufficiency among males on the X-chromosome.

**Family history of mental illness has modestly lower rates of DNM burden**

151 of the 588 trios in cohort wave 3 reported having a history of mental illness in their family, while none of the trio in cohort waves 1 and 2 reported any history of mental illness. Each parent reported their family history, but do not detail the specific nature of mental illness running in the family. In theory, we expect that affected probands with no history of mental illness are more likely to incur genetic liability in the form of a DNM rather than inherited genetic variation. While we do not see a significant increase in overall DNM rates among probands with no family history of mental illness in the multiple regression model (OR=0.86, *p*=0.14), the direction did suggest a lower rate. Follow up analyses find that PTV (fold-enrichment=1.73, *p*=0.12) and missense (fold-enrichment=1.15, *p*=0.24) DNMs are suggestively higher in probands with no history of mental illness in their family, however neither reach statistical significance. Within cohort wave 3, where all trios having a history of mental illness in their family were sequenced, we do see a modestly significant enrichment in DNMs among probands with no family history of mental illness (fold-enrichment=1.26, *p*=0.02) with PTV, missense, and synonymous DNMs all contributing to this enrichment (**Supplementary Figure 10: Family history of mental illness on DNM burden**). Overall, the pattern is consistent with expectations, and the modest effect size is likely indicative of the relatively smaller effect of DNM on SCZ liability more broadly.

**Enriched DNM burden in probands with deficits in sustained attention and executive function**

Previous analysis of DNMs with IQ measures in ASD probands (Iossifov, et al. 2014; Robinson, et al. 2014; Samocha, et al. 2014; Kosmicki, et al. 2017), and scholastic achievement in SCZ probands (Fromer, et al. 2014) show co-morbidity with cognitive impairment among proband carriers of PTV DNMs, accounting for a substantial proportion of the DNM burden enrichment seen in these cohorts. Furthermore, the contribution of rare coding variants among SCZ cases show substantial overlap with a diagnosis of intellectual disability (Singh, et al. 2017). While no direct measures of IQ or scholastic achievement scores in the Taiwanese cohort were collected, tests of sustained attention in the Continuous Performance Test (CPT) and executive function from the Wisconsin Card Sorting Test (WCST) were assessed in patients. Details of the tests and the metrics ascertained have been previously described for both the CPT (Chen, et al. 2004) and WCST (Lin, et al. 2013), with a review of their utility as potential SCZ endophenotypes (Chen 2013) and association to SCZ polygenic risk scores among in the Taiwanese cohort (Wang, et al. 2018). For the current study, we examined Z-scores from each test that controlled for sex, age, and education. For the CPT, we used Z-scores from the *d′* measure of sensitivity, which is derived from the hit rate (probability of response to target trials) and the false-alarm rate (probability of response to non-target trials). Z-scores were quite similar between the undegraded and degraded test ($r$=0.73), and we used scores from the degraded test as they showed higher test-retest reliability in previous samples (Chen, et al. 2004). Test measures for the degraded CPT were available for 1306 probands. For the WCST, we averaged the Z-scores from Categories Achieved and Total Errors (reverse-scored) to capture both success and failure rates. Test measures for the WCST were available for 1327 probands.

To determine how DNM burden is associated with sustained attention and executive function, we ran a median split on each Z-score, creating a “high” and “low” score group. We then examined DNM burden between both groups and against mutation model expectations, restricting our burden analysis to DNMs within Agilent capture intervals and the 17925 well-covered genes. Along with exome-wide rates, we also restricted to two candidate gene sets enriched among SCZ probands (supplementary section 14). One gene set encompasses high brain expression, which is the inclusive combination of the BrainSpan high brain-expressed gene set and the GTEx brain enriched gene set, totaling 10376 of the 17925 well-covered genes. The other gene set, defined as genes under constraint, is the inclusive combination of RVIS intolerant, missense constraint, and pLI > 0.9 gene sets, totaling 4083 of the 17925 well-covered genes.

Among CPT scoring groups, we see an elevation in DNM burden among missense DNMs in low CPT scorers over high CPT scores (fold-enrichment=1.16, $p$=0.04) and the mutation model (fold-enrichment=1.11, $p$=0.03; **Supplementary Figure 11: DNM burden on CPT scores of sustained attention**). We did not find any difference in PTV DNM burden between low and high CPT scores. This elevation in missense DNMs for low CPT scorers is evident in both high brain-expressed and constrained genes. Among constrained genes, we also see an elevation of synonymous variants in low CPT scorers relative to high CPT scorers (fold-enrichment=1.6, $p$=0.01), contributing to a significantly higher DNM rate for low CPT scorers among all DNM in constrained genes relative to both high CPT scorers (fold-enrichment=1.29, $p$=0.006) and the mutation model (fold-enrichment=1.21, $p$=0.006). When we tested DNM burden against WCST scoring groups, we see an elevation in DNM burden among PTV DNMs in low WCST scorers relative to the mutation model (fold-enrichment=1.44, $p$=0.002) and against high WCST scorers (fold-enrichment=1.39, $p$=0.06; **Supplementary Figure 12: DNM burden on WCST scores of executive function**). We did not find any difference in missense DNM burden between low and high WCST scores. This elevation in PTV DNMs for low WCST scorers is particularly evident when we restrict to highly brain-expressed genes, both relative to the mutation model (fold-enrichment=1.84, $p$=7e-6) and against high WCST scorers (fold-enrichment=1.46, $p$=0.05), with the signal in constrained genes being similar (model fold-enrichment=1.80, $p$=0.001; two-sample fold-enrichment=1.67, $p$=0.05). Additional analyses regressing DNM burden on Z-scores directly, as well as splitting into tertiary groupings (bottom 3$^{rd}$ scores vs top 3$^{rd}$ scores), produced similar results (data unpublished).

Overall, the elevation of missense DNMs concentrated in low CPT scores and PTV DNMs concentrated in low WCST scores is consistent with previous observations that the enrichment of DNM burden in SCZ probands is, in part, co-morbid with evidence of mild cognitive impairment, suggesting that the subset of risk DNMs conferring risk to SCZ are also likely to confer risk to multiple neurodevelopmental disorders. While the scores used here serve as a proxy for cognitive ability, they nonetheless support the idea that large effect DNMs will subsequently affect a broad range of phenotypic outcomes related to brain function.

#### Section 10 - DNM burden in combined SCZ cohorts

Per-exome DNM rates were examined in aggregate, as well as within mutational types (synonymous, missense, PTV). Rates were tested against the calibrated mutational model using two-sided Poisson probabilities (see supplementary section 8). For all whole exome burden analyses, we restricted our analysis to DNMs within the Agilent v2 target intervals. To buffer against any potential biases reflected in the model, we also tested against DNM rates in control trios and unaffected siblings from published studies (described in detail below). Because per-trio counts were unavailable from many studies, we modeled the per-exome DNM rate as a Poisson distributed statistic, whereby the mean and variance are equal, and used a Poisson exact test to determine significance of DNM rates between SCZ probands in unaffected siblings/controls.

##### Study inclusion using synonymous DNM rate

By incorporating the Taiwanese cohorts with previously published DNM studies with both SCZ probands and unaffected siblings/controls, numerous factors can affect the observed rates of DNMs in each cohort (e.g. capture platform, depth of coverage, quality control). We used the synonymous DNM rate as a quasi-null control to see if any cohort is an outlier relative to other groups. Among all cohorts analyzed in this study, only one cohort (Xu, et al. 2012) showed a significantly lower synonymous rate relative to remaining cohorts (fold-enrichment=0.47, $p$=7.2e-6; **Supplementary Figure 13: Synonymous DNM rate by study**). We omitted this cohort from whole exome DNM burden, but retained it for gene set and single gene analyses.

##### DNM burden between Taiwanese cohorts and published SCZ studies

When we compare DNM burden to published SCZ DNM studies, none of the three Taiwanese cohorts differed significantly from the combined published SCZ cohort with respect to all coding, PTV, missense, and synonymous DNMs (all $p > 0.05$; **Supplementary Figure 14: Taiwanese vs. published SCZ DNM rate**), suggesting that the combination of published SCZ and Taiwanese cohorts does not introduce any significant bias with respect to whole exome burden. A full list of DNM counts among all SCZ cohorts is available in **Supplementary Spreadsheet 7: SCZ DNM calls**.

##### DNM rates enriched in SCZ cases relative controls and model expectations

When we compare the exome-wide DNM rates in the combined SCZ cohort against controls/unaffected siblings in the published literature, overall coding DNM rates in SCZ probands are significantly enriched relative to controls (fold-enrichment=1.08, $p$=0.009). When we partition by mutation type, PTV (fold-enrichment=1.28, $p$ = 0.21) and missense (fold-enrichment=1.09, $p$=0.02) DNM rates are significantly higher than controls, while synonymous DNM rates remain similar (fold-enrichment=1.05, $p$=0.44; **Supplementary Table 5: DNM burden partitioned by presence in non-psych ExAC cohort**). When we compare SCZ probands to the mutation model, we restricted to genes that met coverage across cohorts (supplementary section 8). Overall coding DNM rates in SCZ probands are not enriched over expectation (fold-enrichment=1.01, *p*=0.43). When we partition by mutation type, protein-truncating (fold-enrichment=1.18, *p*=0.01) and missense (fold-enrichment=1.06, *p*=0.03) DNM rates are significantly higher than expectation, while synonymous DNM rates are significantly lower (fold-enrichment=0.87, *p*=0.001). Given that this depletion in synonymous rates is consistent across both SCZ probands and controls, we don't think this effect is specific to SCZ, but more likely to represent a miscalibration of the mutation model for synonymous DNMs or upstream quality control in DNM discovery that penalizes synonymous DNM calls (such as filtering on allele frequency).

**DNM enrichment in SCZ probands are restricted to mutations not present in ExAC**

One empirical measure of DNM severity is whether the mutant allele is observed in a larger reference population. If a DNM, even with predicted pathogenic consequences, is seen in ostensibly healthy individuals, we can infer to some degree that the consequence of that mutation is likely to be less severe than mutations not present in the reference sample. Comparing to the ~45K non-psychiatric samples from the ExAC reference database ("non-psych ExAC"; (Lek, et al. 2016)), we find that 26% of DNMs found in SCZ probands are observed in this dataset (Kosmicki, et al. 2017). Of note, while individuals the Southeast Asian ancestry are not well represented in the ExAC v1 database, the presence of extant alleles at DNM sites in any population lead to the same inference of likely reduced pathogenicity of the mutation.

When we re-evaluate the burden of DNMs between SCZ probands and controls after partitioning by presence in non-psych ExAC cohort, we see the enrichment in SCZ probands is confined to DNMs not present in ExAC (fold-enrichment=1.12, *p*=2e-3), with no enrichment among DNMs present in ExAC (fold-enrichment=0.99, *p*=0.73; **Supplementary Table 5: DNM burden partitioned by presence in non-psych ExAC cohort**). The increased enrichment is driven by PTV (fold-enrichment=1.25, *p*=0.03) and missense (1.12, *p* = 0.01) DNMs, while rates of synonymous DNMs remain consistent (OR=1.05, *p* = 0.52). Overall, the increased rate of PTV and missense DNMs support the hypothesis that DNMs conferring risk to SCZ are more likely at sites invariant in the larger population.

**Supplementary Table 5: DNM burden partitioned by presence in non-psych ExAC cohort**

| | SCZ rate | Control rate | Fold enrichment (*p*-value) | SCZ rate | Control rate | Fold enrichment (*p*-value) | SCZ rate | Control rate | Fold enrichment (*p*-value) |
|---|---|---|---|---|---|---|---|---|---|
| | Present in ExAC | | | Not present in ExAC | | | Total | | |
| All coding | 0.26 | 0.26 | 0.99 (0.91) | 0.73 | 0.66 | 1.12 (2e-3) | 0.99 | 0.92 | 1.08 (1e-3) |
| PTV | 0.01 | 0.02 | 0.58 (0.05) | 0.09 | 0.08 | 1.25 (0.03) | 0.10 | 0.09 | 1.13 (0.21) |
| Missense | 0.16 | 0.16 | 1.01 (0.94) | 0.48 | 0.43 | 1.12 (0.01) | 0.65 | 0.60 | 1.09 (0.03) |
| Synonymous | 0.08 | 0.08 | 1.05 (0.70) | 0.16 | 0.15 | 1.05 (0.52) | 0.24 | 0.23 | 1.05 (0.44) |

**Table legend**: DNM burden among 2541 SCZ probands compared to 2182 controls. DNMs were partitioned by the presence of an allelic site in the 45k non-psychiatric ExAC cohort.

**Comparison with other neurodevelopmental disorders**

To better understand the relative contribution of coding DNMs to SCZ, we compared SCZ probands against probands diagnosed with intellectual disability (ID), developmental delay (DD), and Autism (ASD; see supplementary section 14). We focused on the exome-wide burden of PTV DNMs, and the burden of missense DNMs among evolutionarily constrained genes (defined here as the inclusive combination of RVIS, missense constraint, and pLI > 0.99) – two categories where there is a significant enrichment in DNM burden across all disorders analyzed. Unlike the previous burden analysis, we include all 2772 SCZ probands and restrict our analysis to 17925 well-covered genes.

DNM burden in SCZ probands is markedly reduced relative to all other neurodevelopmental disorders among both PTVs exome-wide and missense DNM in evolutionarily constrained genes, indicating that the contribution of DNMs towards a SCZ diagnosis accounts for a much smaller fraction of samples than earlier onset neurodevelopmental disorders (**Figure 2: DNM burden by disease**). Notably, while the DNM signal in ASD falls between SCZ and ID/DD phenotypes, follow-up work on ASD probands has shown that many individuals with ASD carrying candidate DNMs also have lower IQ (Kosmicki, et al. 2017), and many genes implicated from DNMs in ASD probands are more strongly associated with ID/DD phenotypes. These findings suggest that this signal is less likely to reflect strictly ASD-associated symptoms, but rather the co-morbidity of ASD features that arise in the face of severe cognitive impairment. For SCZ, the enrichment could simply reflect a small fraction of SCZ patients co-morbid with other neurodevelopmental features. In fact, scholastic achievement scores among the PTV carriers in the Bulgarian SCZ trio cohort (Fromer, et al. 2014) were significantly lower than other Bulgarian SCZ probands, and low scorers in sustained attention and executive function in the Taiwanese cohort show a higher burden of PTV and missense DNMs (supplementary section 9).

**Section 11 – DNM burden in predicted damaging missense sites**

The functional consequences of missense coding changes vary widely depending on the gene affected and/or the amino acid perturbed, varying from no discernable phenotypic changes (benign) in many cases while showing severe consequences in others (damaging). Several *in silico* prediction tools exist to predict the likely consequence of a missense mutation, and among ultra-rare variants analyzed in SCZ case-control exomes, predicted damaging missense variants were enriched in SCZ cases whereas predicted non-damaging missense variants showed no enrichment (Genovese, et al. 2016). Missense DNMs were annotated using 11 prediction algorithms provided in dbNSFP v2.9 (Liu, et al. 2011, 2013), along with the combined damaging prediction used in the Swedish SCZ case-control cohort.

We tested for enrichment in predicted damaging missense DNMs in SCZ probands using a two-sample binomial test, which conditions on the overall DNM rate difference between SCZ probands and controls, thus allowing us to use the full sample set (2772 SCZ probands and 2216 controls). We looked at the full set of missense DNMs (**Supplementary Figure 15: Missense predictor enrichment**) and those not present in non-psych ExAC cohort (**Supplementary Figure 16: Non-ExAC missense predictor enrichment**). Polyphen2 HDIV, Polyphen2 HVAR, CADD, and SIFT all show nominal enrichment in SCZ probands ($p < .05$), however no single prediction algorithm surpasses multiple testing correction ($p < 4.5\text{e-}3$ for 11 tests). Overall, while predicted damaging DNMs show further enrichment in SCZ over missense changes overall, the effect is modest. We incorporate the top performing missense predictions when determining a "SCZ enriched" set in gene recurrence and gene set analyses, which we detail in supplementary section 13.

#### Section 12 – DNM burden in predicted functional synonymous sites

In general, synonymous changes to coding sequence are predicted to have minimal downstream consequences relative to nonsynonymous changes, as the amino acid codon is preserved. Furthermore, among neuropsychiatric disease cohorts, synonymous DNMs have often shown minimal, if any, DNM enrichment in affected probands. However, recent analyses have shown that synonymous sites predicted to affect or interact with regulatory elements do show enrichment in published Autism (ASD) and SCZ coding DNMs (Takata, et al. 2016). In both ASD and SCZ cohorts, synonymous DNMs closer to the splice site were more enriched in cases relative to controls, and within SCZ cohorts, synonymous DNMs falling within frontal cortex derived DNAse Hypersensitivity peaks.

We sought to replicate these findings in the Taiwanese cohort and independent controls, following closely to the annotations used in (Takata, et al. 2016). Distance to splice site was obtained using the SeattleSNPs annotation platform (http://pga.gs.washington.edu), with additional annotation of exonic splicing regulator (ESR) motifs (both enhancers and silencers) to refine sites likely to affect splicing (Fairbrother, et al. 2002; Ke, et al. 2011; Iossifov, et al. 2014). DNAse Hypersensitivity sites (DHS) were obtained from ENCODE tracks available in the UCSC browser (https://genome.ucsc.edu/ENCODE/dataMatrix/encodeDataMatrixHuman.html), and hg19 tracks of narrow peaks from cerebrum/frontal cortex tissue (CFC), frontal cortex tissue (FC), and cerebellum tissue were analyzed. We first tested the main SCZ findings in SCZ from (Takata, et al. 2016) for enrichment in two synonymous DNM annotations relative to all synonymous DNMs. The first is synonymous DNM enrichment within cerebrum/frontal cortex DHS peaks, and the second within 30 bp of a splice site and with an ESR motif (termed 30 bp NSS-ESR). For replication, we compared the 1695 Taiwanese probands against the 1485 controls not used in the primary analysis of (Takata, et al. 2016)**.**

##### Modest enrichment in functional synonymous DNMs in independent replication

Using a two-sample Poisson test, synonymous DNMs within frontal cortex DHS peaks are modestly enriched in Taiwanese probands (73 DNMs in 1695 trios) relative to independent controls (56 DNMs in 1485 trios; fold-enrichment=1.14, one-tailed $p$=0.26), as compared to the enrichment reported in the published set (fold-enrichment=2.58, one-tailed $p$=8e-4). Similarly, synonymous DNMs annotated as 30 bp NSS-ESR are modestly enrichment in Taiwanese probands (92 DNMs in 1695 trios) relative to the independent controls (64 DNMs in 1485 trios; fold-enrichment=1.26, one-tailed $p$=0.09), as compared to the enrichment reported in the published set (fold-enrichment=1.89, one-tailed $p$=0.006). While the replication results suggest that the initial finding may be a false positive finding, further analysis indicates that near splice site variants may in fact be enriched in SCZ probands.

##### Consistent enrichment in NSS-ESR synonymous DNMs within 60 bp

When we look across the range of distance to splice site using a two-sample proportion test, pattern suggests that enrichment in NSS-ESR among SCZ cases is most prominent within 60 bp (or 20 codons) of a splice site (combined fold-enrichment=1.29, $p$=5e-4). This pattern is consistent in both published SCZ and Taiwanese probands in relation to both previously reported and independently published controls (**Supplementary Figure 17: Synonymous DNM enrichment at NSS-ESR sites**), and suggests that near splice site synonymous DNM are contributing to SCZ risk. In fact, among the combined SCZ and controls, 60bp NSS-ESR sites are more significantly enriched than brain-derived DHS peaks and predicted deleteriousness measured using CADD annotation (**Supplementary Figure 18: Synonymous annotation enrichment**).

**Section 13 – SCZ enriched DNM subset**

Given that several additional annotation categories within missense and synonymous DNMs are significantly enriched SCZ probands over controls, we wanted to streamline these annotations into a single predictor for gene set and gene recurrence analysis. To achieve this result, we used a step-wise process of elimination to prioritize annotations that best contribute to the whole-exome burden signal. To start, we restricted to DNMs not present in ExAC, as we see no enrichment among DNMs present in ExAC (section 10). Within both missense and synonymous DNMs, we ranked each annotation by *p*-value, and iteratively removed the most significant predictors of SCZ enrichment until we saw a sign change in the DNM rate (i.e. until controls were more enriched than SCZ cases; **Supplementary Figure 19: SCZ enriched annotation**). We took all annotations that showed a nominally significant enrichment within SCZ cases over controls ($p < .05$). Of note, no annotation reach nominal significance within PTVs, and so all PTVs not present in ExAC were retained as 'SCZ enriched' (90% of all PTVs).

SCZ enriched missense DNMs include those falling within ENCODE DHS peaks from cerebrum, frontal cortex, and cerebellum tissue, as well as those with a high CADD v1.3 PHRED score (> 25). Missense DNMs annotated as SCZ enriched make up 57% of all missense DNMs. SCZ enriched synonymous DNMs include those falling within 60 bp of a splice site and occur within exon-splicing regulator motifs. Synonymous DNMs annotated as SCZ enriched make up 36% of all synonymous DNMs. In all, SCZ enriched DNMs make up 41.7% (1154 of 2769) of all DNMs in the combined SCZ cohort.

**Section 14 - DNM burden in candidate gene sets**

Gene set enrichment (also known as pathway analysis) aims to find if any group of genes, however classified, is enriched among SCZ affected individuals. Often, for DNM and ultra-rare variation, even quite large samples do not have enough statistical power to implicate individual genes as having SCZ associated risk. Gene sets with sufficient exome coverage, however, will provide enough variance in DNM counts to surpass strict exome-wide correction for multiple testing. Here we detail the procedure for determining multiple testing correction, assess the relative power in the current dataset to implicate candidate gene sets, and highlight gene sets most enriched among SCZ DNMs.

To test for gene set enrichment, we fit the data to a binomial model. This model is conditional on the overall DNM rate as it examines the relative proportions of overlap rather than the observed rate. For example, if SCZ probands had twice the rate of PTV DNMs than controls, but 50% fell in brain-expressed genes in each sample, then we would have two-fold enrichment in the PTV rate, but no proportion enrichment for brain-expressed genes among PTVs. For comparing against mutation model expectations, we used a one-sample exact binomial test, with the mutation expectation as our null probability of gene set overlap. For comparing against control DNMs, we used a two-sample exact binomial test. For each candidate gene set, we tested for enrichment among 1) all DNMs, 2) SCZ-enriched (section 13), 3) PTVs, 4) PTV and missense, 5) missense, and 6) synonymous DNMs.

**Gene set enrichment testing correction and power analysis**

When testing many gene sets across several DNM annotations, and with many gene sets having overlapping genes, we wanted to ensure that an accurate portrayal of the type I error was taken in to account. To correct for multiple testing in the current analysis, we simulated 1000 DNM lists in SCZ cases and controls using the mutation expectation model adjusted for coverage QC (supplementary section 8). In each simulation, probabilities of selecting a gene were based on their mutation expectation, and counts of PTV, missense, and synonymous DNMs were identical to those observed in SCZ and control cohorts. We then ran gene set enrichment on each simulated DNM list against 85 candidate gene sets (**Supplementary Spreadsheet 8: Candidate gene sets**), and retained the lowest *p*-value from six DNM annotations tested. The 5% family-wise error rate (FWER) is the 50$^{th}$ lowest p-value among the 1000 *p*-values, and represents the 5% type I error rate among the 85 candidate gene sets.

Among the six annotations tested, the 5% FWER was quite consistent (*p*-value range: 7.3e-4 to 1.1e-3). Further, FWER is consistent for one-sample (mutation model) and two-sample (SCZ vs control) tests. Collapsing all six tests together, the 5% FWER is *p*=8e-4. We use this *p*-value threshold moving forward for all candidate gene set tests, and as the criterion for achieving statistical power in our power analysis.

Statistical power in our gene set analysis was measured by what percent of gene sets could surpass the 5% FWER at full penetrance and at the highest observed fold-enrichment in the dataset. Among case-control tests, a fully penetrant gene set would need at least 18 DNMs to surpass the 5% FWER. Among gene sets with at least 18 overlapping DNMs in both cases and controls, the highest fold-enrichment observed is 2.57-fold, and we would need a minimum of 56 DNMs to reach FWER at this level of penetrance. Among candidate gene sets, 55.66% of tests consider at least 56 DNMs (39% of PTV tests). Under a scenario where we had 80% power to detect a 3-fold enrichment, we would need to observe at least 73 overlapping DNMs in both cases and controls (1.6% of all DNMs, or 15% of PTVs). Using these numbers, the smallest gene set with 73 overlapping DNMs has 93 genes, and with 73 overlapping PTVs, 863 genes.

Statistical power using the mutation model varies from the case-control comparison as it is a one-sample test against a fixed expectation. In this scenario, a fully penetrant gene set would need at least 9 DNMs to surpass the 5% FWER. Among gene sets with at least 9 overlapping DNMs in SCZ probands, the highest fold-enrichment observed is 5.3-fold, and we would need a minimum of 15 DNMs to reach FWER at this level of penetrance. Among candidate gene sets, 70% of tests consider at least 15 DNMs (57% of PTV tests).

While the mutation model is a more sensitive test than case-control enrichment tests, mis-specification of the model can lead to bias in the comparison. While we addressed this issue at the whole-exome in the mutation rate model testing (supplementary section 8), we are mindful that more subtle estimation errors in mutation expectation are still possible. To better organize our assessment of gene sets enriched for SCZ DNMs, we first examine our overall results both against the mutation model and comparing to control trio DNMs, then focus on enrichment in different gene set categories against only the mutation model, where we are better powered to assess smaller gene sets.

**Highly brain expressed and constrained genes are enriched for SCZ DNMs**

We tested the entire set of DNMs in SCZ probands against 85 candidate gene sets. We restricted our gene set analysis to the 17925 genes that meet sufficient exome coverage (supplementary section 8), and used permutation procedures to estimate our multiple testing correction cutoff ($p$ < 8e-4) at a 5% alpha level. Among all DNMs, only four independent gene sets are significantly enriched in both the mutation model and against controls, whereas a larger proportion of gene sets surpass multiple testing correction when compared to the mutation model, owing to the increased statistical power when testing against a theoretical model. Results from all candidate gene set are available in **Supplementary Spreadsheet 9: Candidate gene set results**.

Among the four gene sets that surpass multiple testing correction in both tests, two of these sets represent the broad suite of genes highly expressed across human brain tissues and cell types, as evidenced by expression patterns derived from post-mortem brain tissue (downloaded from http://www.brainspan.org/ and https://www.gtexportal.org/home/). The other two sets encompass genes intolerant to coding variation, whether it be relative to common functional variation (Residual Variation Intolerance Score or RVIS, (Petrovski, et al. 2013)**;** genic-intolerance.org), or relative to modeled expectations (missense constraint and loss-of-function intolerance (pLI); (Samocha, et al. 2014; Lek, et al. 2016)**;** exac.broadinstitute.org). These gene sets suggest that perturbations to a broad set of critically important brain-expressed genes contribute to SCZ risk, reinforcing the notion that the genetic risk for SCZ comprises a strongly polygenic architecture, even among the rarest class of genetic variation.

SCZ enriched DNMs determined from the overall case/control burden do not markedly improve the signal, but remain enriched for missense constrained/RVIS intolerant genes even after removing all highly expressed brain genes (fold-enrichment=2.03, $p$=0.026), while non-enriched DNMs remain enriched for highly brain expressed genes after removing missense constrained/RVIS intolerant genes (fold-enrichment=1.10, $p$=0.034; **Supplementary Figure: FWER passing gene set enrichment**). Overall, this broad pattern suggests that the SCZ risk conferred by DNMs is not confined to a specific cell type, predicted consequence, or property of a gene. Beyond these results, gene set enrichment among SCZ enriched DNMs does not produce much in the way of unique results relative to all DNMs, nor was it amenable to the DNM model comparisons, so we did not pursue this specific annotation any further.

**Highly constrained genes are enriched for SCZ DNMs**

We find consistent enrichment in gene sets defined by the observed depletion of rare functional variation among the ExAC reference cohort, whether is in depleted rare functional variation relative to common functional variation (RVIS), rare missense to synonymous deviation (missense constraint), or rare PTV to synonymous deviation (pLI). This enrichment is greatest among genes identified using multiple predictors, whereas genes with less evidence of constraint, such as genes with pLI score between 0.9 and 0.99, are not enriched **(Supplementary Figure 21: Evolutionary constraint gene set enrichment and Supplementary Figure 22: Evolutionary constraint gene set PTV enrichment)**. In all, these results make clear that DNMs in SCZ probands are enriched for genes under negative selection and likely to lead to severe fitness consequences when perturbed. This insight, however, does not point to any specific biological pathway, cell type, or function that uniquely gives rise to SCZ risk.

**No brain-specific cell type is uniquely enriched for SCZ DNMs**

Within our set of highly expressed brain genes, we wanted to see if a specific brain cell type was enriched for SCZ DNMs. Using gene sets derived from brain cell-type transcriptional profiling (Cahoy, et al. 2008), we focused on neuronal, oligodendrocyte, and astrocyte cell types. Overall, all classes of DNMs were enriched among genes highly expressed in all three cell types, with the smallest enrichment and largest discrepancy between model and control coming from missense DNMs. For tissue-specific gene expressed genes, we see more enrichment coming from neuronal and oligodendrocyte tissue than astrocytes, however this difference is not very substantial (**Supplementary Figure 23: Brain expression gene set enrichment**). Intriguingly, the strongest effect is seen among synonymous DNMs for genes specifically expressed in neurons (fold-enrichment=1.46, $p$=8e-4), further highlighting the notion that SCZ DNM risk is not restricted to nonsynonymous changes.

**Overlap of SCZ DNMs with other neurodevelopmental disorders**

An earlier study found a significant gene overlap between DNMs in SCZ probands and DNMs seen in probands with Autism or Intellectual Disability (Fromer, et al. 2014), suggesting an underlying gradient of shared neurodevelopmental pathology across risk genes. Exome DNM results have subsequently expanded in the past few years, and we compared our DNMs against those observed in Autism (3982 probands (Iossifov, et al. 2012; Neale, et al. 2012; O'Roak, et al. 2012; Sanders, et al. 2012; De Rubeis, et al. 2014; Iossifov, et al. 2014), Intellectual disability (971 probands, (de Ligt, et al. 2012; Rauch, et al. 2012; Lelieveld, et al. 2016), and Developmental Delay (4293 probands, (Deciphering Developmental Disorders 2017)). Reinforcing earlier findings among larger samples, we see enrichment across DNMs observed in all three diagnoses against the mutation model (Autism PTV/Missense fold-enrichment=1.14, *p*=3e-5; Intellectual disability PTV/Missense fold-enrichment=1.18, *p*=4.6e-3; Developmental Delay PTV/Missense fold-enrichment=1.1, *p*=9e-4). When we split the gene sets into those shared across multiple neurodevelopmental disorders and those specific to each, only the shared genes retain significant enrichment (**Supplementary Figure 24: Neurodevelopmental disease gene set enrichment**). This finding reinforces the hypothesis that perturbed genes conferring risk for neuropsychiatric disease are likely to be penetrant across diagnostic criteria.

**Deviation of model and case control enrichment in gene set analysis**

In contrast to constrained and brain expressed genes, the signal we see among potentially synaptic genes and within neurodevelopmental disorder gen sets is non-significant when comparing against control trios while significant in the mutation model. Despite the effort used to better calibrate the mutation model to the uneven coverage distribution of exome sequencing (supplementary section 8), there may remain uncorrected bias in the model, particularly among gene sets ascertained from exome sequencing studies. Along with model bias, this may also simply be a lack of statistical power in the case-control analysis relative to the mutation model, or even true enrichment of DNMs in these genes among controls despite no diagnosis of neuropsychiatric disease (most of the control trios are unaffected siblings of ASD probands). To better assess what may be driving the discrepancy, we increased the exome coverage requirement for gene inclusion to 100% at 10x coverage (from 75% at 10x coverage) to be retained in the analysis. This dropped the number of genes examined from 17,925 to 12,311, with the retained genes being more robustly covered by sequence reads. More deviation between case-control and mutation model results at better coverage would suggest that coverage bias is driving the discrepancy between case-control and model results, while no change or better concordance would suggest that either a lack of statistical power or enrichment in controls better explains the discrepancy in our initial results.

For potentially synaptic genes, restricting to genes with better coverage does show more concordance of the case-control enrichment with the initial mutation model signal, however the lack of statistical power at these effect sizes still has the case-control comparison as non-significant (**Supplementary Figure 25: Synaptic gene set enrichment by exome coverage**). In contrast, gene sets derived from exome sequencing in neurodevelopmental disorders now show less enrichment in case-control results at better coverage (**Supplementary Figure 26: Neurodevelopmental disease gene set enrichment by exome coverage**). For both sets, however, the mutation model results show significantly more enrichment when restricting to better coverage genes. This finding was not expected from simulation results, which predicted a depletion from expectation, not an enrichment, for coverage biased gene sets when restricting to better covered genes. This result may reflect that our use of 10x sequencing depth as the means of coverage calculation may be insufficient, and coverage bias driven by much deeper covered genes is amplified more readily among the smaller exome target.

When we divide our gene set enrichment results in the neurodevelopmental gene sets by DNM annotation, the discrepancy is strongest among missense and synonymous DNMs, as it remains concordant among PTV DNMs, suggesting that much of the residual bias is not affecting PTV DNMs. This may be in part because PTVs in neurodevelopmental disorders are much more penetrant for disease, but they also make up a smaller fraction of the coding DNM landscape. The miscalibration of the model in missense and synonymous DNMs suggests that the ascertainment of the neurodevelopmental disease gene set enriches for large, very well covered genes that are likely to show DNM hits in multiple exome sequencing studies despite no meaningful association with neurodevelopmental disorder. Overall, these comparisons suggest that sequence coverage plays an integral role in driving the overlap of DNM hits, and for gene sets derived from exome sequence studies, presents a potential confound that needs to be carefully controlled for in mutation model estimates to provide accurate results. Thus, the inclusion of case-control results is an important check on the validity of mutation model assumptions.

**Section 15 - DNM burden in GO and SynaptomeDB databases**

UniProt human GO term annotations were downloaded from the February 16, 2016 distribution at http://geneontology.org/page/download-annotations. We removed gene entries including uncharacterized proteins, proteins with unknown genes, had redundant gene symbols in the same GO term, or contained a 'NOT' qualifier. This resulted in 16276 gene sets, 765 of which had at least 50 genes. Gene sets downloaded from SynaptomeDB (http://metamoodics.org/SynaptomeDB/index.php) represent a compilation of 488 GO, KEGG, BIOCARTA, REACTOME, and TFT/MOTIF annotations related to synaptic biology and function, 147 of which had at least 50 genes. By restricting our analysis to only gene sets with at least 50 genes in GO and SynaptomeDB, we analyzed a total of 911 gene sets.

We ran the same permutation procedure as the candidate gene set analysis (supplementary section 14) to estimate our multiple testing correction cutoff at a 5% alpha level (cutoff $p$=6e-5 for the mutation model test and cutoff $p$=2e-4 for the case/control test). No single gene set surpassed multiple testing correction in either comparison. Our top gene sets in both tests are listed in **Supplementary Spreadsheet 10: GO+SynaptomeDB enrichment** and the QQ plot of $p$-values for both comparisons shown in **Supplementary Figure 27: GO/SynaptomeDB gene set model enrichment** and **Supplementary Figure 28: GO/SynaptomeDB gene set case-control enrichment**. When comparing against the mutation model, the most significant gene set is among all DNMs overlapping genes involved the neurotransmitter secretion biological process (GO:0007269, 64 genes tested, fold-enrichment=2.12, $p$=4e-4). Among PTVs, the most significant enrichment overlaps genes involved in chromatin organization biological process (GO:0006325, 207 genes tested, fold-enrichment=2.81, $p$=4e-4;). When comparing against controls, the most significant gene set is among all DNMs overlapping genes making up the biosynthetic process in Synaptome DB (73 genes tested, $p$ = 3e-4, 15.8 fold-enrichment).

Among the top gene sets, neurotransmitter secretion highlights a specific biological process, whereas chromatin organization and biosynthesis make up a broader suite of core biological processes. Neurotransmitter secretion is described as "the regulated release of neurotransmitter from the presynapse into the synaptic cleft via calcium-regulated exocytosis during synaptic transmission." While not surpassing correction for multiple testing, the suggestive association with neurotransmitter secretion highlights the perturbation of pre-synaptic activity as a risk factor for SCZ, while the enrichment in chromatin organization genes further support the role of chromatin processes in SCZ risk, as they have been previously implicated for SCZ risk in common variant GWAS (Schizophrenia Working Group of the Psychiatric Genomics 2014), TWAS (Gusev, et al. 2018) and rare variant scans (McCarthy, et al. 2014; Singh, et al. 2017), and among autism DNMs (De Rubeis, et al. 2014).

**Section 16 – Single gene DNM enrichment**

To see if any single gene was a putative risk factor for SCZ, we tested for per-gene enrichment of DNMs in the combined SCZ cohort. Enrichment is measured using a one-sided Poisson test against the gene level mutation expectation. Where applicable, gene level mutation expectations adjusted by exome coverage were used (supplementary section 8), otherwise expectations from canonical GENCODE transcripts were used, which assume full transcript coverage. Expected rates are the annotation-specific mutation rate multiplied by the number of inherited chromosomes.

We tested three annotation categories: 1) PTV, 2) PTV and missense, and 3) All coding DNMs (PTV, missense, and synonymous). Exome-wide significance threshold was set at $p < 8.7\text{e-}7$ to correct for multiple testing (5% alpha for 19164 genes x 3 tests). No gene surpassed exome-wide significance, with our lowest *p*-value across all three tests being 7.7e-6, which doesn't surpass exome-wide significance even if only single test was considered.

Among PTVs, the most significant gene is SET Domain Containing 1A (*SETD1A*), with 3 PTVs observed in two previously published SCZ trio cohorts, ((Guipponi, et al. 2014; Takata, et al. 2014), pLI=1, *p*=7.7e-6). *SETD1A* has since been identified as an exome-wide significant gene association in combined trio and case/control exome sequencing of SCZ, whereby follow-up of patients carrying a PTV in *SETD1A* often presented with an associated neurodevelopmental disorder (Singh, et al. 2016). Among the combined missense and PTV test, the most significant signal is in the TATA-Box Binding Protein Associated Factor, RNA Polymerase I Subunit C (*TAF1C*), which has four missense variants coming from two separate cohorts (pLI=0, *p*=6e-5). *TAF1C* is involved in RNA polymerase activity in the ribosome, and while the gene does not show evidence of constraint, it is present in many of the high brain expressed gene sets and all four missense variants are listed as 'probably damaging' in PolyPhen2 HDIV predictions.

Among all coding DNMs, the top signal resides in the pogo transposable element with ZNF domain (*POGZ*) gene, with five DNMs (one PTV, two missense, and two synonymous DNMs) across three separate cohorts (pLI=1, *p*=3.7e-5). Notably, both synonymous mutations occur within 60 bp of the splice site junction. POGZ is a highly constrained gene that has been associated with White-Sutton Syndrome, a form of intellectual disability, and is estimated to be responsible for 0.14% of individuals diagnosed with either autism or intellectual disability (Stessman, et al. 2016).

**Section 17 – Gene recurrence enrichment analysis**

While no single gene surpasses exome-wide correction, many genes are "recurrently" hit by a coding DNM (i.e. more than one DNM hits the gene among SCZ probands). The rate of recurrently hit genes can indicate how likely it is that such genes are indeed risk factors for SCZ. To test for an enrichment in recurrently hit genes, we compared SCZ probands against simulated DNMs using the mutation model (supplementary section 8). We performed the same test using control DNMs to ensure that any significant results were not the result of mis-specification in the mutation model. Of note, individuals with multiple DNMs in the same gene were restricted to only the single DNM with the most severe consequence (supplementary section 3), and will not bias the outcome of the recurrence test.

To estimate recurrence rates in the mutation expectation, we used a bootstrap re-sampling method to simulate an equivalent count of observed DNMs hitting genes, with the probability of selecting a DNM being its mutation expectation. For example, given we observe 296 PTVs in well-covered genes in SCZ probands, we simulate 296 DNMs using PTV probabilities from the mutation model, and then count how many DNMs fall into recurrently hit genes. For each test, we ran 100000 simulations, retaining the number of DNMs found recurrently hit genes for each iteration. Empirical *p*-values were obtained comparing our observed proportion of DNMs falling in recurrently hit genes to the distribution of simulated probability of gene recurrence.

We restricted to the 17925 genes meeting sufficient exome coverage (supplementary section 8). We tested gene recurrence probabilities among 1) All DNM, 2) PTV, 3) Missense, and 4) Synonymous DNMs. We also split the analysis by genes within the two most significant gene sets – highly brain expressed (10376 genes) and constrained (4083 genes) gene sets – to see if the overall enrichment in gene recurrence was confined to these categories of genes. When we compare our observed rates of recurrent genes against simulated DNMs, genes with recurrent PTVs are significantly enriched above expectation (16 genes, fold-enrichment=3.15, $p$=3e-5), a result not seen in controls (4 genes, fold-enrichment=1.5, $p$=0.27). While we see modest enrichment in genes recurrently hit by missense and/or synonymous DNMs, the enrichment is similar in controls (**Table 1: Gene recurrence rates in SCZ probands and controls**), and suggests deviations of the mutation model to the subtle effects of exome coverage and QC parameters. A full list of recurrent PTV genes is available in **Supplementary Spreadsheet 11: Recurrent PTV genes** and all recurrent PTV/Missense genes in **Supplementary Spreadsheet 12: Recurrent PTV+Missense genes**.

**Recurrently hit genes more likely to have high brain expression**

Given that high brain expression and constraint were consistently enriched in the gene set analysis (supplementary section 14), we wanted to see if recurrently hit genes were more likely to fall within these categories as well. Here, we define high brain expression as the inclusive combination of the BrainSpan high brain-expressed gene set and the GTEx brain enriched gene set, which encompass 10376 of the 17925 well-covered genes. We define constraint as the inclusive combination of RVIS intolerant, missense constraint, and pLI > 0.9 gene sets, totaling

4083 of the 17925 well-covered genes. The full set of comparisons are available in **Supplementary Spreadsheet 13: Gene recurrence by gene set**.

When we split the PTV signal, we do see a stronger enrichment within brain expressed genes (14 recurrently hit genes, fold-enrichment=3.25, *p*=5.6e-5) than outside (2 recurrently hit genes, fold-enrichment=2.02, *p*=0.26), whereas enrichment is smaller within constraint genes (8 recurrently hit genes, fold-enrichment=2.23, *p*=0.02) than outside constrained genes (8 recurrently hit genes, fold-enrichment=3.96, *p*=1.1e-3). When we consider other DNM annotations, we see enrichment among brain expressed genes that are not seen in the constraint lists considered (7019 genes; **Supplementary Table 6: Gene recurrence rates in highly brain expressed genes not under constraint**). The enrichment is consistent across all annotations in SCZ probands, whereas no significant enrichment is seen among controls. This finding suggests that genes highly expressed in the brain, when perturbed, can be risk factors for SCZ despite not showing evidence of strong selective pressure in the coding region.

**Executive function and sustained attention scores of Taiwanese trio probands with recurrent PTV genes**

SCZ patients carrying a PTV in highly constrained genes often show co-morbid intellectual disability (Singh, et al. 2017). To see if the genes with recurrent PTVs in SCZ probands exhibit a substantial effect on cognitive capabilities, we examined where carriers ranked on measures of sustained attention (1306 probands) and executive function (1325 probands) from the Taiwanese cohort (supplementary section 9). Among probands with measures of sustained attention, eighteen probands carried a PTV in one of the recurrently hit genes (**Supplementary Figure 29: CPT scores of recurrent PTV gene carriers in Taiwanese cohort**). Eleven of the eighteen had z-scores below the sample median, with PTV carriers in *TRIO* and *CHD8* scoring in the lowest $10^{th}$ percentile. Both genes are highly constrained and have been previously associated as DNM risk factors for ASD and intellectual disability in PTV carriers. Among probands with measures of executive function, sixteen probands carried a PTV in one of the recurrently hit genes (**Supplementary Figure 30: WCST scores of recurrent PTV gene carriers in Taiwanese cohort**). Ten of the sixteen had z-scores below the sample median, with PTV carriers in *TTN* and *HIVEP3* scoring in the lowest $10^{th}$ percentile. Interestingly, the proband carrying a PTV in *CHD8* scored in the $80^{th}$ percentile, showing a stark difference between sustained attention and executive function scores in this individual. Given the small sample size of PTV carriers with recurrent genes in the Taiwanese cohort, we are not adequately powered to make a strong inference about the impact of these specific PTVs on neurocognitive assessments. However, the distribution of scores suggests that PTV carriers in recurrently hit genes so far are not all pre-disposed to have co-morbid cognitive impairment.

**Supplementary Table 6: Gene recurrence rates in highly brain expressed genes not under constraint**

| | DNM | Observed recurrent genes | Expected recurrent genes | Fold-enrichment | Empirical *p*-value |
|---|---|---|---|---|---|
| 2772 SCZ probands | | | | | |
| All DNM | 854 | 92 | 73.6 | 1.25 | 6e-3 |
| PTV | 99 | 6 | 1.2 | 4.82 | 1e-3 |
| Missense | 546 | 43 | 31.9 | 1.35 | 0.02 |
| Synonymous | 219 | 10 | 5.7 | 1.75 | 0.06 |
| 2216 controls | | | | | |
| All DNM | 622 | 42 | 40.5 | 1.04 | 0.42 |
| PTV | 70 | 0 | 0.62 | 0 | 1 |
| Missense | 402 | 20 | 18.0 | 1.11 | 0.34 |
| Synonymous | 150 | 1 | 2.7 | 0.37 | 0.94 |

**Section 18 – Ultra-rare variant transmission among recurrently hit genes**

To further dissect the contribution of recurrently hit genes in SCZ probands, we examined the contribution of rare transmitted variation within these genes. While the lack of a SCZ diagnosis in the parents does lower the expected contribution that rare inherited variants play in proband risk, the polygenic nature of SCZ risk predicts that undiagnosed individuals carry SCZ risk alleles.

We used parent-proband transmission counts in the full Taiwanese cohort (1695 trios). Variants were filtered to sites with a minimum depth of 10x and with heterozygous calls having an alternate allele fraction of at least 30% in both parent and proband. Restricting to PASS sites in GATKs Variant Qualty Recalibration Score (VQSR) showed significant over-transmission, so to ensure that we re-capitulated the null expectation of equal transmission for rare variation, we followed the procedure in (Kosmicki, et al. 2017) that relaxed the VQSR inclusion parameter until we observed an equal transmission of synonymous parental singletons (**Supplementary Figure 31: Synonymous transmission by variant quality score**). For the transmission analysis presented here, we also filtered to ultra-rare variants, which are parental singletons in the current dataset that are also not seen in the non-psychiatric ExAC cohort (supplementary section 5). We used the transmission disequilibrium test (TDT**;** (Spielman, et al. 1993)) to determine the significance of over/under-transmission of parental singletons to SCZ probands.

When considering recurrently hit genes, there are a small number of very large genes where the observation of multiple DNMs is expected by chance. For example, mutation model expectations estimate 0.77 PTV DNM to overlap the gene Titin (TTN) in SCZ probands, so the observation of 2 DNMs ($p$=0.18) is non-significant. To this end, we removed any recurrently hit gene with a Poisson test $p$-value > 0.05.

TDT tests among recurrently hit PTV genes shows a modestly significant enrichment in ultra-rare PTVs (**Supplementary Table 7: Ultra-rare variant transmission rates in recurrently hit genes**), with a 3.25-fold enriched ratio of transmitted PTVs to non-transmitted PTVs. The contribution of transmitted PTVs comes largely from two genes, the Trio Rho Guanine Nucleotide Exchange Factor gene (TRIO; 4 transmitted ultra-rare PTVs and 0 non-transmitted ultra-rare PTVs), and the Dynein Axonemal Heavy Chain 9 gene (DNAH9, 6 transmitted ultra-rare PTVs and 1 non-transmitted ultra-rare PTV). TRIO also shows a suggestive over-transmission among ultra-rare predicted damaging missense variants (defined here as 'probably damaging' in PolyPhen2, 'deleterious' in SIFT, and not seen in ExAC; 11 transmitted to 2 non-transmitted, $p$=0.01), which is not seen in DNAH9 (8 transmitted to 7 non-transmitted, $p$=0.8; **Table 2: Genes recurrently hit by PTV DNMs**). When we examine the larger set of recurrently hit genes, including missense and synonymous DNMs, we see a depletion of transmitted ultra-rare PTVs, which is unexpected, as this is not reflected in the exome-wide list of well covered genes. When we consider ultra-rare predicted damaging missense transmission, we do not see much enrichment in recurrently PTV gene set (1.09-fold enriched ratio, $p$=0.64), and this modest level of enrichment persists across recurrently hit genes more broadly (**Supplementary Table 7: Ultra-rare variant transmission rates in recurrently hit genes**).

In general, the results suggest that our most definitive list of SCZ risk genes, namely genes recurrently hit by PTV DNMs, is nominally enriched for inherited rare variation. More inclusive annotations to define recurrently hit DNM genes and rare transmitted variants, however, do not show this signal.

**Supplementary table 7: Ultra-rare variant transmission rates in recurrently hit genes**

| Gene set | Genes | transmitted | non-transmitted | ratio | *p* |
|---|---|---|---|---|---|
| | Ultra-rare PTV | | | | |
| Recurrent PTV | 15 | 14 | 3 | 3.25 | 0.02 |
| Recurrent PTV or missense | 203 | 131 | 160 | 0.82 | 0.09 |
| Recurrent coding DNM | 316 | 185 | 235 | 0.79 | 0.01 |
| All well-covered genes | 17925 | 5866 | 5762 | 1.02 | 0.33 |
| | Ultra-rare predicted damaging missense | | | | |
| Recurrent PTV | 15 | 59 | 54 | 1.09 | 0.64 |
| Recurrent PTV or missense | 203 | 818 | 746 | 1.10 | 0.07 |
| Recurrent coding DNM | 316 | 1177 | 1098 | 1.07 | 0.10 |
| All well-covered genes | 17925 | 21892 | 21271 | 1.03 | 3e-3 |

#### Section 19 – Contribution of coding DNM to SCZ risk

When we consider our top findings from the gene set analysis, we can estimate the proportion of coding DNMs that contribute to a SCZ diagnosis. We combined the four gene sets that surpassed FWER correction, namely highly brain expressed genes from BrainSpan and GTEx resources, and constrained/intolerant genes from the top end of RVIS and missense constraint measurements. This list considers 10664 well-covered genes in all, and we find that 69.3% of coding DNMs fall within these genes in SCZ probands, while only 62.1% are overlap in controls, suggesting that 7.2% of coding DNMs contribute to a SCZ diagnosis. When we project this estimate as a Poisson distributed rate parameter, we estimate that around 195 of the 2772 SCZ probands carry a DNM that contributes to their SCZ diagnosis. It is important to keep in mind that the ascertainment of all SCZ trios are due to both parents being undiagnosed for SCZ, and represents an upper end of the contribution that DNMs have on SCZ risk.

##### Gene set comparison with rare variation in case-control SCZ studies

The recent analysis of a large Swedish SCZ cohort (Genovese, et al. 2016) examined 4.9k SCZ cases and 12.3k controls, and showed similar enrichment in many of the gene sets analyzed here. To further quantify the consistency of gene set enrichment, we compared fourteen gene sets examined in both SCZ case-control and trio studies. We contrasted the reported odds ratios from (Genovese, et al. 2016) against the proportion enrichment for PTV and missense DNMs among SCZ probands relative to the mutation model (**Supplementary Figure 32: Gene set model comparison with SCZ case-control exomes**). In only one of the gene set comparisons did we see a depletion in DNMs (91 SCZ GWAS genes), and for ten of the fourteen gene sets analyzed, the SCZ case-control dURV odds ratio fell between the PTV/missense and PTV-only fold-enrichment in SCZ probands, suggesting a marked consistency in effect size enrichment for these gene sets. We also compared dURVs against the trio case-control gene set enrichment in SCZ-enriched DNMs (**Supplementary Figure 33: Gene set case-control comparison with SCZ case-control exomes**), seeing a similar pattern of consistency to those the mutation model. Overall, these results indicate that a similar pattern of polygenic burden emerges across study designs, and for rare variant analysis in SCZ, both case-control and trio based studies are likely to reveal a similar genetic signature of rare variation conferring risk for SCZ.

#### Supplementary Figures

#### Supplementary Figure 1: SCZ vs control synonymous DNM rate

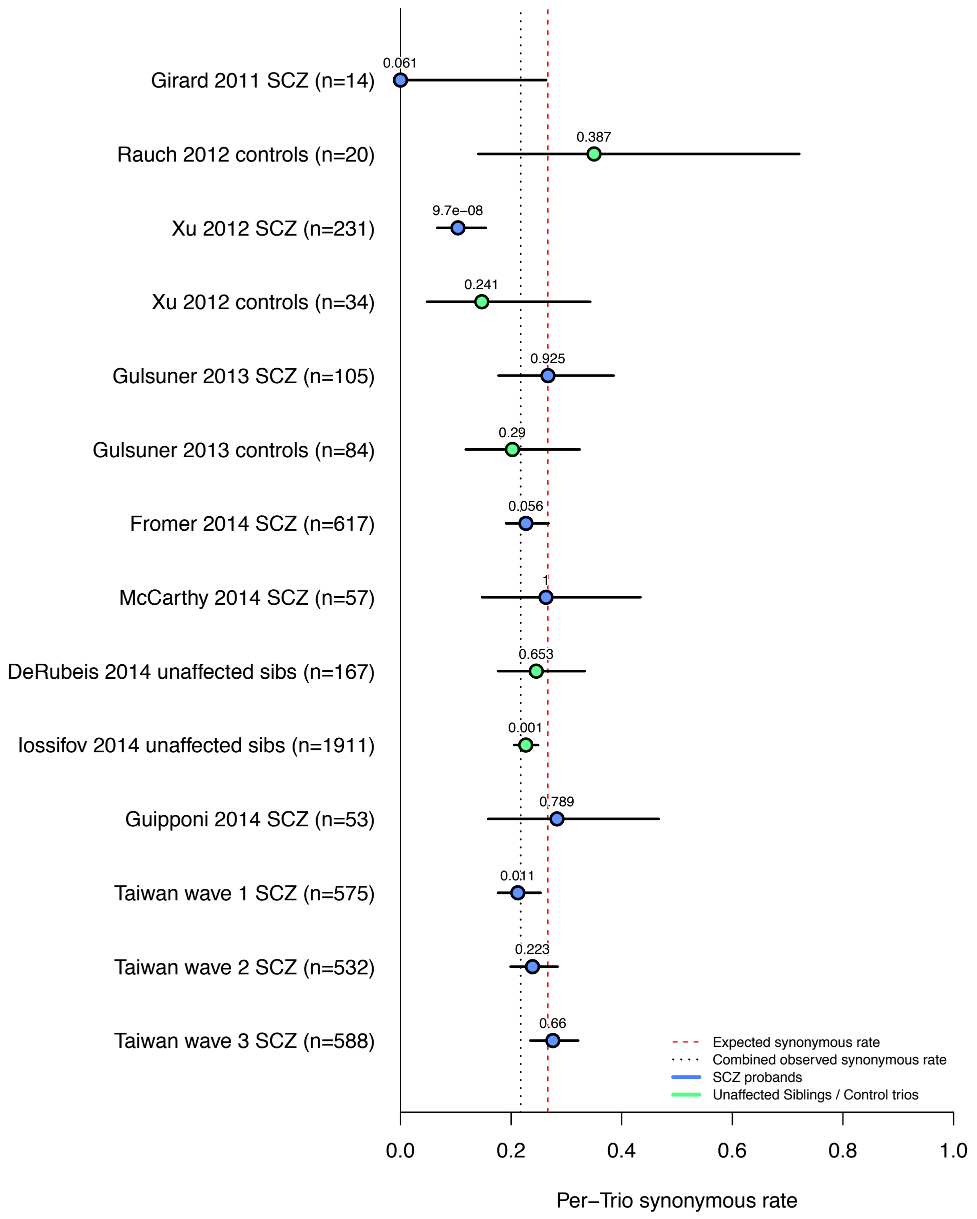


**Figure legend:** Per-trio synonymous DNM rate by study and case/control classification. DNMs were restricted to validated and published DNMs falling within hg19 Agilent v2 target intervals.

**Supplementary Figure 2: Taiwanese vs published SCZ synonymous DNM rate**

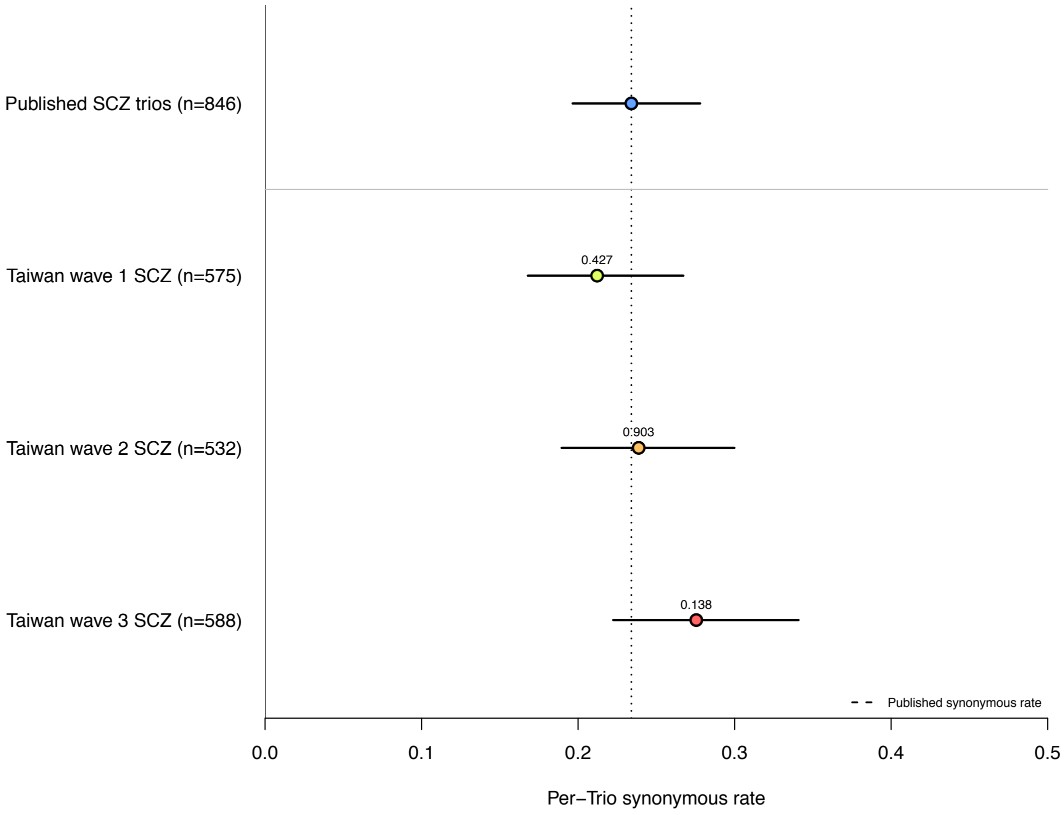


**Figure legend:** Per-trio synonymous DNM rate split by published SCZ trio studies and Taiwanese trio cohorts. Two-sample Poisson test *p*-values compare each Taiwanese trio cohort to the published rate. DNMs were restricted to validated and published DNMs falling within hg19 Agilent v2 target intervals.

#### Supplementary Figure 3: Mutation model coverage simulation

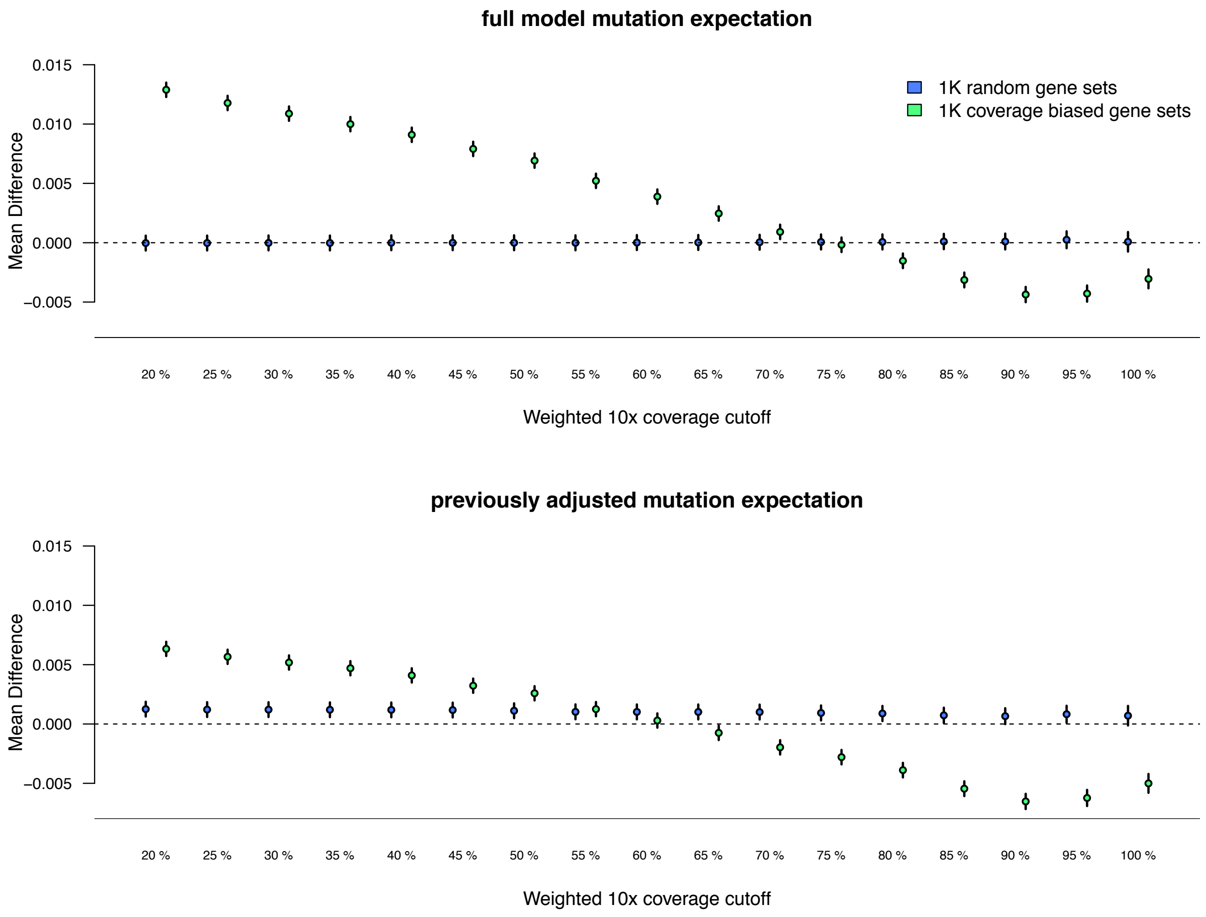


**Figure legend:** Mean difference between observed and expected gene overlap between selected gene sets and mutation model expectation(y-axis). Weighted 10x coverage cutoff retains the genes that meet the specified percentage of the canonical coding region at 10x coverage in the Bulgarian and Taiwanese trio cohorts (y-axis). Top graph uses the mutation rates without any previous coverage adjustment, where bottom graph uses previously published coverage adjustment criteria.

**Supplementary Figure 4: mutation model gene set size simulation**

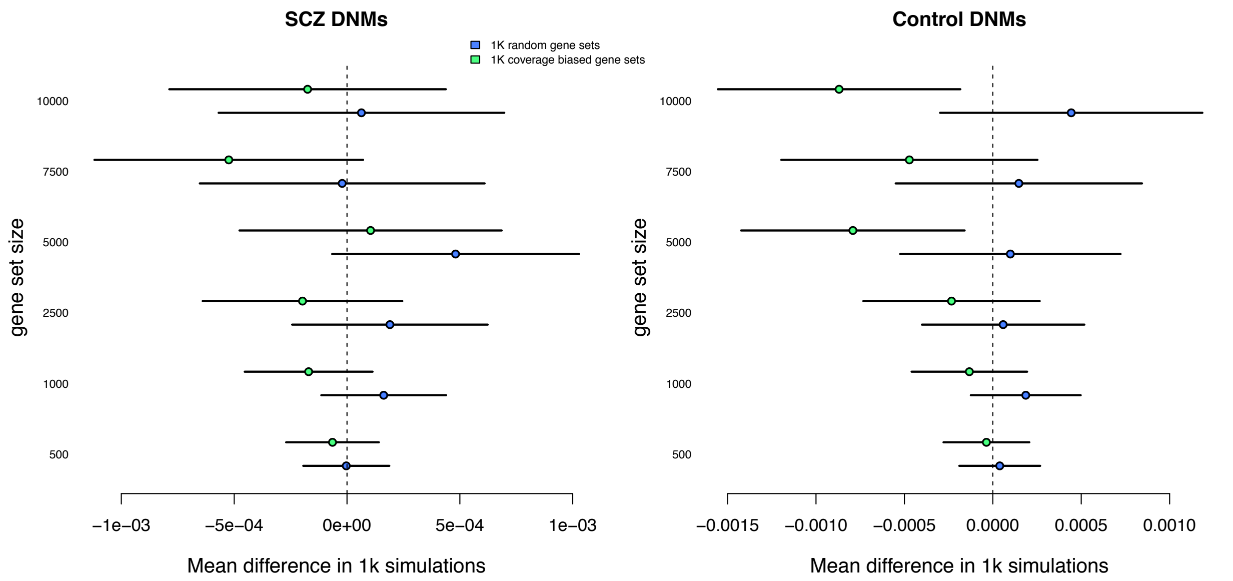


**Figure legend:** Mean difference in various gene set sizes when using the selected coverage adjustment (75% of gene at 10x) from the simulation model. The model was optimized using the 10k gene set in SCZ DNMs (top coordinates of left graph). Selected coverage adjustment was also tested against control DNMs with various gene set size simulations (right graph). Note that control exome coverage or DNMs were not used to select the best fit model.

**Supplementary Figure 5: Exome wide coverage adjustment**

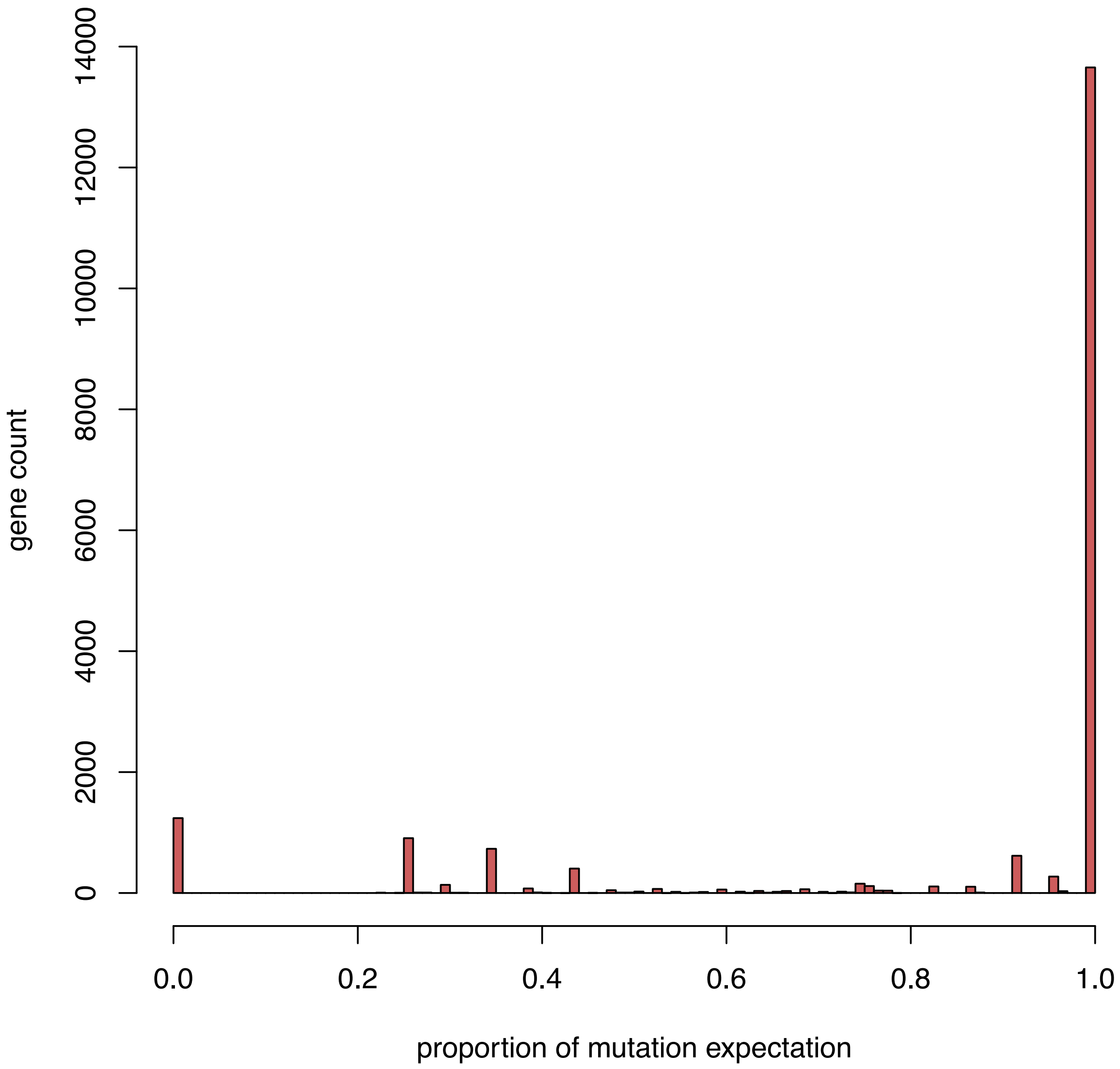


**Figure legend:** Mutation expectation adjustment in the selected 75% coverage adjustment model. Genes with less than 20% of their coding region meeting coverage were removed from the model (1241 genes or 6.5% of modeled genes). In contrast, 13,655 (71%) genes met 75% coverage in their entire coding region, and no adjustment was made to the mutation expectation.

**Supplementary Figure 6: Exome-wide burden in Agilent capture**

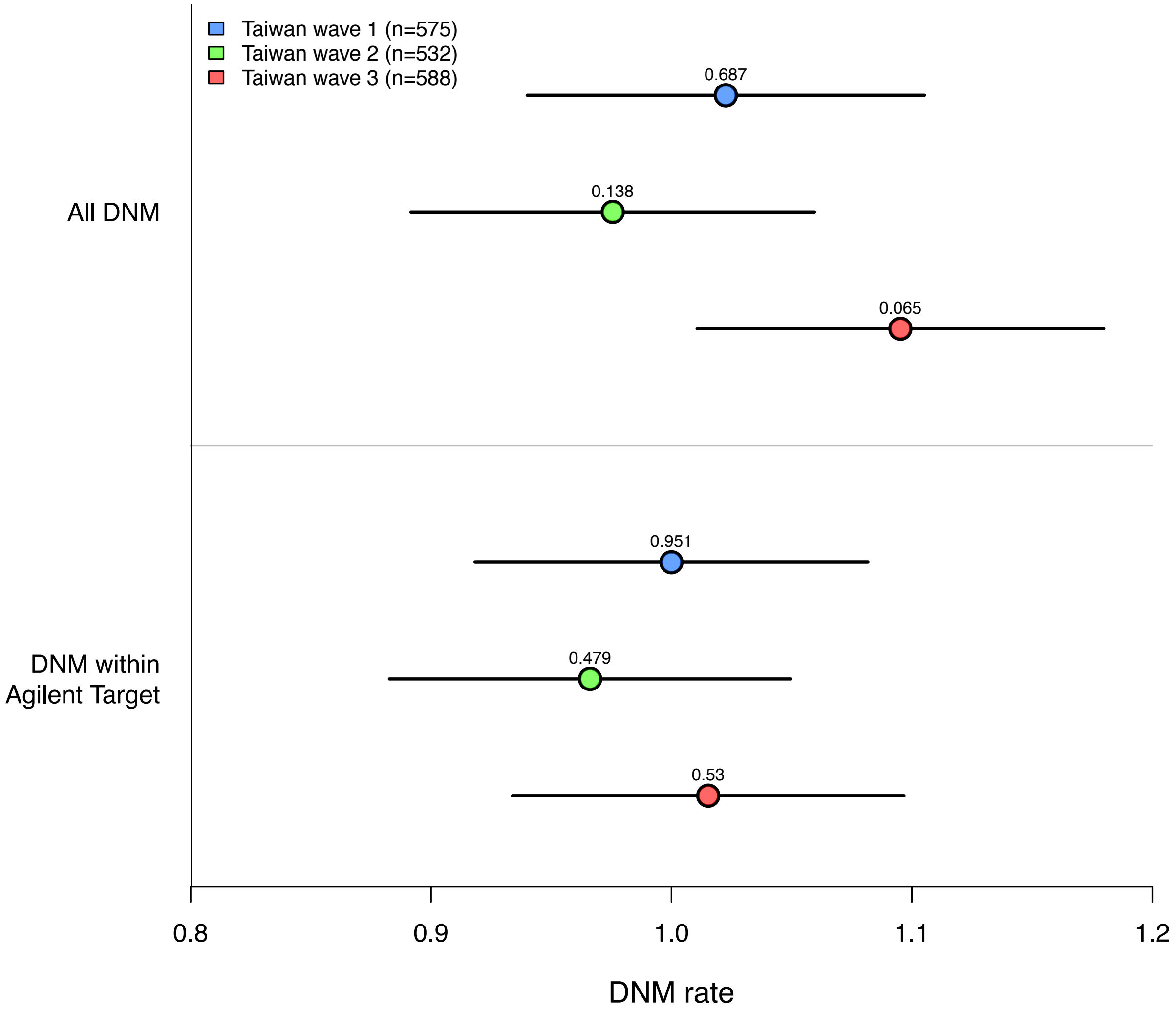


**Figure legend:** Overall DNM rate in Taiwanese trio sequencing waves before (top) and after (bottom) restricting to Agilent Sure Select v2 exome capture intervals. Two-sample Poisson rate *p*-values test the DNM rate in the selected cohort compared to the DNM rate in the other two cohorts.

**Supplementary Figure 7: Parental age on DNM burden**

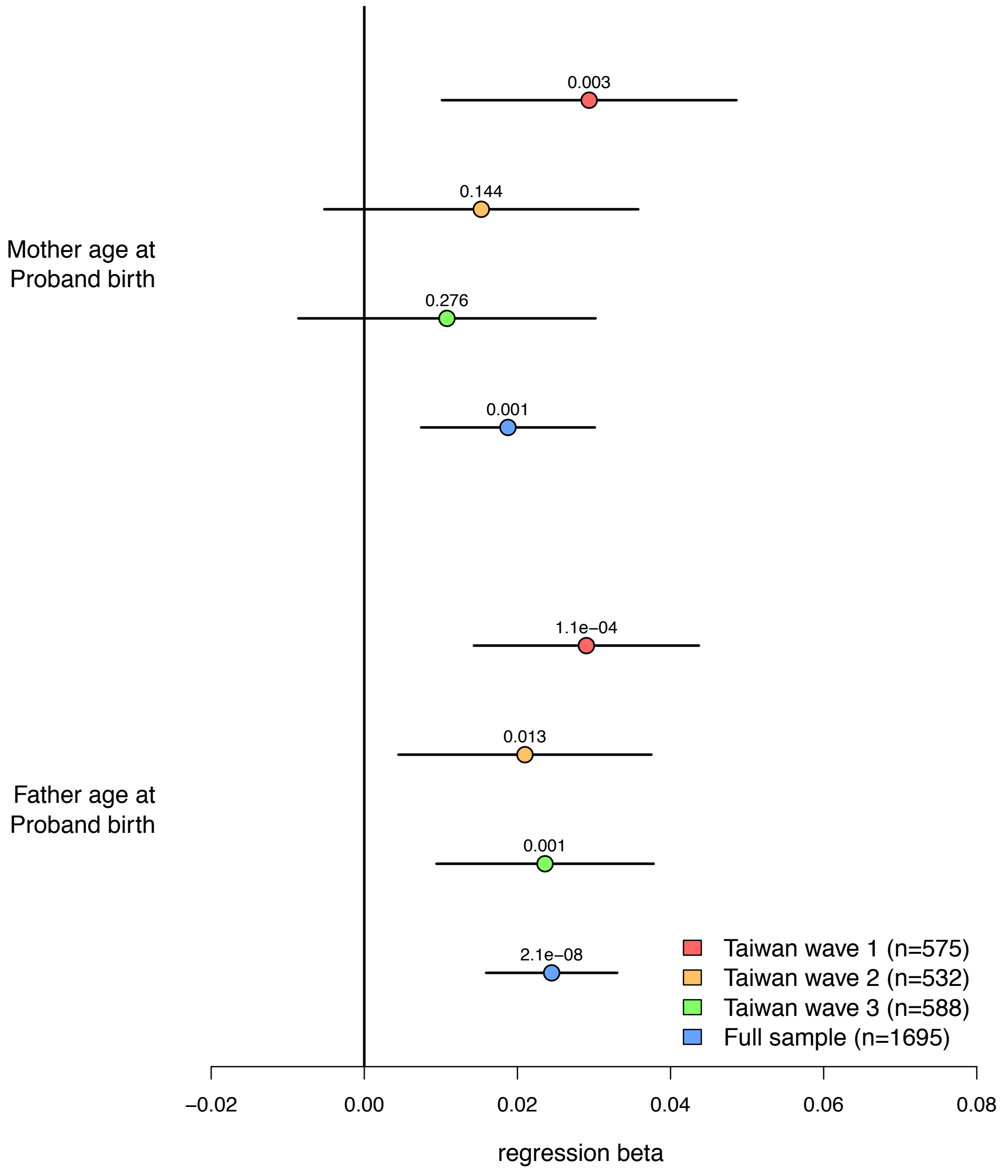


**Figure legend:** Effect of mother age (top) and father age (bottom) at birth on DNM rate among Taiwanese trio sequencing waves and full cohort. Results pictured are from a bivariate regression model with no additional covariates.

#### Supplementary Figure 8: Participant sex on autosomal DNM burden

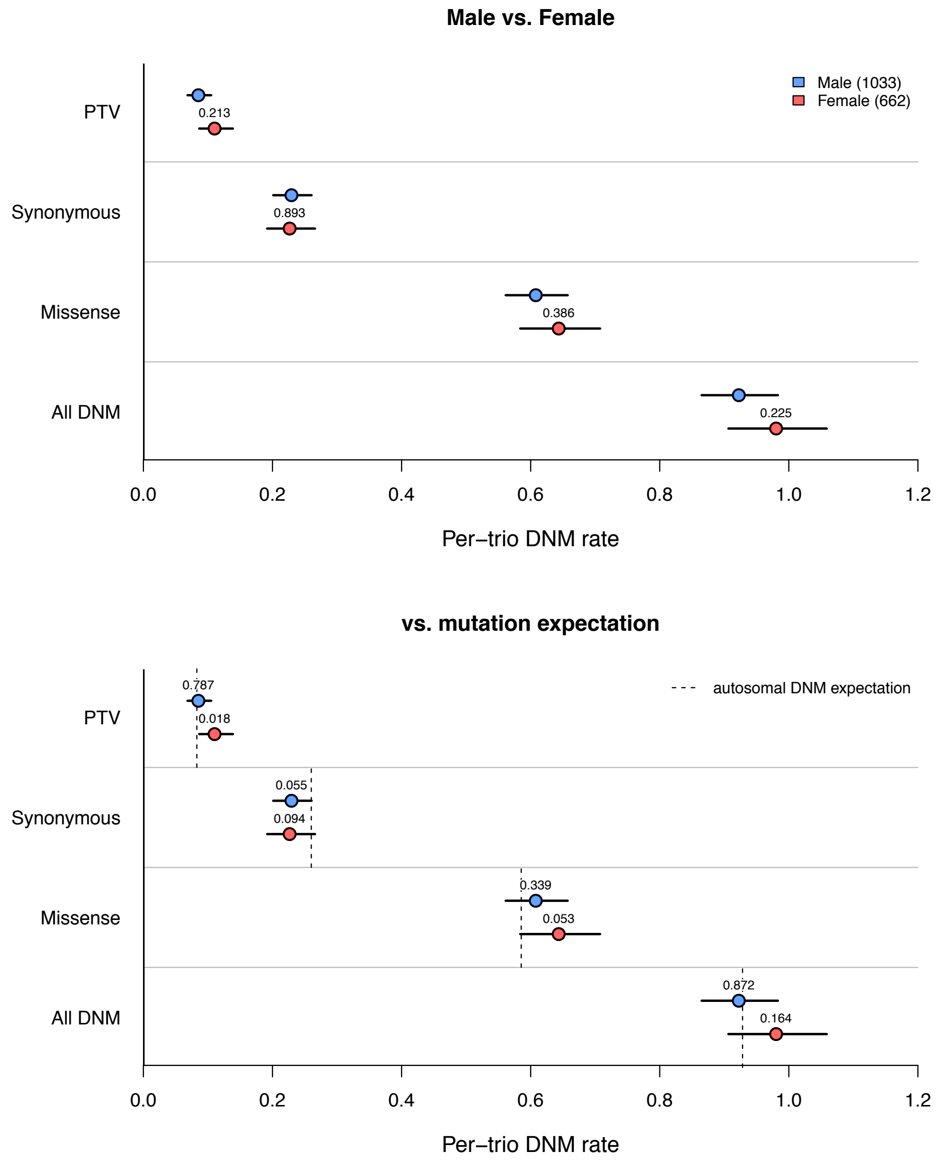


**Figure legend:** Autosomal DNM rate in Taiwanese trios split by proband sex and DNM annotation. (**Top**) DNM rates between male and female probands were tested using a 2-sided exact Poisson test. (**Bottom**) DNM rates in male and female probands compared to the DNM model (dotted lines) under a Poisson model. All *p*-values are two-sided, and “All DNM” encompasses synonymous, PTV, and missense DNMs.

**Supplementary Figure 9: Participant sex on X chromosome burden**

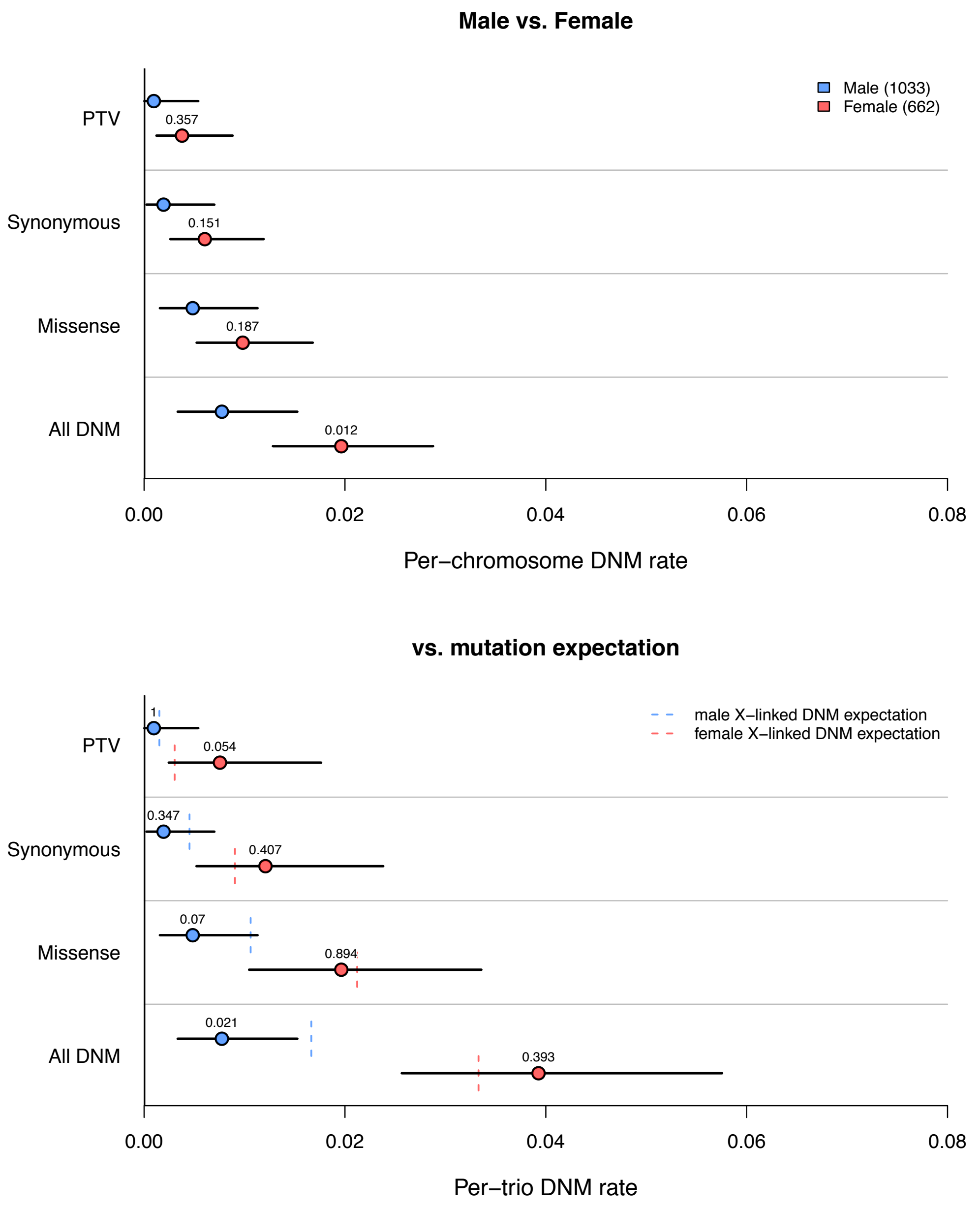


**Figure legend:** X-linked DNM rate in Taiwanese trios split by proband sex and DNM annotation. (**Top**) Per-chromosome DNM rates between male and female probands tested using a 2-sided exact Poisson test, with males inheriting one X chromosome and females inheriting two X chromosomes. (**Bottom**) Per-trio DNM rates in male and female probands compared to the mutation expectation (dotted lines) under a Poisson model. All *p*-values are two-sided, and "All DNM" encompasses synonymous, PTV, and missense DNMs.

**Supplementary Figure 10: Family history of mental illness on DNM burden**

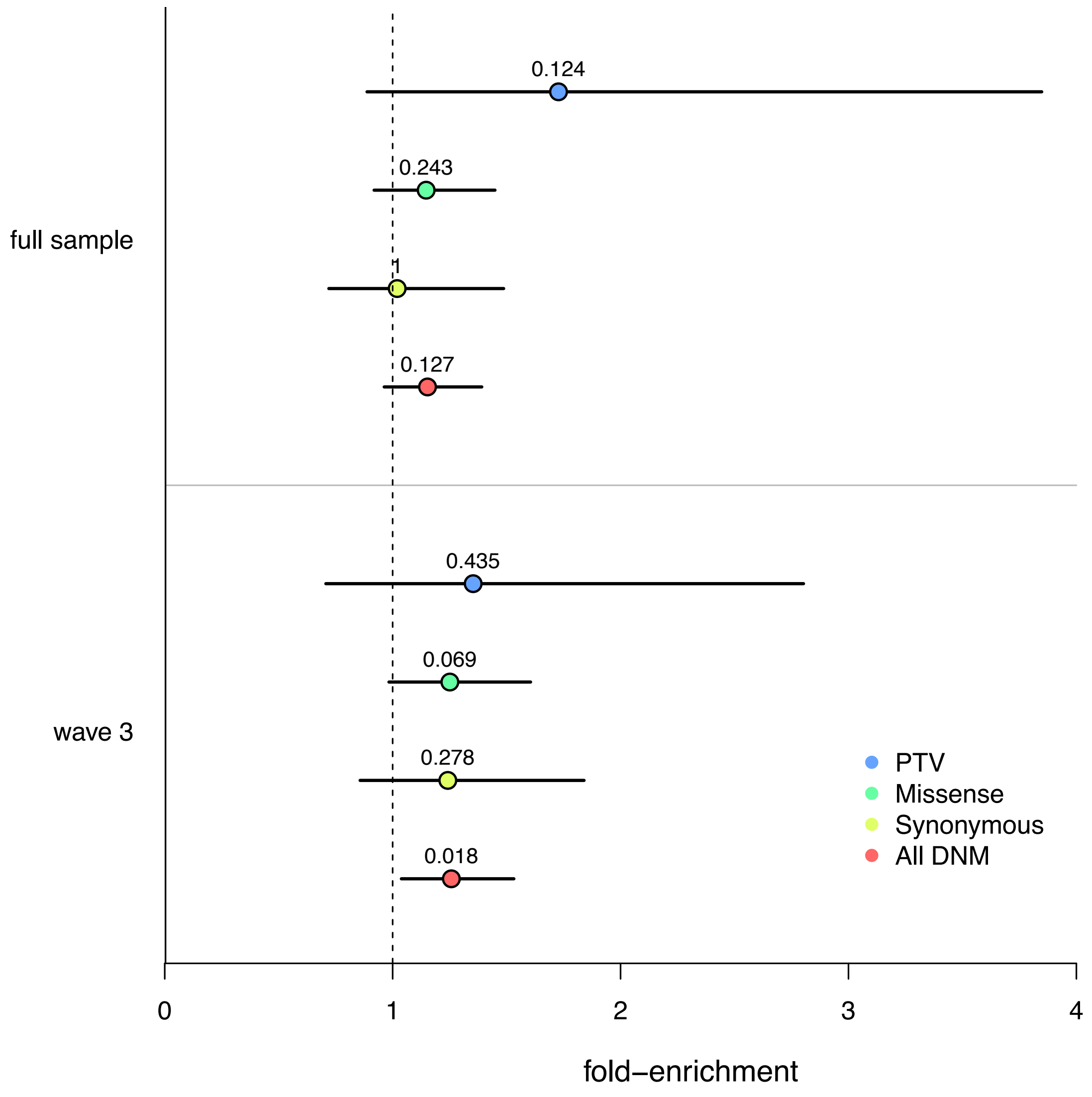


**Figure legend:** Effect of family history of mental illness on DNM burden among Taiwanese trio cohorts. Fold-enrichment represents the DNM burden of SCZ probands with no family history of mental illness over SCZ probands with a family history of mental illness. Results pictured are from a Poisson regression model with no additional covariates.

**Supplementary Figure 11: DNM burden on CPT scores of sustained attention**

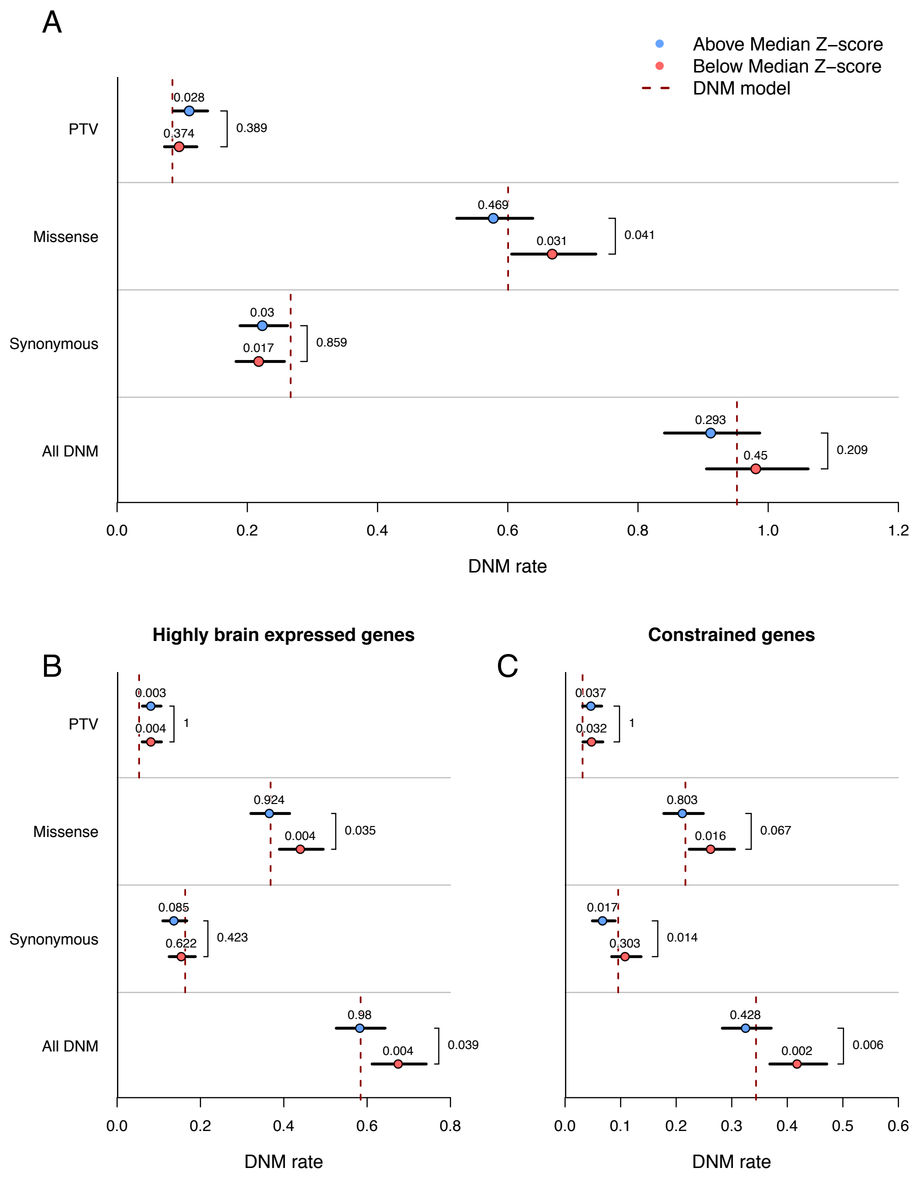


**Figure legend:** Proband DNM rates between high (blue) and low (red) Z-scores in the Continuous Performance Test (CPT) using a median split. Figure A) Exome-wide DNM rates split by primary annotation, where "All DNM" encompasses synonymous, PTV, and missense DNMs. Poisson rate *p*-values compared to the DNM model expectation (dotted lines) are directly above the rate estimate in each group, and two-sample, two-sided Poisson exact test *p*-values between high and low scorers are listed to the right of the rate estimates. DNM rates restricted to enriched gene sets among SCZ probands: 10,376 highly brain expressed genes (Figure B) and 4,083 constrained genes (Figure C).

#### Supplementary Figure 12: DNM burden on WCST scores of executive function

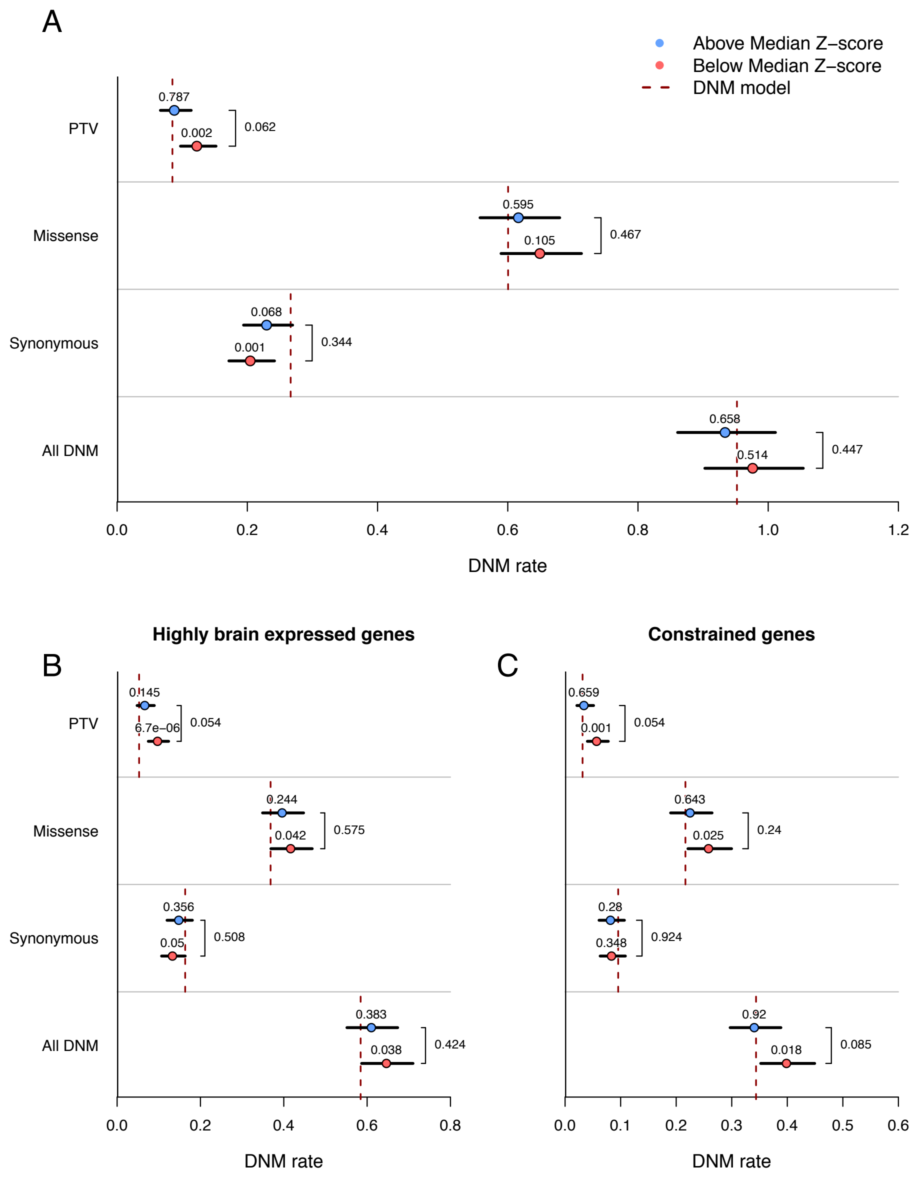


**Figure legend:** Proband DNM rates between high (blue) and low (red) Z-scores in the Wisconsin Card Sorting Task (WCST) using a median split. Figure A) Exome-wide DNM rates split by primary annotation, where “All DNM” encompasses synonymous, PTV, and missense DNMs. Poisson rate *p*-values compared to the DNM model expectation (dotted lines) are directly above the rate estimate in each group, and two-sample, two-sided Poisson exact test *p*-values between high and low scorers are listed to the right of the rate estimates. DNM rates restricted to enriched gene sets among SCZ probands: 10,376 highly brain expressed genes (Figure B) and 4,083 constrained genes (Figure C).

**Supplementary Figure 13: Synonymous DNM rate by study**

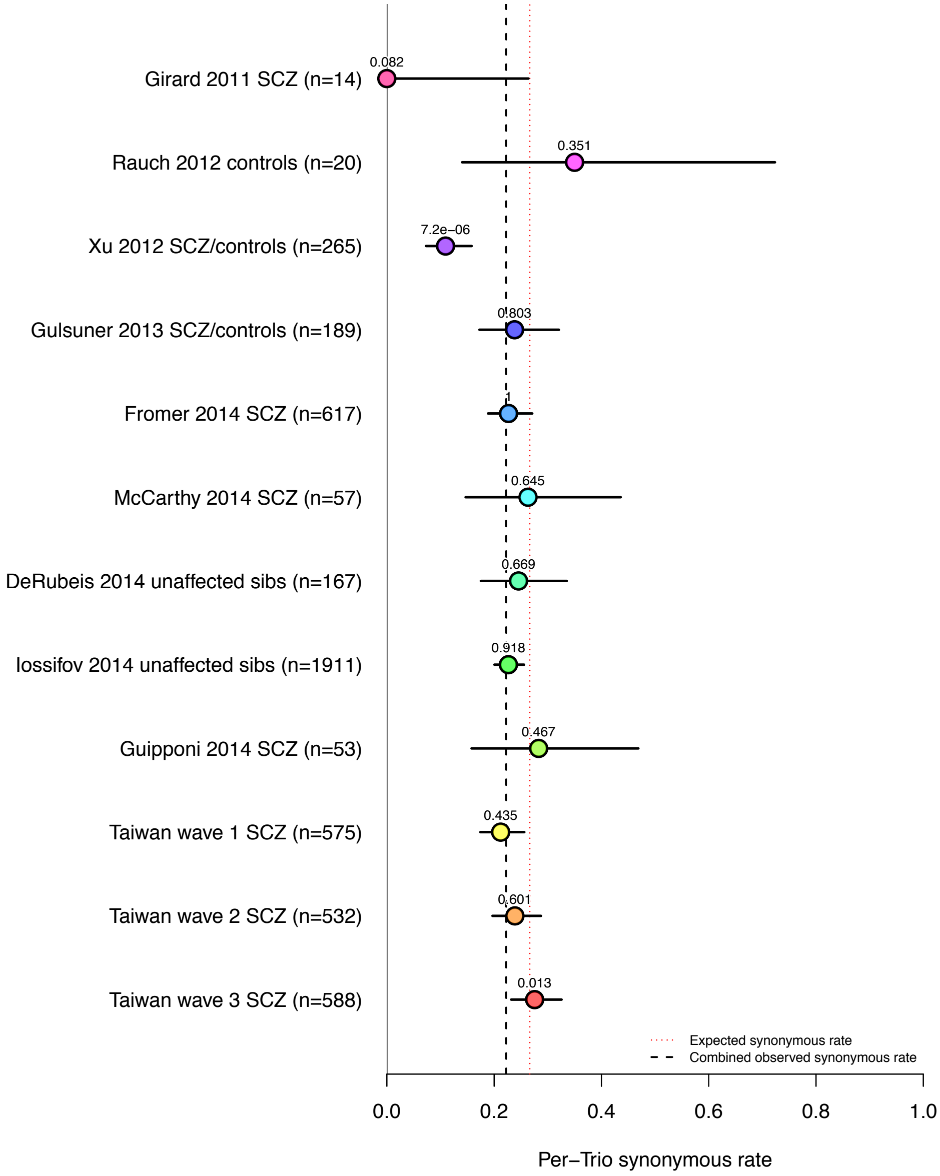


**Figure legend:** Per-trio synonymous DNM rate by study. DNMs were restricted to validated and published DNMs falling within hg19 Agilent v2 target intervals. Poisson two-sample test *p*-values compare the study synonymous DNM rate to the synonymous DNM rate of all other studies combined.

#### Supplementary Figure 14: Taiwanese vs. published SCZ DNM rate

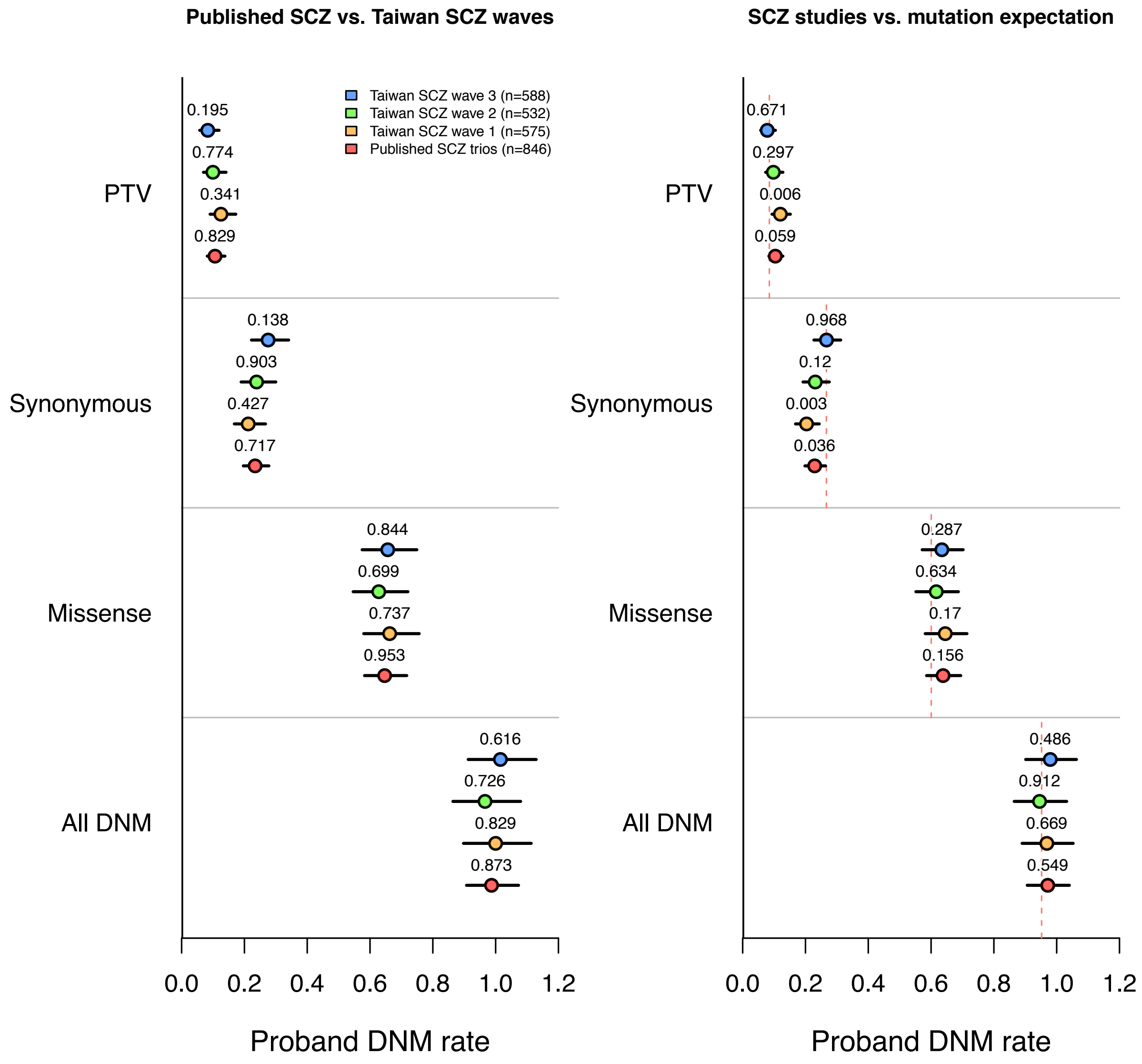


**Figure legend**: DNM rate in published SCZ and Taiwanese trios split by DNM annotation. (**Left**) Per-trio DNM rates between cohorts tested using a 2-sided exact Poisson test, with each cohort being compared to the other three cohorts combined. (**Right**) Per-trio DNM rates in each cohort compared to the mutation expectation (dotted lines) under a Poisson model. All *p*-values are two-sided, and "All DNM" encompasses synonymous, PTV, and missense DNMs.

**Supplementary Figure 15: Missense predictor enrichment**

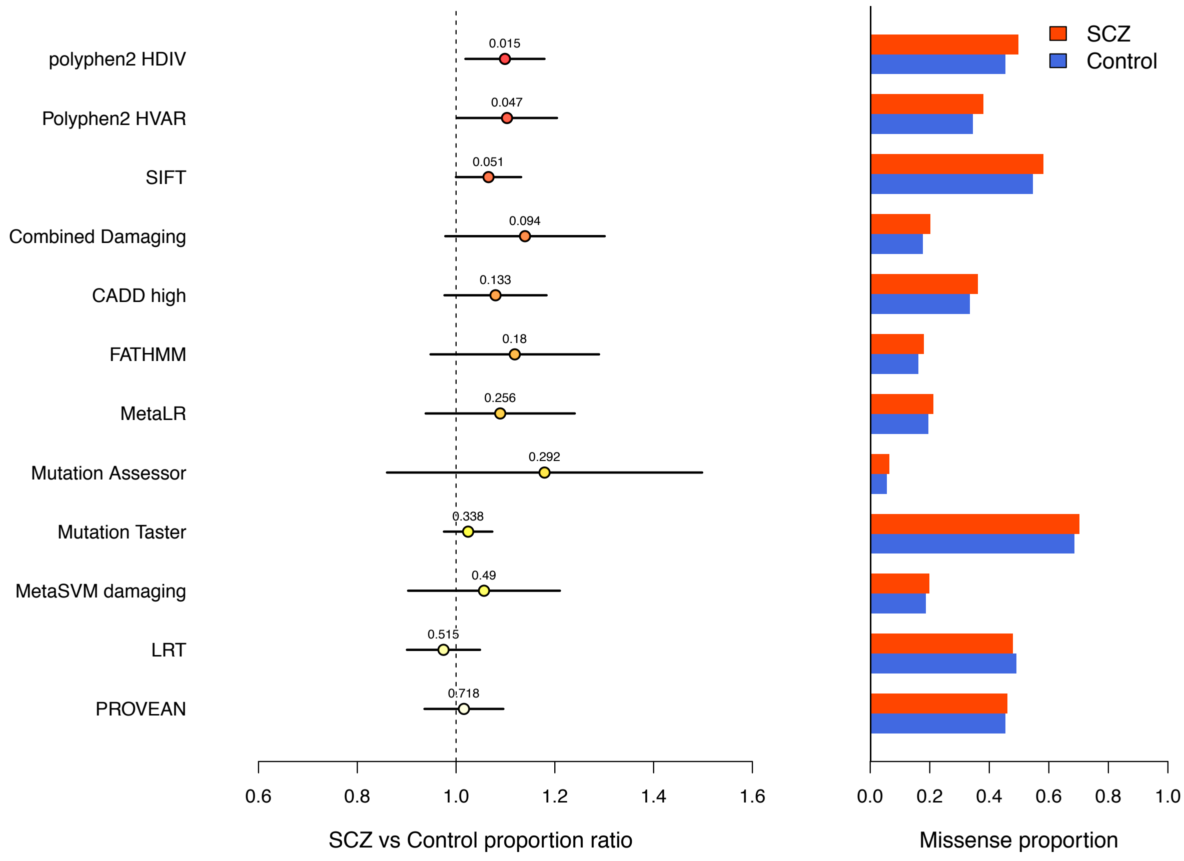


**Figure legend:** Proportion of predicted damaging missense DNMs relative to overall missense DNM rate in SCZ probands and controls. Prediction algorithms are sorted by the two-sample proportion test *p*-value.

**Supplementary Figure 16: Non-ExAC missense predictor enrichment**

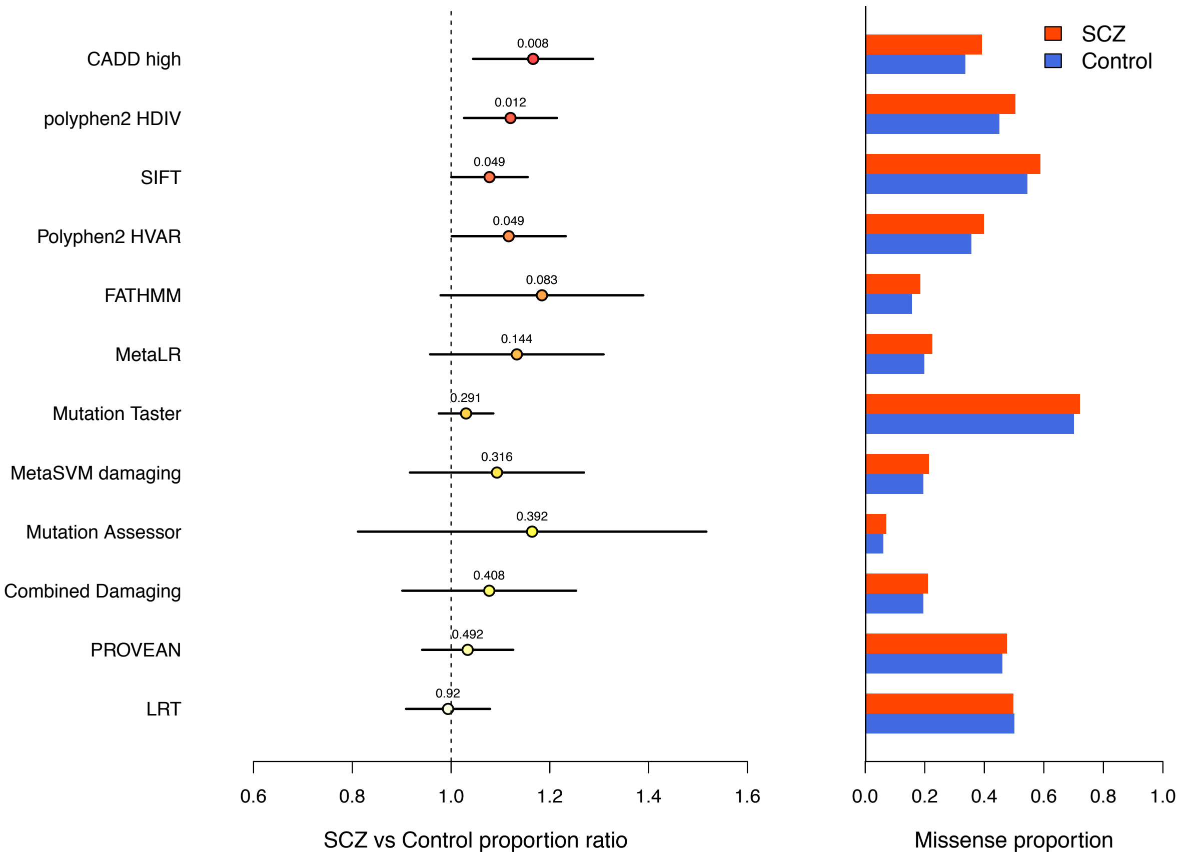


**Figure legend:** Among DNMs not seen in the non-Psych ExAC cohort, the proportion of predicted damaging missense DNMs relative to overall missense DNM rate in SCZ probands and controls. Prediction algorithms are sorted by the two-sample proportion test *p*-value.

**Supplementary Figure 17: Synonymous DNM enrichment at NSS-ESR sites**

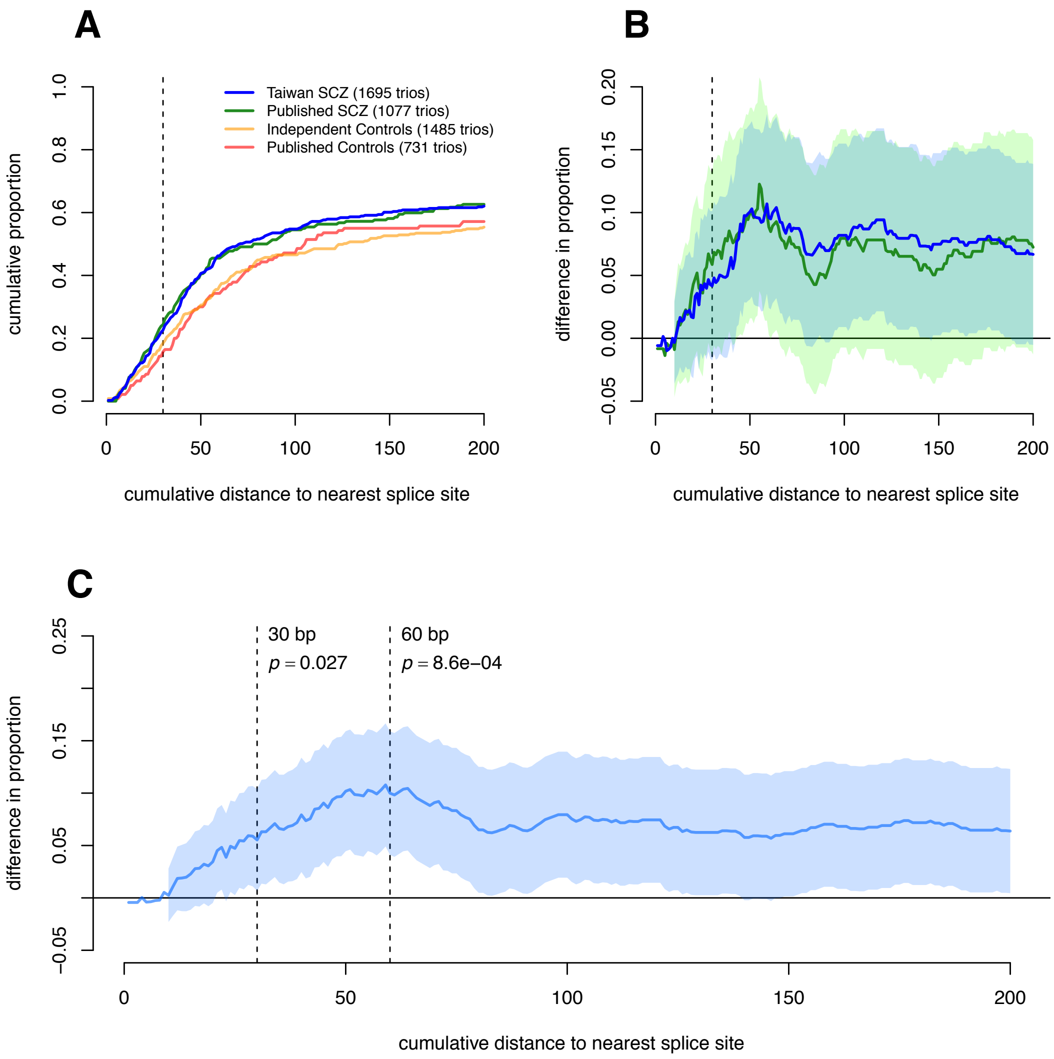


**Figure legend:** Among synonymous sites within exon-splice regulator (ESR) motifs, the enrichment between SCZ probands and controls as a function of distance to nearest splice site. **Figure A:** Cumulative proportion of ESR synonymous sites as distance to nearest splice site increases. Cohorts are split by previously published and analyzed SCZ probands and controls (Published), and the Taiwanese trio cohort and published controls not previously analyzed (Independent). Dotted line 30bp from splice site is the suggested definition of a “near splice-site” mutation. **Figure B:** Difference in proportion (and shaded 95% CI) between Taiwanese cohort (blue) and published SCZ cohorts (green) and the combined controls. **Figure C:** Difference in proportion (and shaded 95% CI) for combined SCZ and control cohort. Vertical dotted lines show the suggested (30bp) and best-perfoming (60 bp) “near-splice site” definition.

**Supplementary Figure 18: Synonymous annotation enrichment**

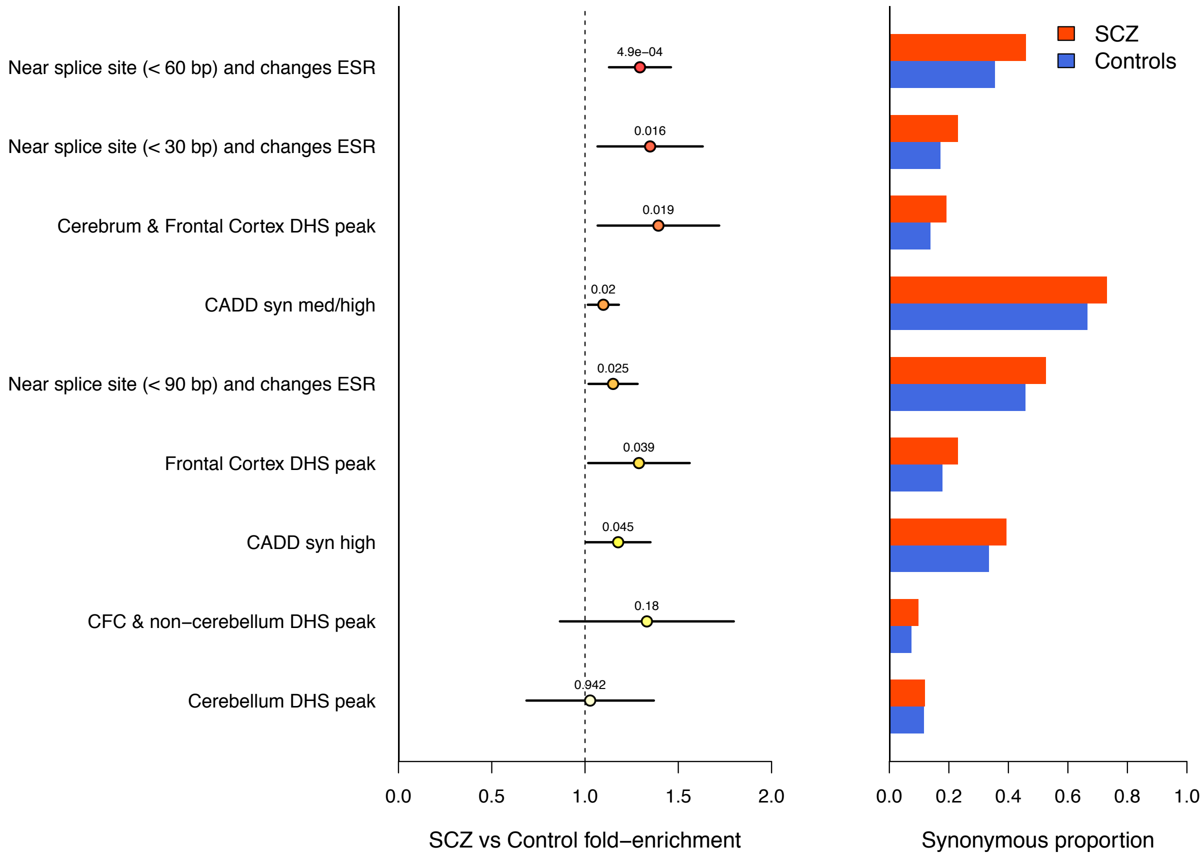


**Figure legend:** Proportion of "functional" annotations in synonymous DNMs relative to overall synonymous DNM rate in SCZ probands and controls. Functional annotations are sorted by the two-sample proportion test *p*-value.

#### Supplementary Figure 19: SCZ enriched annotation

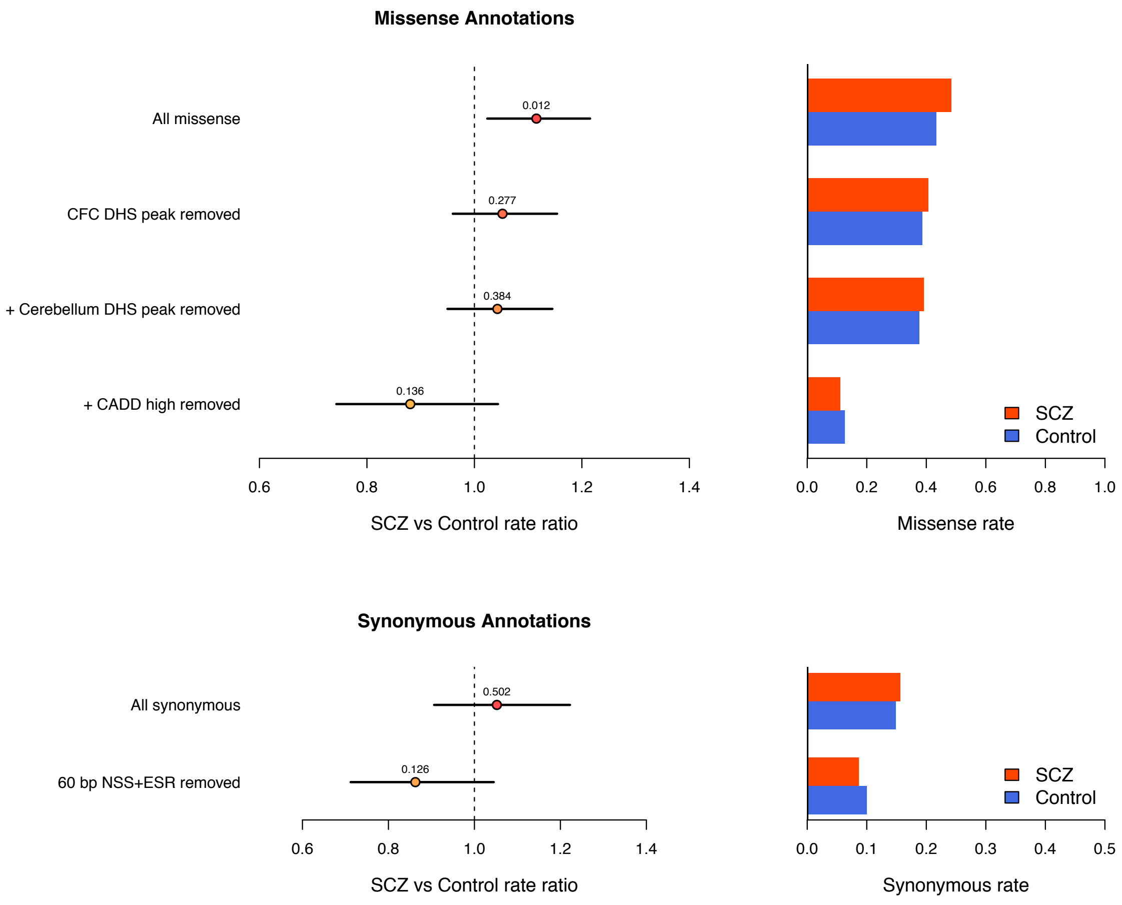


**Figure legend:** Selection of secondary annotations enriched ($p < 0.05$) in SCZ probands relative to controls. DNMs are restricted to those not seen in the non-Psych ExAC cohort, and annotation selection is sorted by their proportion test *p*-value (lowest *p*-value first). Annotations are applied in a step-wise fashion until the DNM rate in controls is greater than that seen in SCZ probands. Among PTV DNMs, no annotation has a *p*-value < 0.05, so no annotations were applied. Results are presented as both rate ratios (left graphs) and overall rates (right graphs).

**Supplementary figure 20: FWER passing gene set enrichment**

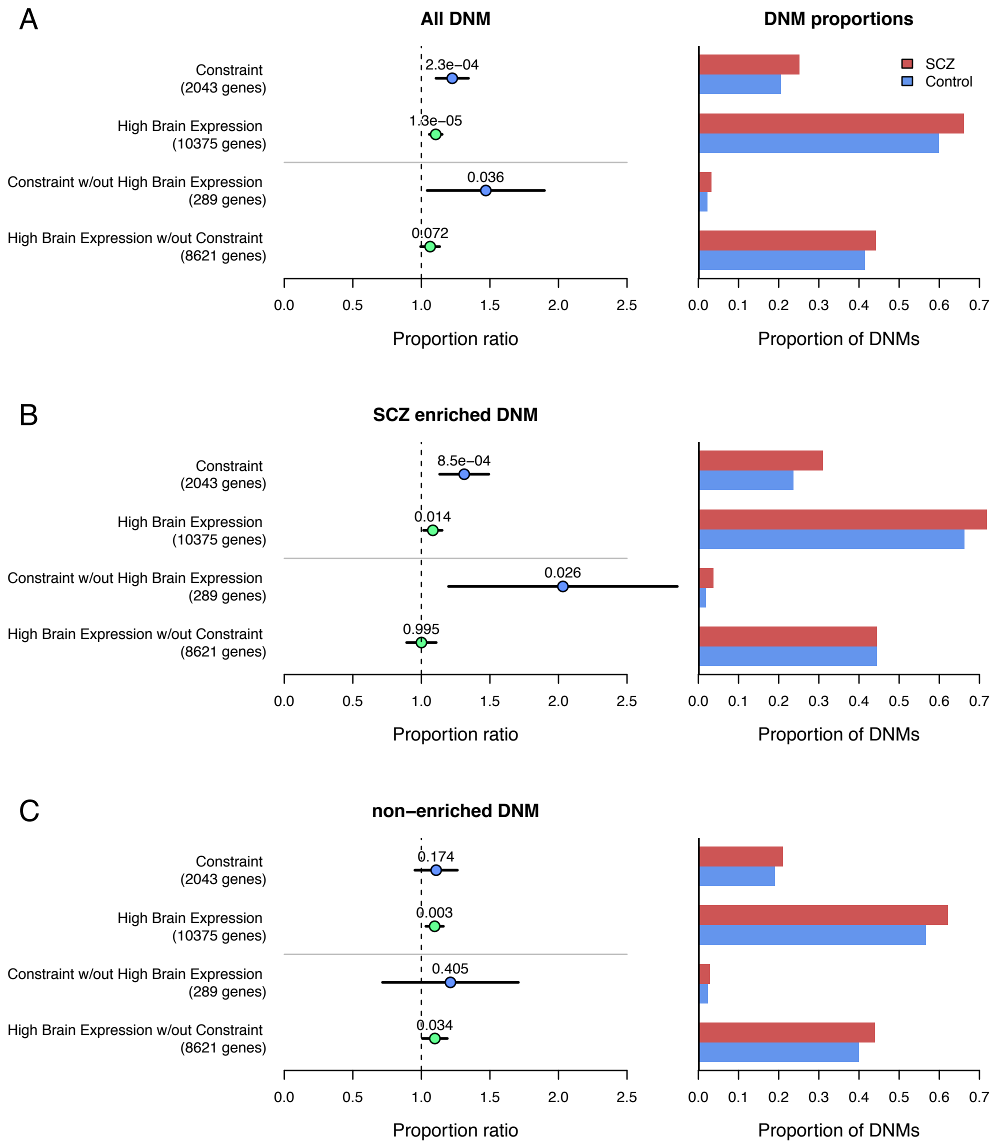


**Figure legend:** Gene set enrichment among the principal gene sets that surpassed multiple testing correction in both the mutation model and relative to controls. "Constraint" is the inclusive combination of missense constraint and RVIS intolerant gene sets, and "High Brain Expression" is the inclusive combination of BrainSpan high brain expression and GTEx brain enriched gene sets. **Figure A:** Gene set enrichment in SCZ probands compared to controls among all DNMs, with proportion ratios and 95% CI (left graph) as well as overall proportion of total DNMs. DNM lists were further split by SCZ-enriched DNMs (**Figure B**) and non-enriched DNMs (**Figure C**).

**Supplementary figure 21: Evolutionary constraint gene set enrichment**

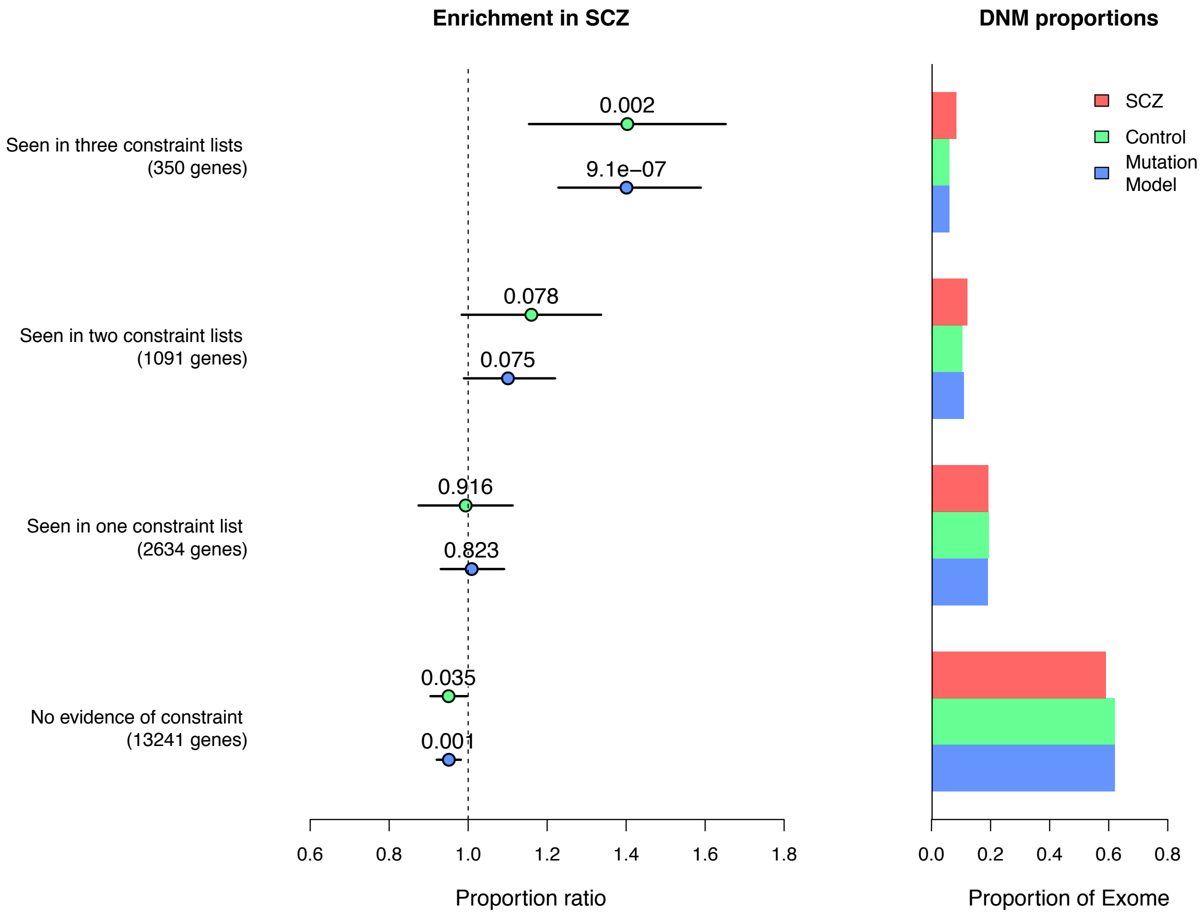


**Figure legend**: Gene set enrichment among all DNMs within the overlap of high pLI ( > 0.9; 3431 genes), missense constrained (1587 genes), and RVIS intolerant (859 genes) gene sets. Results are presented as the proportion ratio (**left graph**) relative to controls (green) and the mutation model (blue), and overall proportions (**right graph**).

**Supplementary figure 22: Evolutionary constraint gene set PTV enrichment**

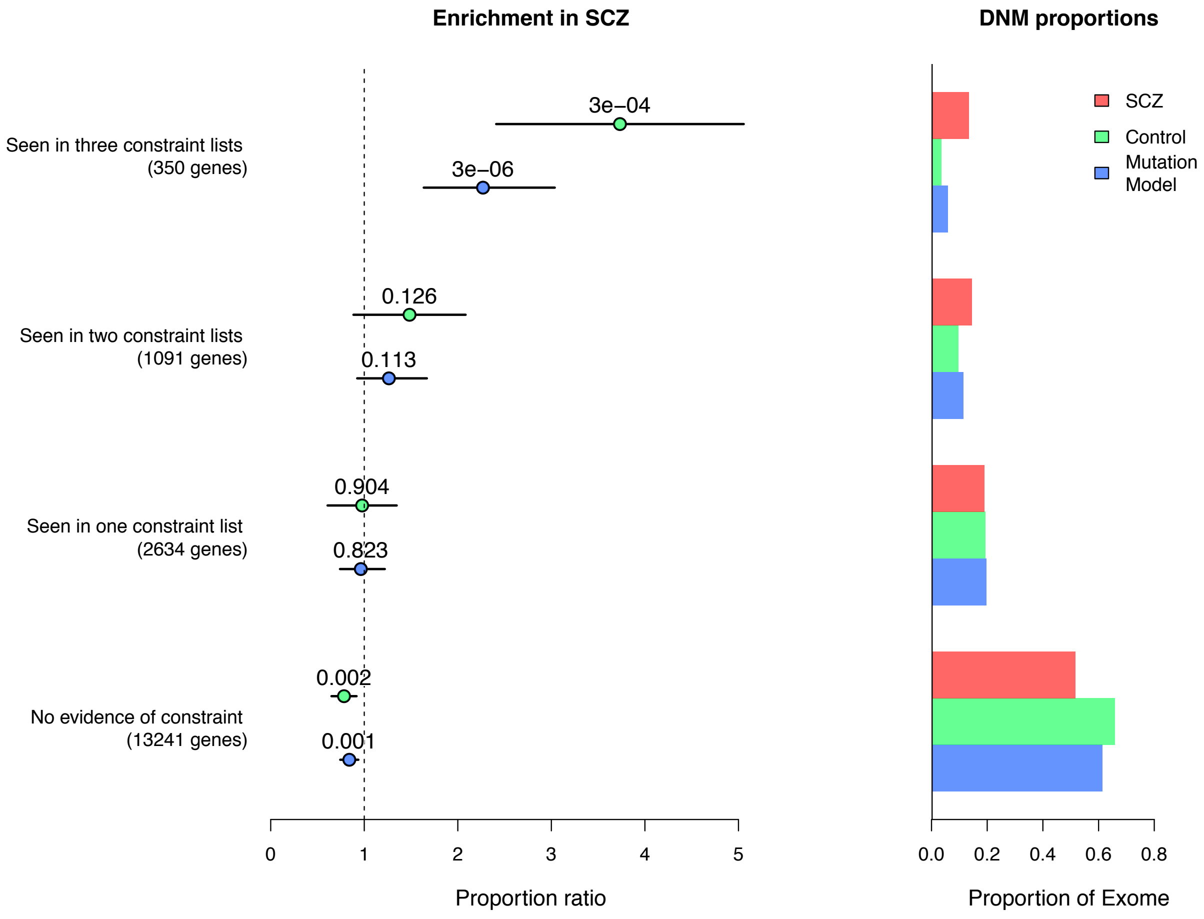


**Figure legend**: Gene set enrichment among PTV DNMs within the overlap of high pLI ( > 0.9; 3431 genes), missense constrained (1587 genes), and RVIS intolerant (859 genes) gene sets. Results are presented as the proportion ratio (**left graph**) relative to controls (green) and the mutation model (blue), and overall proportions (**right graph**).

### Supplementary figure 23: Brain expression gene set enrichment

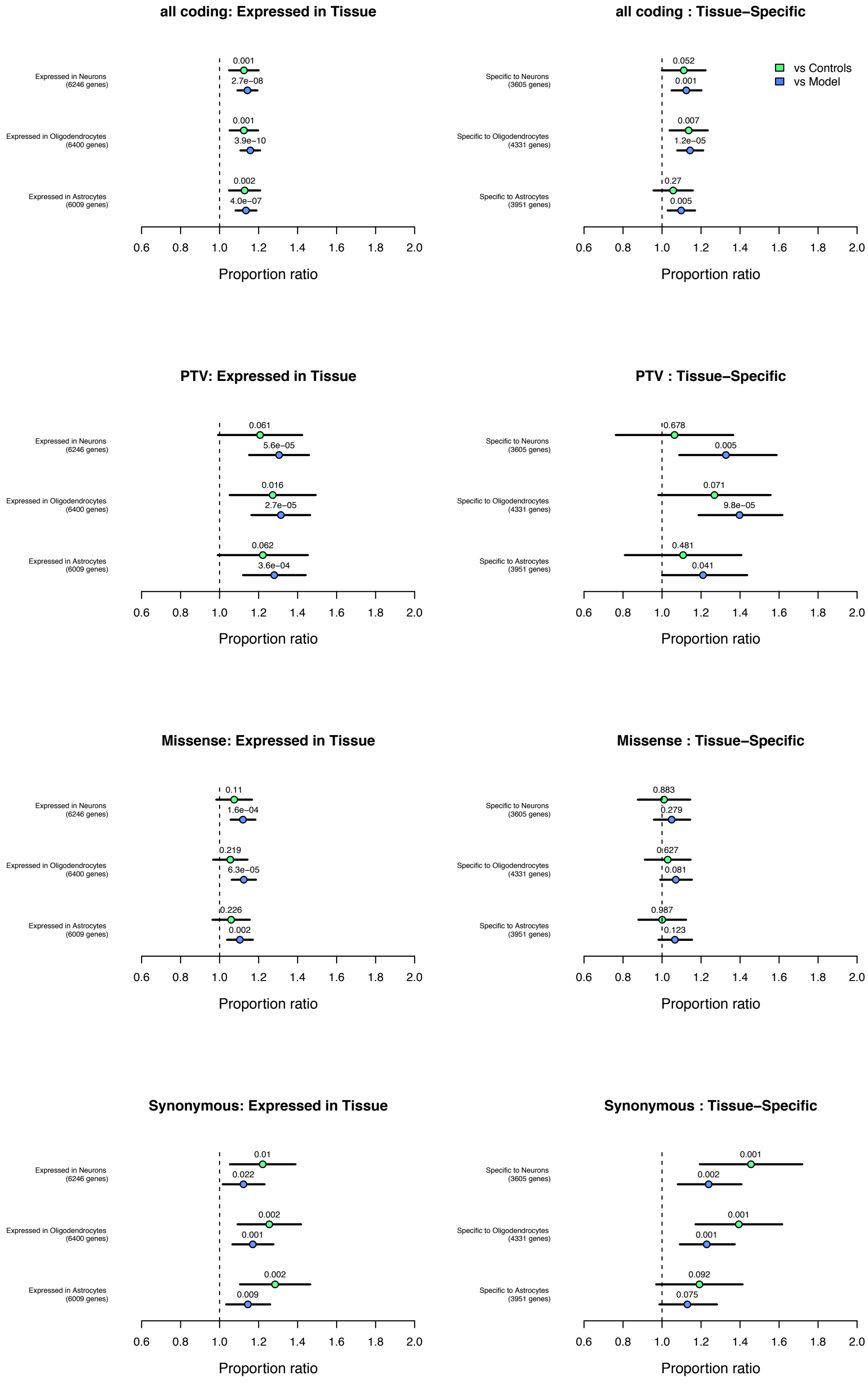

**Figure legend:** Partitioning of high brain expression enrichment signal to specific brain cell tissue types. Brain cell tissue type expression is split by high expression (**left graphs**) and specific expression (**right graphs**). DNMs are split by annotation (**top to bottom**) and tested against controls (**green**) and the mutation model (**blue**).

#### Supplementary figure 24: Neurodevelopmental disease gene set enrichment

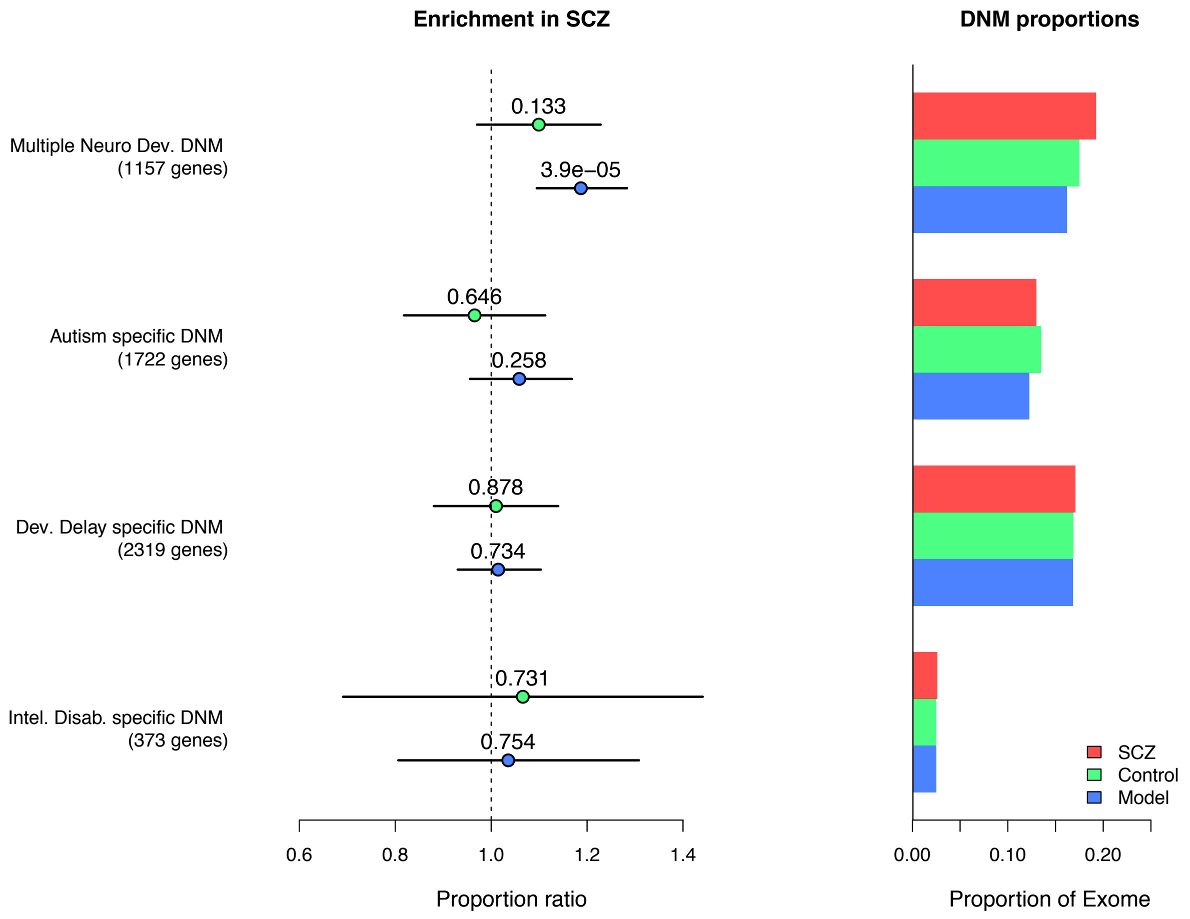


**Figure legend**: Gene set enrichment among all DNMs in SCZ probands within the overlap of genes implicated in Autism, Developmental Delay, and Intellectual Disability trio exome studies. Results are presented as the proportion ratio (**left graph**) relative to controls (green) and the mutation model (blue), and overall proportions (**right graph**).

**Supplementary figure 25: Synaptic enrichment by exome coverage**

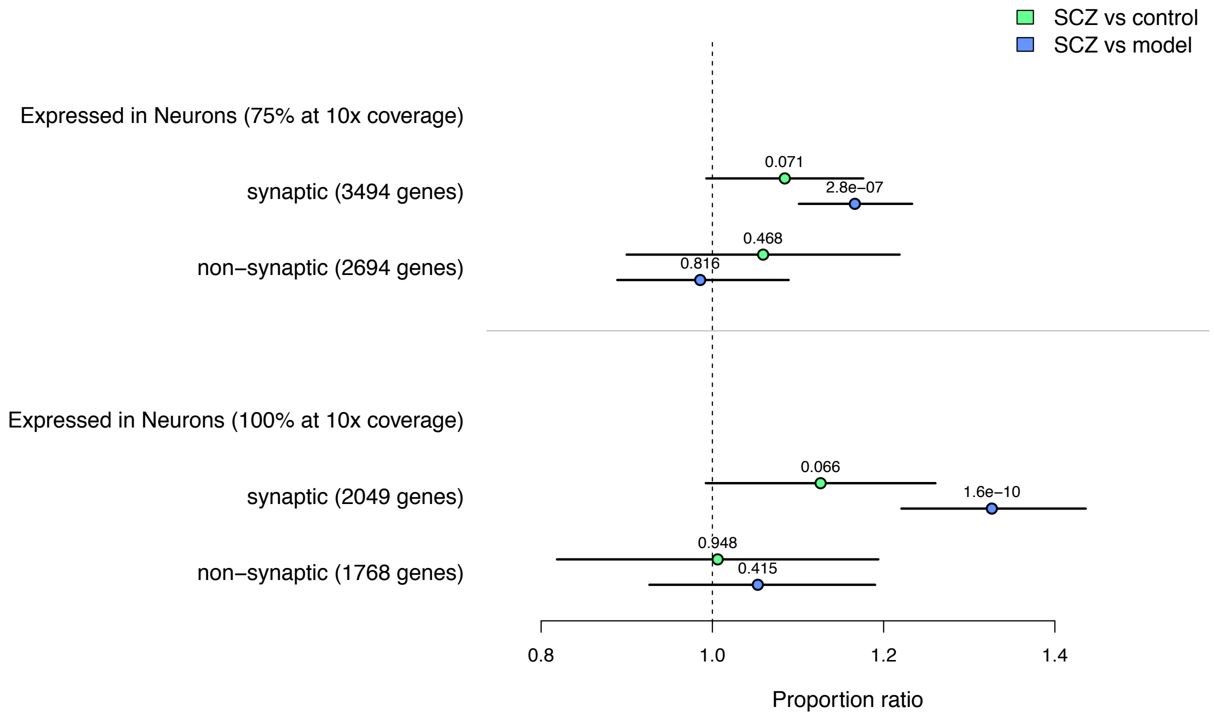


**Figure legend**: Gene set enrichment for all DNMs in SCZ probands within potentially synaptic genes. Results are presented as the proportion ratio relative to controls (green) and the mutation model (blue). The top graph represents 17,925 genes where at least 75% of their coding sequence is covered at 10x. The bottom graph represents 12,311 genes where all 100% of their coding sequence covered at 10x.

**Supplementary figure 26: Neurodevelopmental disease gene set enrichment by exome coverage**

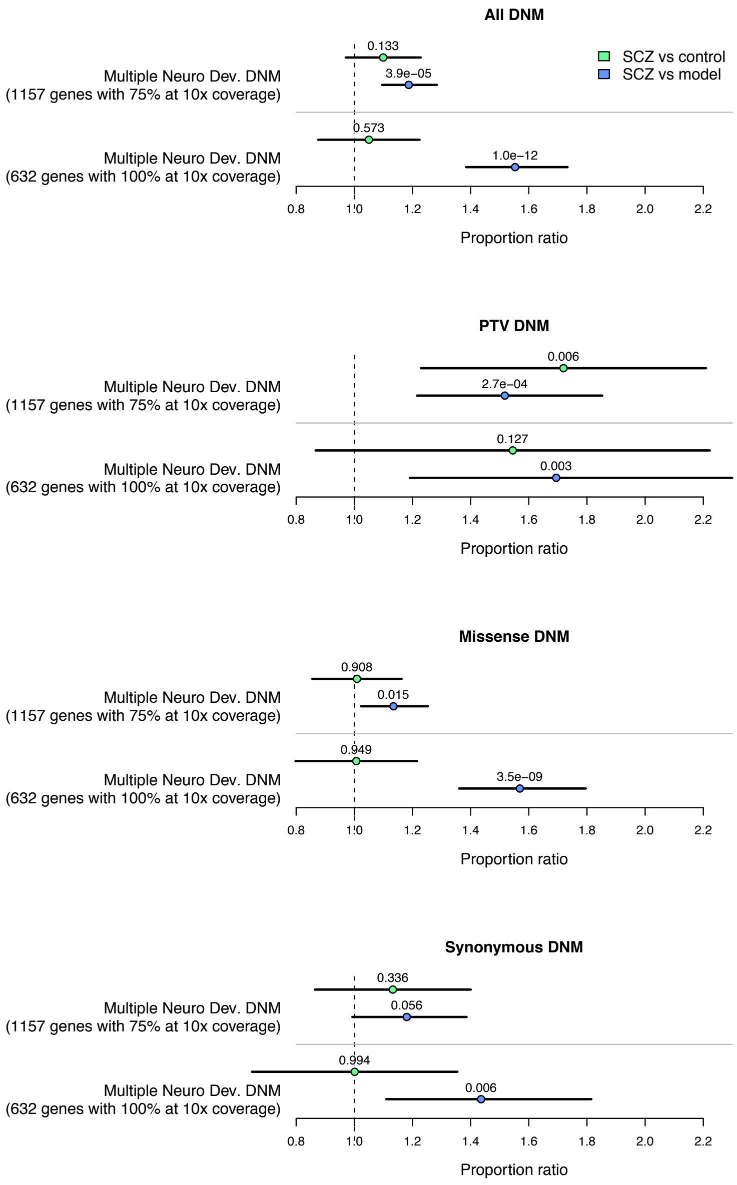


**Figure legend**: Gene set enrichment among various DNM annotations in SCZ probands within the overlap of genes implicated in Autism, Developmental Delay, and Intellectual Disability trio exome studies. Results are presented as the proportion ratio relative to controls (green) and the mutation model (blue). With each annotation, the top comparison represents 17,925 genes where at least 75% of their coding sequence is covered at 10x, and the bottom comparison represents 12,311 genes where all 100% of their coding sequence covered at 10x.

**Supplementary figure 27: GO/SynaptomeDB gene set model enrichment**

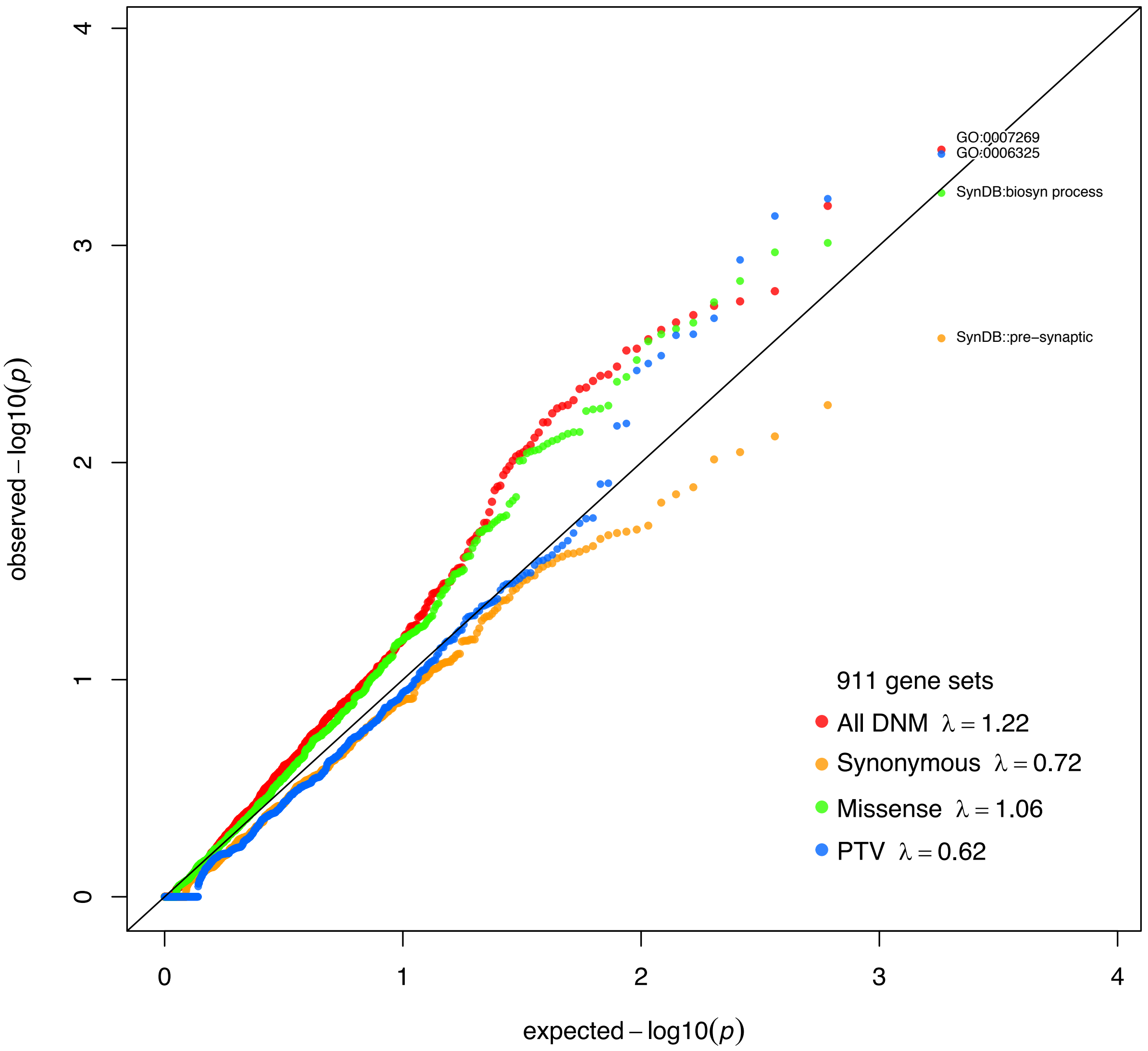


**Figure legend:** QQ plot of gene set enrichment among all DNMs in SCZ probands compared to mutation model expectations in 911 qualifying GO and SynaptomeDB gene sets.

**Supplementary figure 28: GO/SynaptomeDB gene set case-control enrichment**

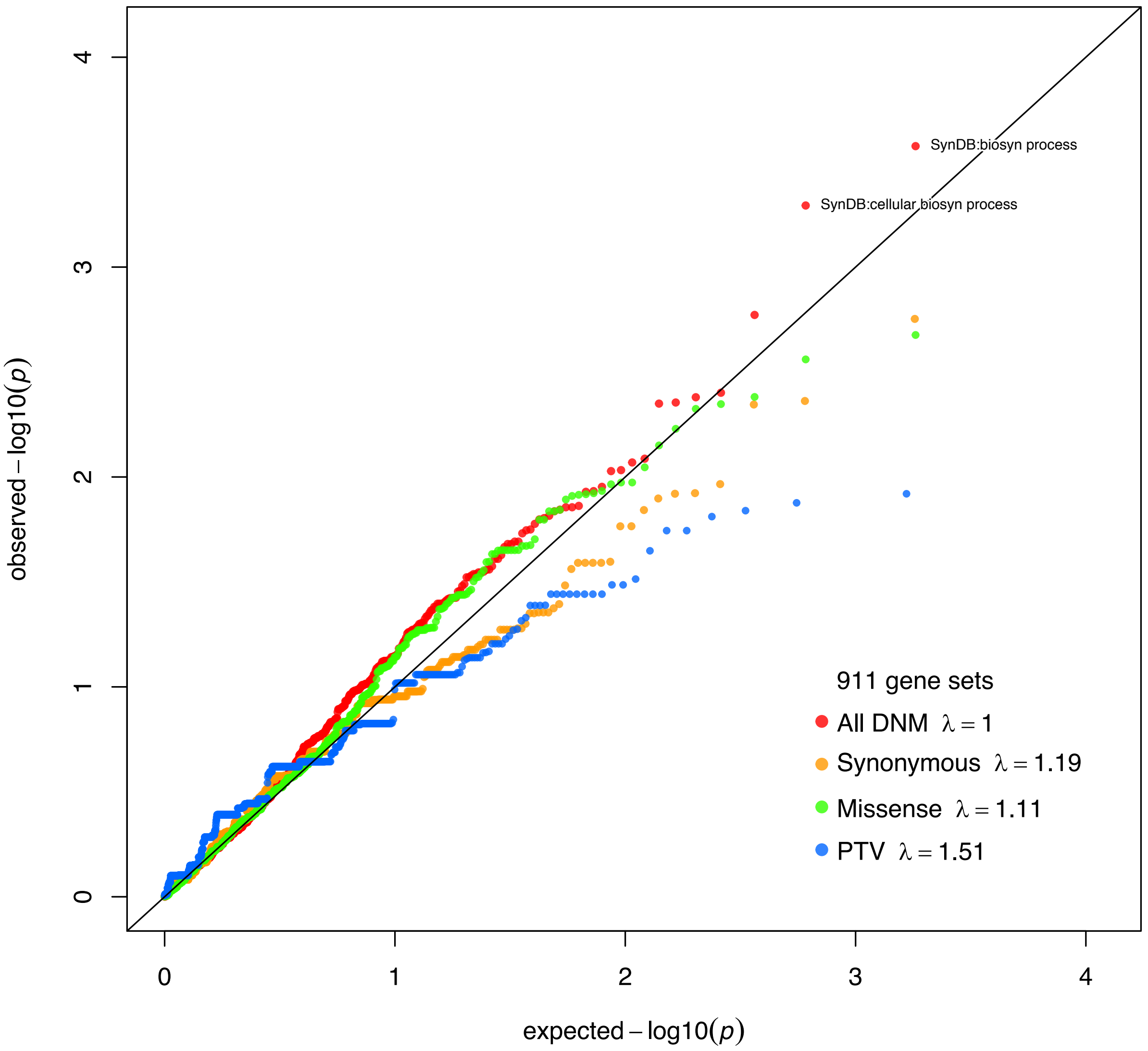


**Figure legend:** QQ plot of gene set enrichment among all DNMs in SCZ probands compared to controls in 911 qualifying GO and SynaptomeDB gene sets.

#### Supplementary Figure 29: CPT scores of recurrent PTV gene carriers in Taiwanese cohort

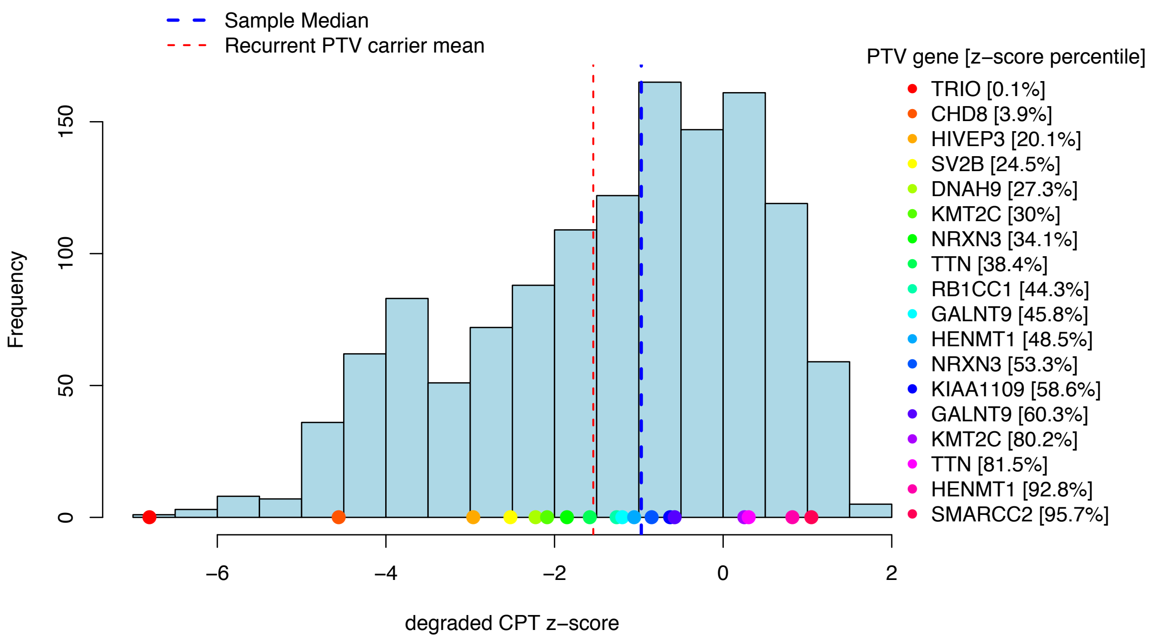


**Figure legend:** Distribution of z-scores from the Continuous Performance Test (CPT) available from 1306 probands in the Taiwanese cohort. Lower z-scores indicate poorer performance. Points plotted on the x-axis represent z-scores of probands carrying a PTV DNM in one of 16 recurrently hit genes among the full SCZ cohort. The right-hand legend also lists the gene symbol and z-score percentile of the proband PTV carrier.

**Supplementary Figure 30: WCST scores of recurrent PTV gene carriers in Taiwanese cohort**

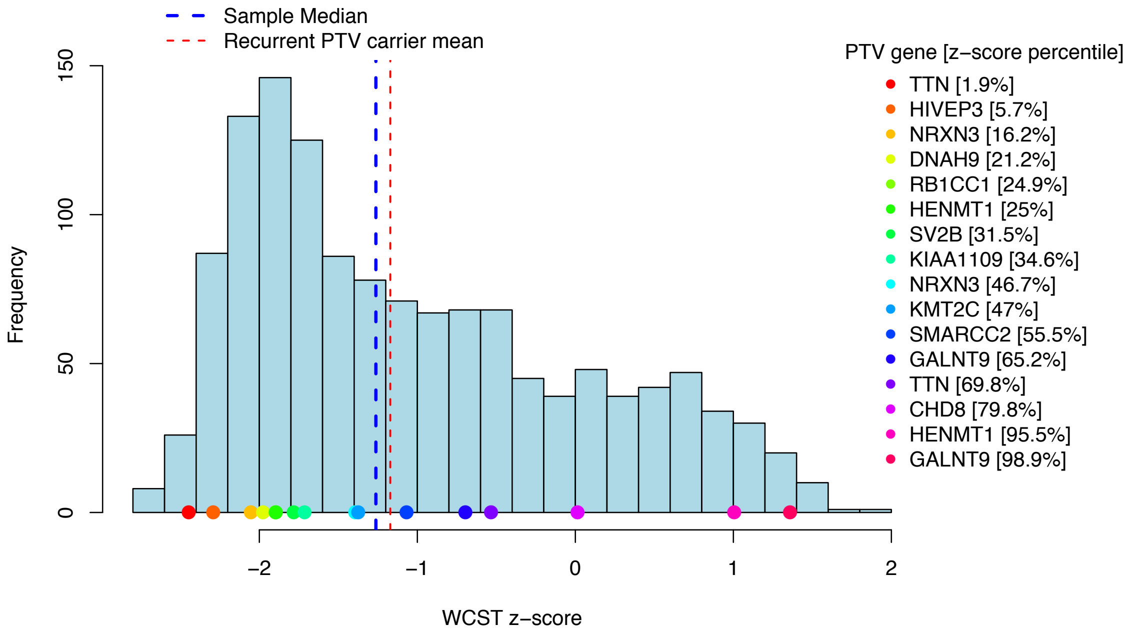


**Figure legend:** Distribution of z-scores from the Wisconsin Card sorting Task (WCST) available from 1325 probands in the Taiwanese cohort. Lower z-scores indicate poorer performance. Points plotted on the x-axis represent z-scores of probands carrying a PTV DNM in one of 16 recurrently hit genes among the full SCZ cohort. The right-hand legend also lists the gene symbol and z-score percentile of the proband PTV carrier.

**Supplementary Figure 31: Synonymous transmission by variant quality score**

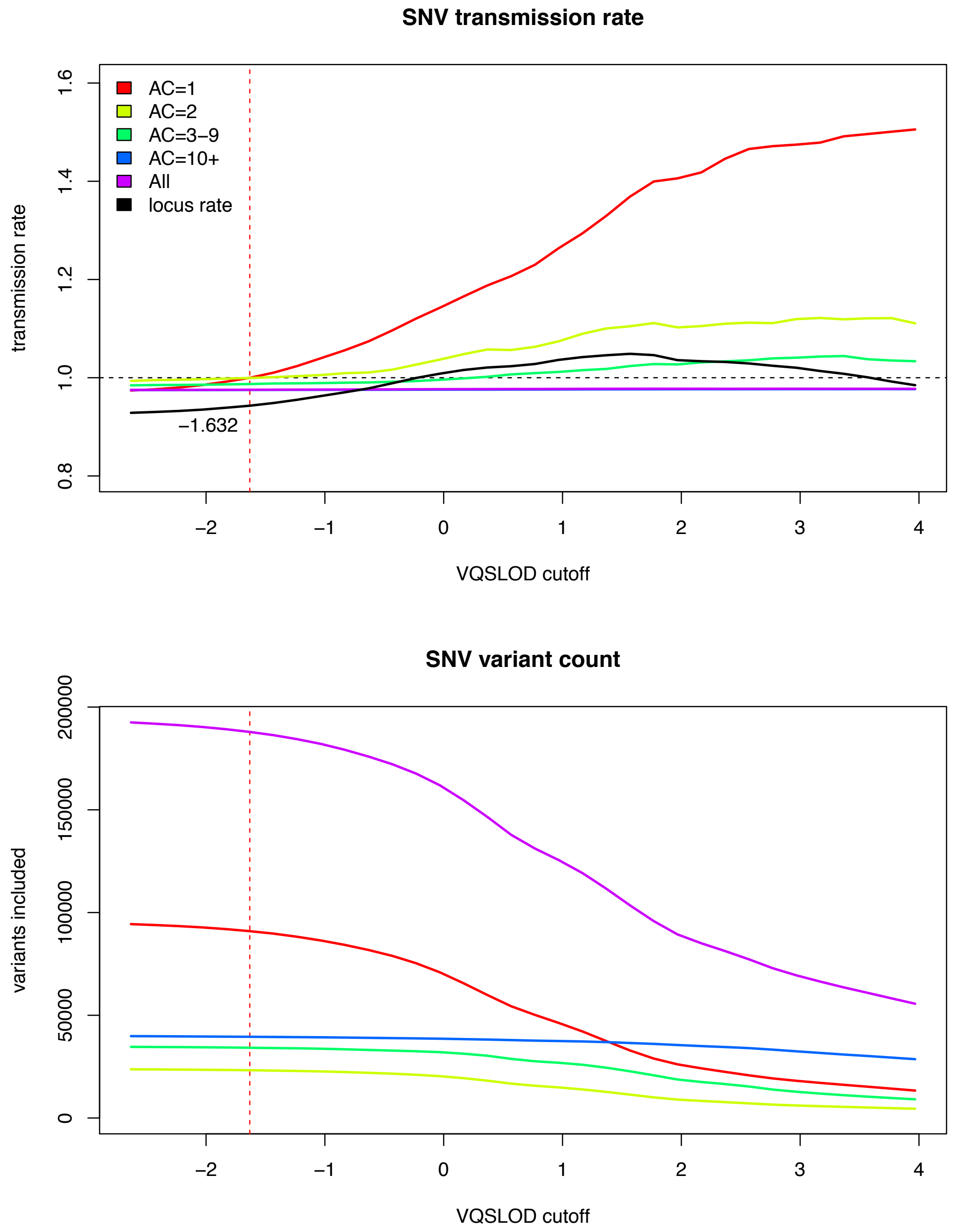


**Figure legend:** Adjustment of variant quality inclusion to ensure a balanced transmission rate among synonymous singletons in the parents of the Taiwanese cohort. **Top graph**: Variants are included if they have a VQSLOD score equal to or greater than the cutoff specified (x-axis), with the transmission rate being the sum of transmitted variants divided by the sum on non-transmitted variants. **Bottom graph**: the number of variants passing the VQSLOD cutoff.

**Supplementary Figure 32: Gene set model comparison with SCZ case-control exomes**

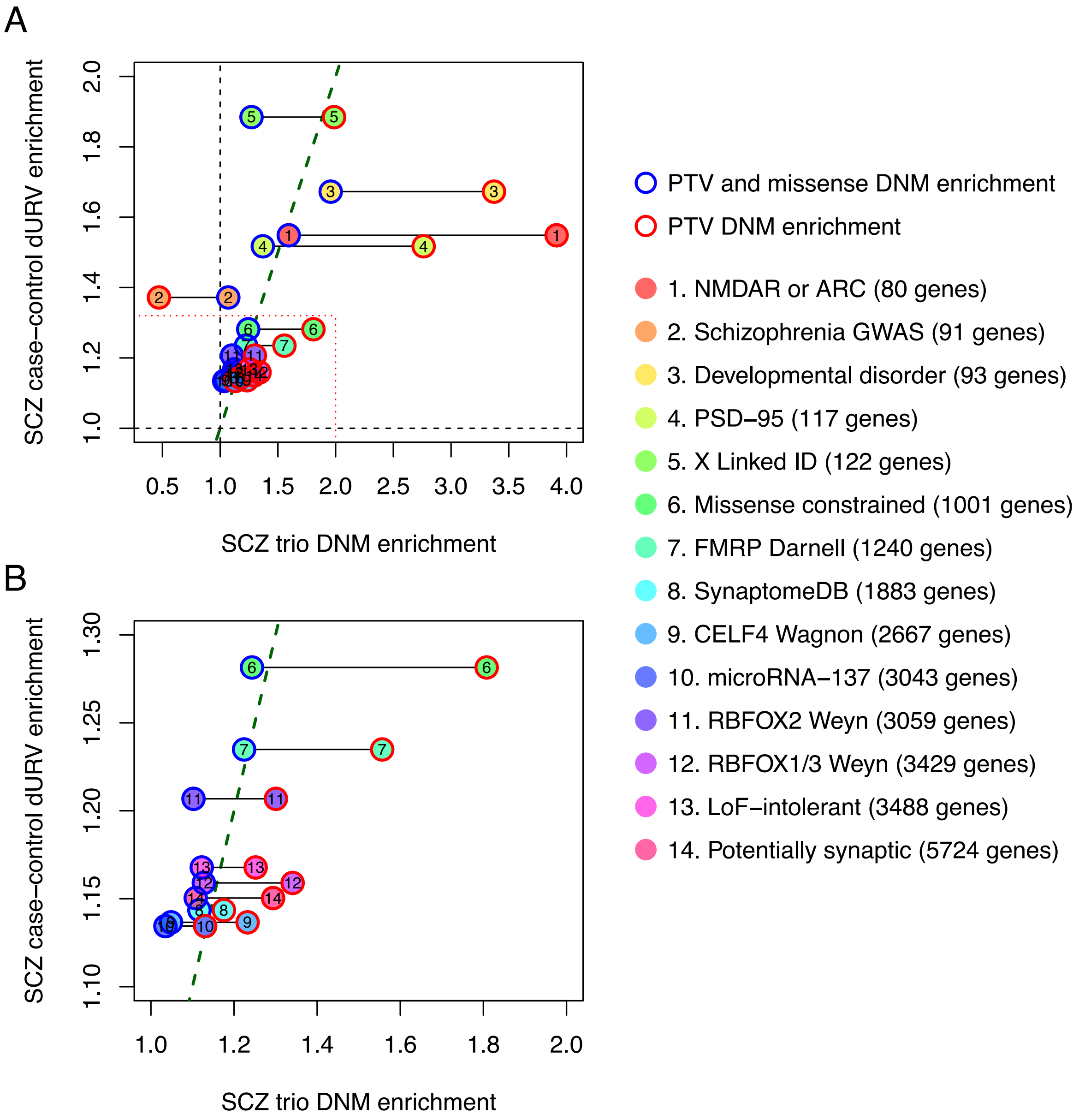


**Figure legend:** Comparison of effect size estimates in fourteen gene sets tested in both SCZ case-control exomes and mutation model gene set enrichment in SCZ trio exomes. **Figure B** is simply a closer view of the gene sets within the red-dotted lines of **figure A**. **X-axis:** SCZ trio DNM enrichment is the proportion enrichment of PTV and missense DNMs (blue circle) and PTV DNMs (red circle) relative to the mutation model, with black lines indicating the difference between these two enrichment tests. **Y-axis:** SCZ case-control dURV (damaging or disruptive ultra-rare variant) enrichment is the odds ratio of SCZ cases carrying a dURV in the gene set relative to controls. The green dotted line has a slope of 1, where points to the left of the line mean that they are more enriched in SCZ case-control exomes than in SCZ trio exomes, and points to the right of the line are more enriched in SCZ trio exomes than in SCZ case-control exomes. Gene counts listed in the legend match the SCZ case-control study (Genovese et al., 2016), and differences in gene counts in the SCZ trio enrichment are usually due to restricted to well-covered genes.

**Supplementary Figure 33: Gene set case-control comparison with SCZ case-control exomes**

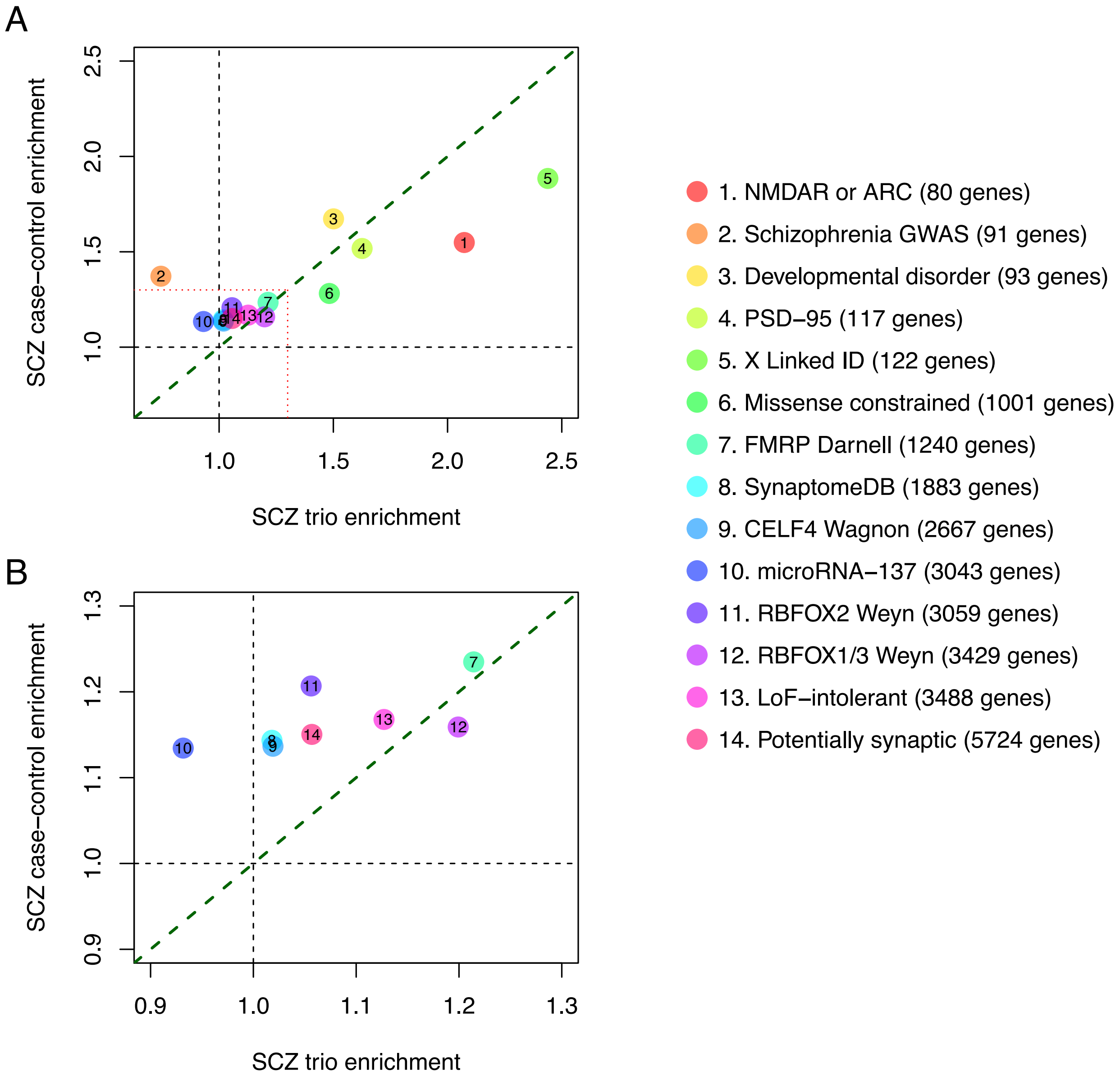


**Figure legend:** Comparison of effect size estimates in fourteen gene sets tested in both SCZ case-control exomes and case-control gene set enrichment in SCZ trio exomes. **Figure B** is simply a closer view of the gene sets within the red-dotted lines of **figure A**. **X-axis:** SCZ trio DNM enrichment is the proportion enrichment of SCZ-enriched DNMs. **Y-axis:** SCZ case-control dURV (damaging or disruptive ultra-rare variant) enrichment is the odds ratio of SCZ cases carrying a dURV in the gene set relative to controls. The green dotted line has a slope of 1, where points to the left of the line mean that they are more enriched in SCZ case-control exomes than in SCZ trio exomes, and points to the right of the line are more enriched in SCZ trio exomes than in SCZ case-control exomes. Gene counts listed in the legend match the SCZ case-control study (Genovese et al., 2016), and differences in gene counts in the SCZ trio enrichment are usually due to restricted to well-covered genes.
